# Human brain anatomy reflects separable genetic and environmental components of socioeconomic status

**DOI:** 10.1101/2021.07.28.454131

**Authors:** Hyeokmoon Kweon, Gökhan Aydogan, Alain Dagher, Danilo Bzdok, Christian C. Ruff, Gideon Nave, Martha J. Farah, Philipp D. Koellinger

## Abstract

Recent studies report that socioeconomic status (SES) correlates with brain structure. Yet, such findings are variable and little is known about underlying causes. We present a well-powered voxel-based analysis of grey matter volume (GMV) across levels of SES, finding many small SES effects widely distributed across the brain, including cortical, subcortical and cerebellar regions. We also construct a polygenic index of SES to control for the additive effects of common genetic variation related to SES, which attenuates observed SES-GMV relations, to different degrees in different areas. Remaining variance, which may be attributable to environmental factors, is substantially accounted for by body mass index, a marker for lifestyle related to SES. In sum, SES affects multiple brain regions through measurable genetic and environmental effects.

**One-sentence Summary:** Socioeconomic status is linked with brain anatomy through a varying balance of genetic and environmental influences.

## Main Text

Socioeconomic status (SES), typically measured by income, education, occupation and neighborhood quality, is a powerful predictor of important life outcomes including physical and mental health, academic achievement and cognitive abilities (*1–5*). The brain plays a central role in these relations, most obviously in mental health and intellectual capabilities, but also in physical health through neuroendocrine and inflammatory pathways (*6, 7*). Thus, neuroscience provides a window on the biosocial pathways linking SES and human health and capabilities.

Neuroscience research on SES has revealed a generally positive relation with overall brain volume, as well as with regional cortical and subcortical volumes and cortical surface areas (*8–10*). We note variability across studies in the regions most associated with SES, which may be due in part to the relatively small samples studied and in part to differences in the ways SES has been measured and analyzed (e.g., choices of covariates) (*10, 11*). One of the goals of the present study is to establish the relation of SES to regional grey matter volumes (GMV), in the largest sample so far examined for voxel-level data, using a comprehensive measure of SES, controls for a number of potential confounds, and a well-powered, pre-registered analysis plan.

The second goal of the study is to differentiate genetic from environmental causes of the SES-GMV relation. As hinted by recent studies (*12–14*), both kinds of mechanisms are plausible. The possibility of environmental influence on brain structure is shown by animal studies in which features of lower SES environments, such as poor nutrition, environmental toxins, chronic stress and limited cognitive stimulation, are experimentally manipulated and found to impact brain structure (*15*), as well as the rare experimental study with human brains (*16, 17*). The possibility of genetic influence is shown by the influence of genes on both human brain structure (*18, 19*) and aspects of SES (*20–22*).

Here, we pursue these two goals using data from the UK Biobank (UKB), a large-scale prospective epidemiological study of individuals aged 40–69 years at recruitment (*23, 24*). After selecting participants who had undergone both MRI and genotyping, and had complete SES information related to occupation, income, education, and neighborhood quality, we excluded participants with clinical diagnoses related to brain pathology, morbid obesity, heavy alcohol drinking, or low data quality (*25*). The resulting sample was 23,931 individuals, with a mean age of 62, 57% of whom were female. This sample size provides 90% statistical power to detect effects as small as *R*^2^ > 0.17% at the 5% significance level (corrected for multiple testing; uncorrected *p* < 2.19×10^-6^). We conducted voxel-based morphometry (VBM) analysis of grey matter volumes (GMV). T1 images were preprocessed with the Computational Anatomy Toolbox (CAT) 12, and anatomical regions were labeled according to the Neuromorphometrics atlas (*26*).

SES was represented in the analyses to follow by two summary measures, derived from available SES variables using a generalized version of principal component (PC) analysis (Fig 1 and S2). This approach better accommodates measurement error and allows us to appreciate the multidimensional nature of SES with just two components. *PC1_SES_* mainly captures the positive correlations between the different SES measures and is most strongly influenced by occupations, occupational wages, and education. *PC2_SES_* primarily reflects occupations and neighborhood qualities that are not strongly linked with educational attainment or income, e.g., individuals who live in relatively poor neighborhoods despite having high educational attainment. As shown later, *PC2_SES_* contributes to capturing non-genetic variation in SES.

**Fig. 1.**
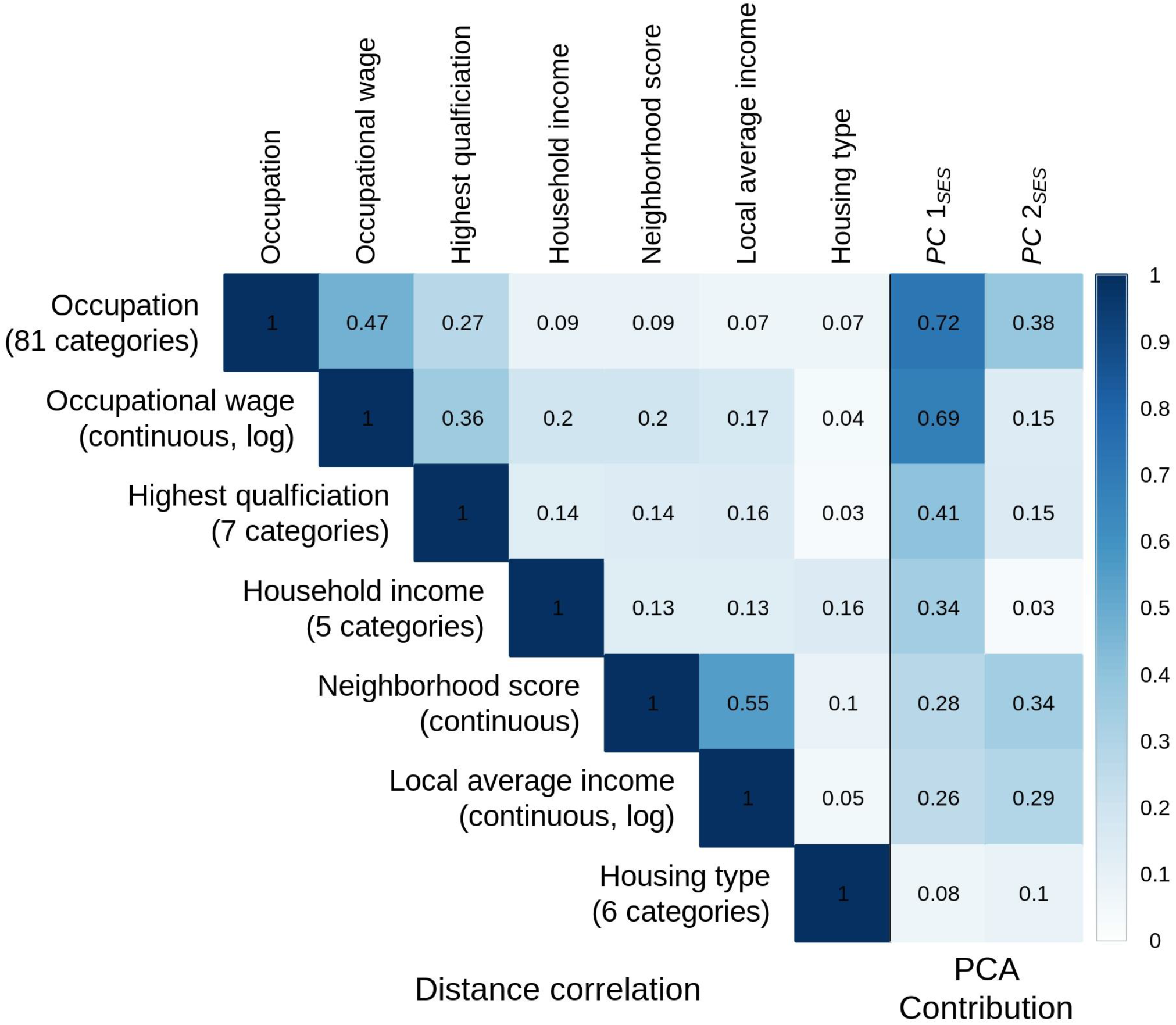
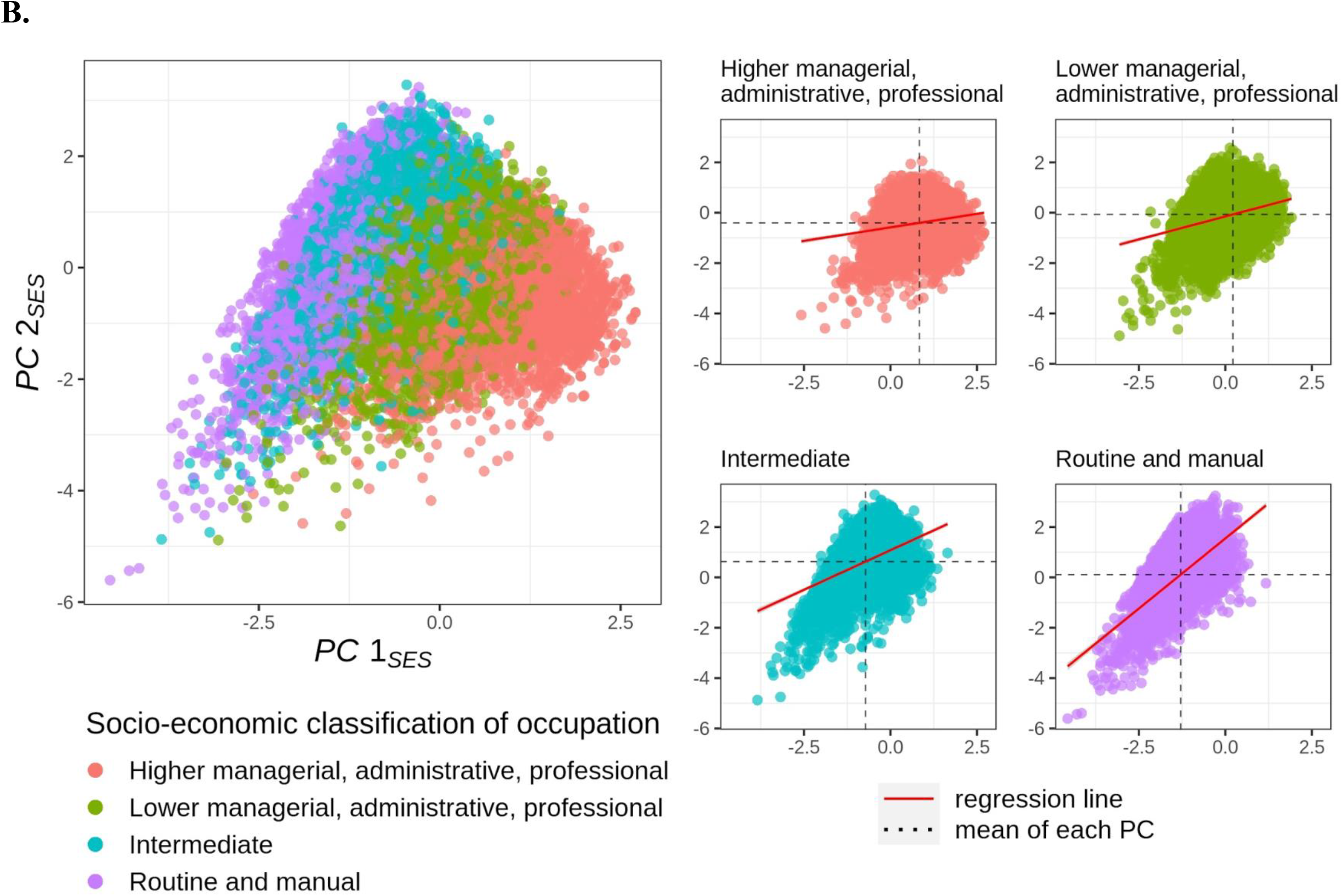
Measures of socioeconomic status and principal component analysis. **A.** On the left, a distance correlation matrix is plotted for seven indices of socioeconomic status (SES). On the right, the squared loadings for each principal component are indicated. **B.** Scatter plots of the first principal component (*PC1*_SES_) against the second component (*PC2*_SES_). The points in different colors represent four SES groups defined by National Statistics Socio-economic Classification, which are approximately clustered by the two PCs. On the right, the same scatter plots are presented for each SES group. The mean values of each PC are indicated for each group. The regression lines are plotted to describe that SES is more complex for the lower SES groups.

We first examined the relation between total intracranial volume (TIV) and SES by regressing TIV on *PC1*_SES_ and *PC2*_SES_, controlling for sex, age, genetic population structure, and a number of image-related technical covariates (*25*). *PC1*_SES_ is positively associated with TIV (standardized *β*= 0.10; *p* = 1.1⨉10^-87^; 95% CI [0.09, 0.11]), while for *PC2*_SES_ the relation is statistically indistinguishable from zero (standardized *β*= 0.01; *p* = 0.14; 95% CI [-0.00, 0.02]). The two PCs together explain 1.6% of the variance of interest in TIV beyond the covariates of no interest (partial *R^2^*)—slightly higher than TIV’s relation to educational attainment (1.4%), and lower than its relation to fluid intelligence (2.6%) (*27*).

Next, we conducted VBM analysis to test the association of these two PCs with regional GMV across the brain, using the same set of covariates. Higher SES is associated with larger GMV across the brain (Fig 2A). 89.5% of the voxels have a statistically significant association with SES at a familywise error rate of 5%, all of which are positive. For statistically significant voxels the average partial *R^2^* is 0.4% and the highest is 1.2%, with the strongest associations in the left ventral striatum and the right frontal pole. Thus, the positive relation between total brain volume and SES arises from many relatively small sources of structural variation that are widespread across the brain.

**Fig. 2.**
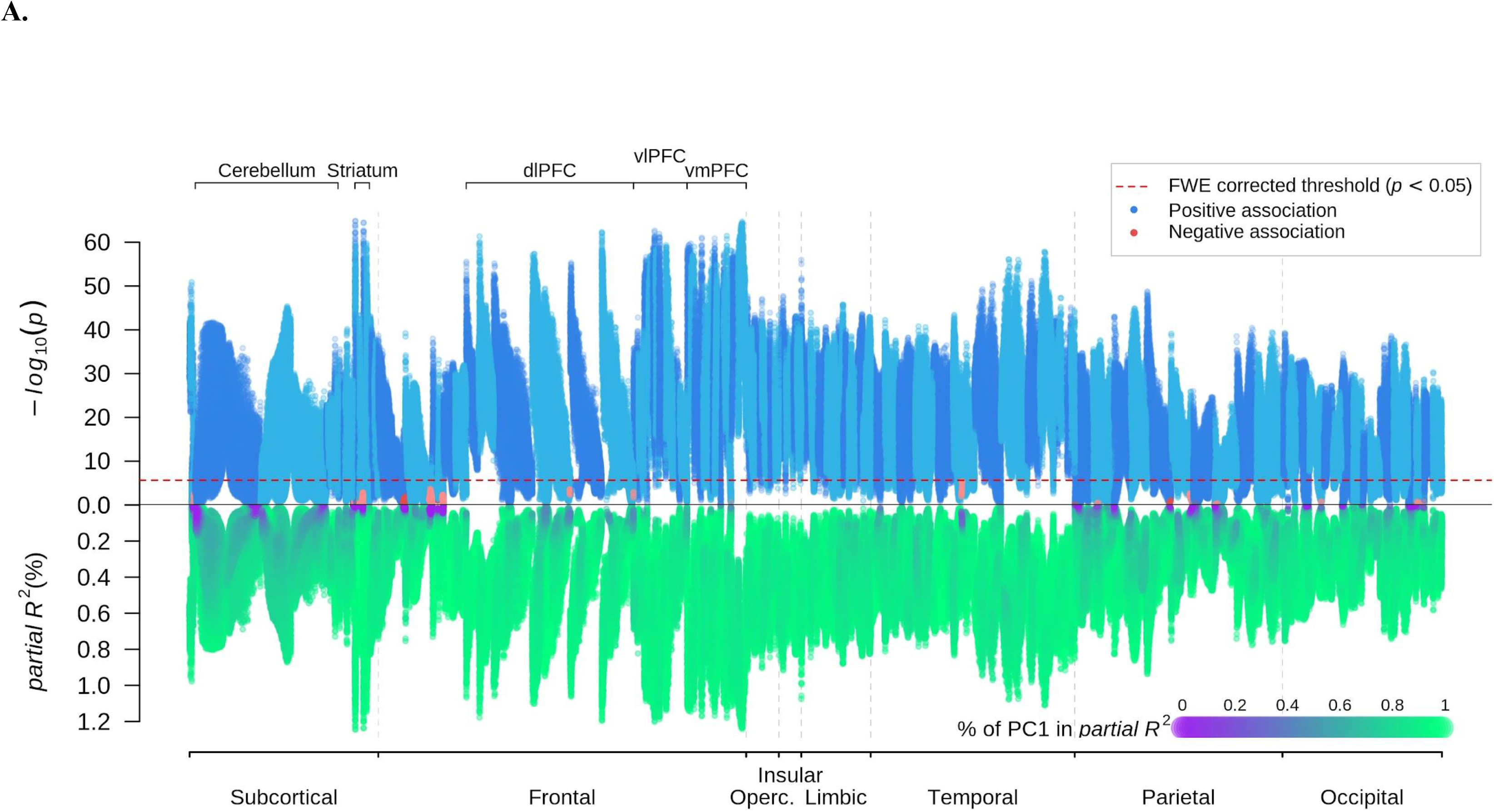
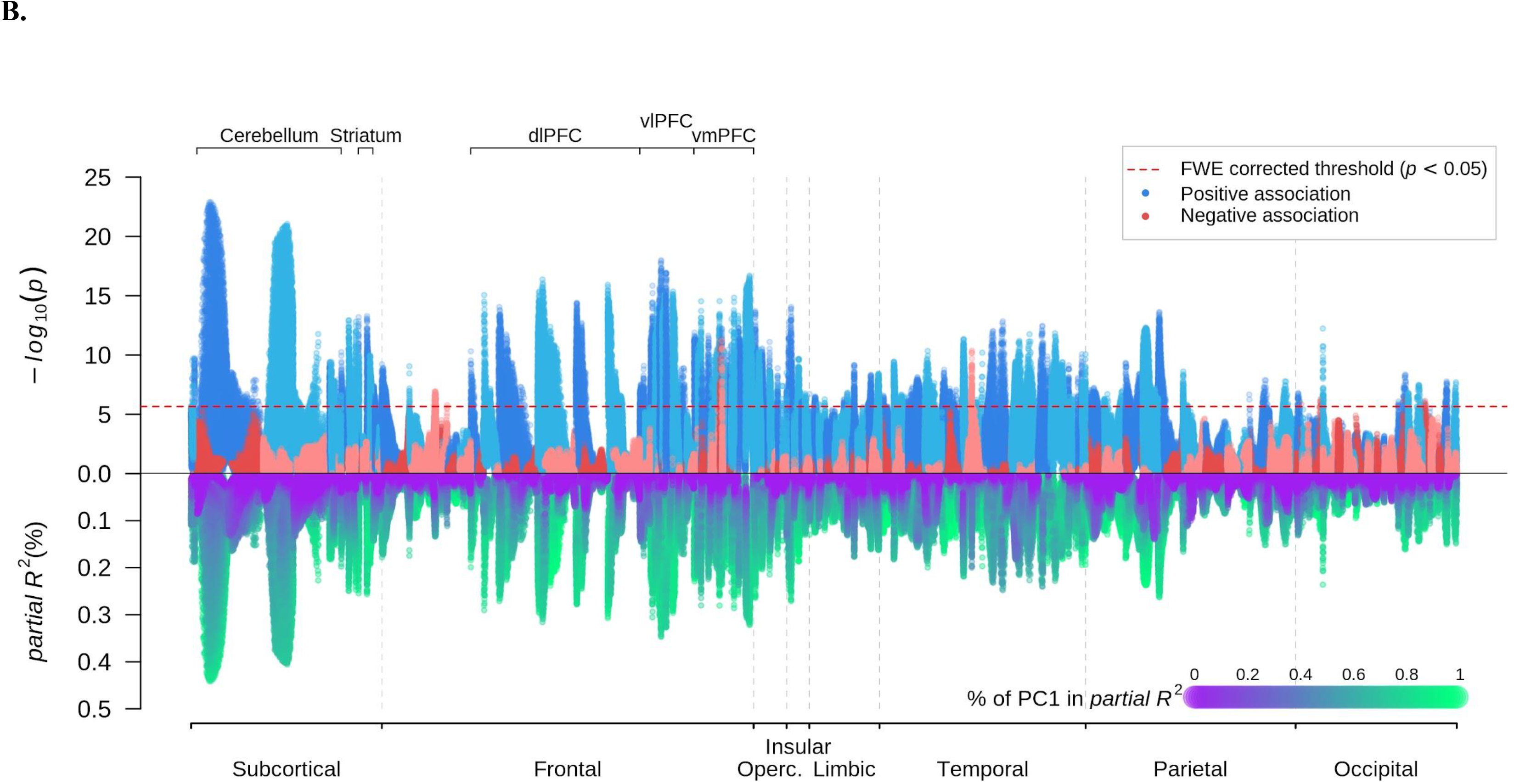
Manhattan plots: voxel-based morphometry of grey matter volume and socioeconomic status. **A.** Univariate voxel-based morphometry (VBM) results on the two principal components (PC) for socioeconomic status. These regressions did not control for total intracranial volume (TIV). *p*-values on *log_10_* scale (upper) and partial *R^2^* (lower) are plotted for each voxel. The sign of the association is that of the first PC. The voxels were anatomically labelled according to the Neuromorphometrics atlas and grouped by the labelled regions. Within each region, the voxels were ordered by their distance to the medoid of their region. **B.** Univariate VBM results with TIV controlled for.

Accordingly, when TIV is controlled for, just 8.5% of the voxels have a statistically significant association with SES and the average effect size in partial *R^2^* is reduced by over half to 0.17% for statistically significant voxels. As shown in Fig 2B, the strongest positive associations between SES and relative GMV fall in the prefrontal, insular, frontal opercular, lateral parietal, and lateral temporal regions, as well as in subcortical areas including the cerebellum, striatum, and thalamus. While SES-GMV associations are mainly driven by *PC1*_SES_, *PC2*_SES_ contribute relatively more in lateral temporal, cerebellar, and ventromedial prefrontal regions than in other regions (Fig 2B and S4A).

The regions implicated in these analyses include many reported in previous studies of SES and brain structure. While the cerebellum has not often been linked to SES, this may reflect its omission from many morphometric studies (but see (*28*), for a study of SES and cerebellar volume specifically, with positive findings). Conversely, hippocampus volume is often noted to correlate with SES. Although this was also found in the present study, it was not among the strongest relations.

We also explored the influence of individual aspects of SES, such as education and income, by conducting a cluster-based analysis (Fig S8-S9) as well as VBM on each measure separately (Fig S5 and S11). The overall pattern of results is similar, with years of schooling being most strongly associated (*25*).

SES-health relations are often stronger at lower levels of SES, where more extreme deprivation may impose unique effects on health (*29, 30*) and this pattern is also seen in SES effects on the cortex in children (*31*). Stronger SES-GMV associations were found here in the lower SES participants of our sample as well (Fig S6) (*32*). Regionally, this is particularly apparent in the striatum (low SES, max partial *R^2^* = 0.65%, TIV adjusted; high SES, max partial *R^2^* = 0.17%, TIV adjusted) (*25*).

An alternative measure of the strength of the SES-GMV relation is the ability of aggregate GMV measures to predict SES. Indeed, the small effect sizes for individual voxels do not imply that the association between SES and overall GMV structure is also small. To show this, we constructed brainwide GMV scores to predict *PC1*_SES_ and *PC2*_SES_ via a stacked block ridge regression (*33*) with 5-fold cross-validation. These scores predict Δ*R^2^* = 4.9% (95% CI [4.4, 5.4]) of out-of-sample variation in *PC1*_SES_ and Δ*R^2^* = 0.5% (95% CI [0.3, 0.7]) in *PC2*_SES_ (*25*).

The second question to be addressed is the contribution of genetic and environmental influence to the SES-GMV relations reported here. We approached this by first estimating the SNP-based heritability of SES and brain measures as well as the pairwise genetic correlations among them, which indicated that the genetic architectures of SES and brain structure are partly overlapping. We then constructed a polygenic index for SES (*PGI*_SES_) using the results of the genome-wide association study (GWAS) from the UKB. In view of the sensitivity of GWAS results to differences in ancestry, we derived the index from UKB participants of European ancestry only, excluding the scanned participants and other participants genetically related to them. The genetic data consisted of relatively common genetic variants (single-nucleotide polymorphisms or SNPs) with minor allele frequency ≥1%, which were related to educational attainment, occupational wages, household income, local average income, and neighborhood quality, combined using Genomic SEM (*21, 25, 34*) (effective *N* = 849,744). *PGI*_SES_ was strongly associated with *PC1*_SES_ (Δ*R^2^* = 7.1%, *p* < 10^-300^) and weakly with *PC2*_SES_ (Δ*R^2^* = 0.02%, *p* = 0.03). *PGI*_SES_ could then be used with images from participants of EA (*N* = 20,799) to help discriminate genetic from environmental causes of GMV differences.

*PGI*_SES_ was then used to predict TIV (Δ*R^2^* = 0.8%, *p =* 7.4⨉10^-64^) and GMV across the entire brain via VBM. The latter analysis revealed positive associations in widely distributed voxels (Fig 3A b.), with the most pronounced associations in the anterior insula, frontal operculum, prefrontal, anterior cingulate, and striatum. There is substantial overlap between the neuroanatomical correlates of SES and *PGI*_SES_. Controlling for TIV, approximately 41% of the GMV voxels associated with SES are also associated with *PGI*_SES_. This overlap is especially apparent in the insular and prefrontal cortices, with roughly 96% and 64% of the voxels associated with *PC*_SES_ also associated with *PGI*_SES_, respectively.

**Fig. 3.**
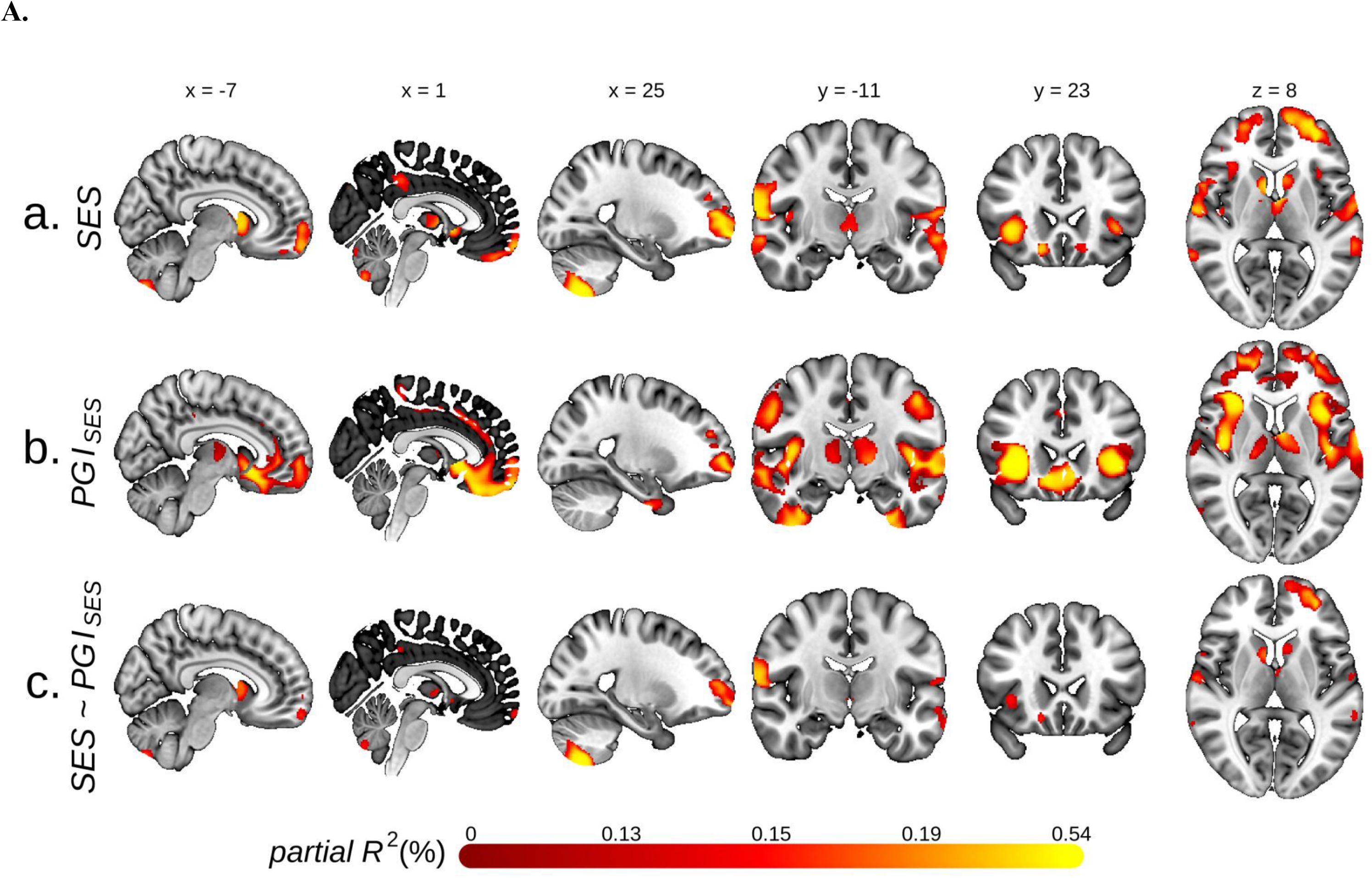
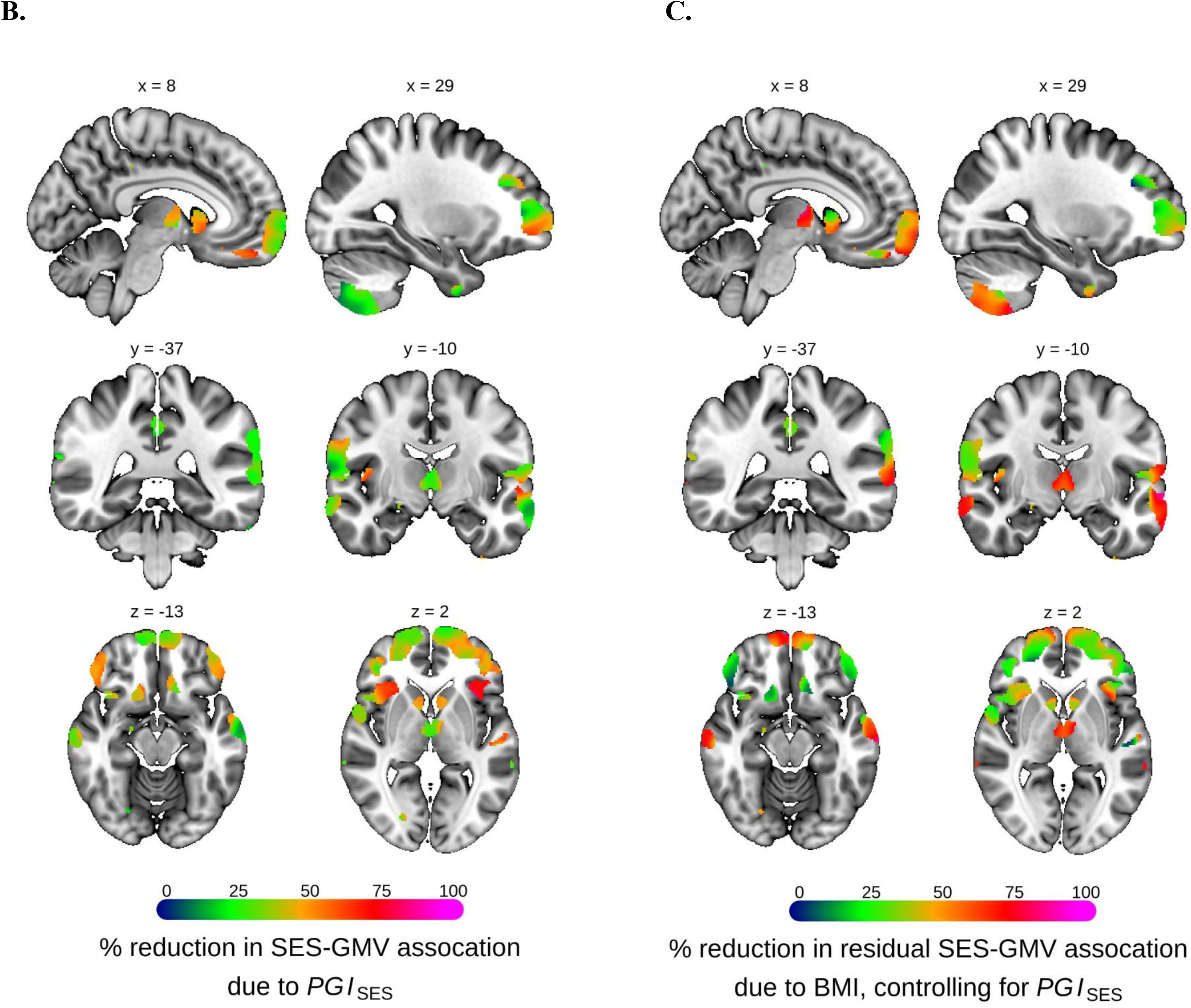
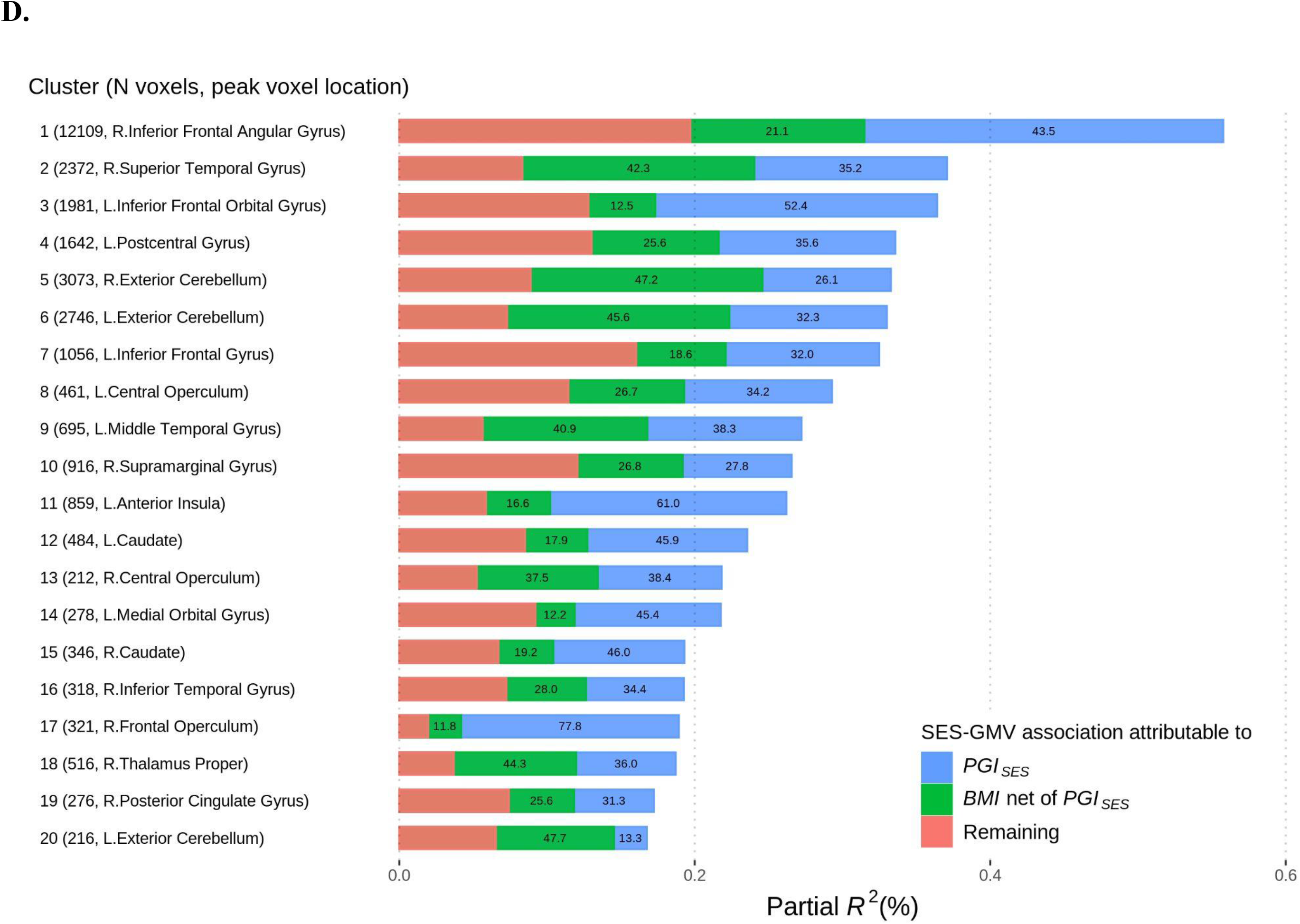
Voxel-based morphometry of socioeconomic status (SES) and its genetic and environmental components. The sample was restricted to individuals of European ancestry. **A.** Univariate voxel-based morphometry (VBM) results, with grey matter volume (GMV) as the dependent variable. Voxels significant at FWE rate of 5% level are plotted for: **a.** the two principal components (PC) measuring socioeconomic status (*SES*), **b.** the polygenic index for SES (*PGI_SES_*), **c.** SES while controlling for *PGI_SES_*. **B.** Percent reduction in the association between GMV and the two PC for SES due to controlling for *PGI_SES_*. **C.** Percent reduction due to controlling for body mass index (BMI) in the residual association between GMV and the two PC for SES after controlling for *PGI_SES_*. The figures plot only voxels which had significant SES-GMV association before *PGI_SES_* and BMI were controlled for. **D.** Associations in partial *R^2^* between the two PC for SES and GMV in voxel clusters attributable to *PGI_SES_* and BMI. The numbers in the bars report the percent share in the SES-GMV association attributable to *PGI_SES_* or BMI partialled out of *PGI_SES_*. The clusters were formed from the VBM results plotted in **A.a.** See Table S9 for more information about the clusters. MNI coordinates are indicated for **A.** and **B.** Measurement error in *PGI_SES_* was adjusted for with genetic instrument variable regression for **B.** and **C.**

We then examined to which extent the shared common genetic architectures of SES and GMV account for the observed phenotypic associations by comparing TIV-adjusted regression results of GMV on SES with and without controlling for *PGI*_SES_. For 13% of the voxels significantly associated with SES before *PGI*_SES_ is controlled for, there is a statistically significant change in at least one of the coefficients for *PC1*_SES_ and *PC2*_SES_ after accounting for *PGI*_SES_ (*25*). Controlling for *PGI*_SES_ reduces the SES-GMV associations across the entire brain, with the greatest reduction in the anterior insula, frontal operculum, ventrolateral prefrontal cortex, and ventral striatum of both hemispheres, consistent with VBM of *PGI*_SES_ mentioned earlier (Fig 3B). When we correct for measurement error in *PGI*_SES_ using genetic instrumental variable regression (*35*), we estimate that *PGI*_SES_ accounts for more than half of the SES-GMV associations for many of these regions. On average, 38% of the SES-GMV associations (*min* = 3%, *max* = 87%) can be attributed to *PGI*_SES_ (*25*).

The remaining associations between GMV and SES could be either due to environmental influences on both or due to rare SNPs, structural variants (e.g. inversions, deletions), or interactions among genes (i.e. epistasis) that *PGI*_SES_ does not fully account for. Forty-three percent of the voxels significantly associated with SES fall into this category, remaining associated with SES after controlling for *PGI*_SES_ (Fig 3A c.). The SES-GMV association is least attenuated by genetic controls in the cerebellum and lateral temporal, lateral parietal, posterior cingulate and primary motor regions, as well as some areas of the dorsolateral and ventromedial prefrontal cortex (vmPFC) and the thalamus. Controlling for *PGI*_SES_ accounts for less than 30% of the SES-GMV association in many of these regions. These results suggest that the aforementioned regions may be particularly susceptible to the influence of the socioeconomic environment. This is consistent with the relatively stronger association of *PC2*_SES_ to GMV in many of these areas, as *PC2*_SES_ was found to be barely heritable (*25*). In sum, a substantial portion of the SES-GMV relation is attributable to known genetics, and that portion varies according to region of the brain. The remaining portion of this relation is also substantial, and likely includes the effects of the environment.

We then annotated the brain regions whose morphology was related to SES or its genetic influences by relating them spatially to fMRI meta-analyses of cognitive function. We used 506 concepts from the Cognitive atlas knowledge base and obtained a meta-analyzed Z-score brain map for each concept using the tool NeuroQuery (*25, 36, 37*). For each concept-associated brain-map, we calculated the mean *χ*^2^ based on voxels that overlapped with the VBM results (Fig 4). Prominent effects of genetic influences (*PGI*_SES_) are seen in brain regions associated with sensory cognitive functions, risk processing, and motivation. The regions found to be likely more environmentally susceptible (i.e. *SES PC ∼ PGI_SES_*) reflect cognitive concepts related to executive control, working memory, and subjective well-being.

**Fig. 4.**
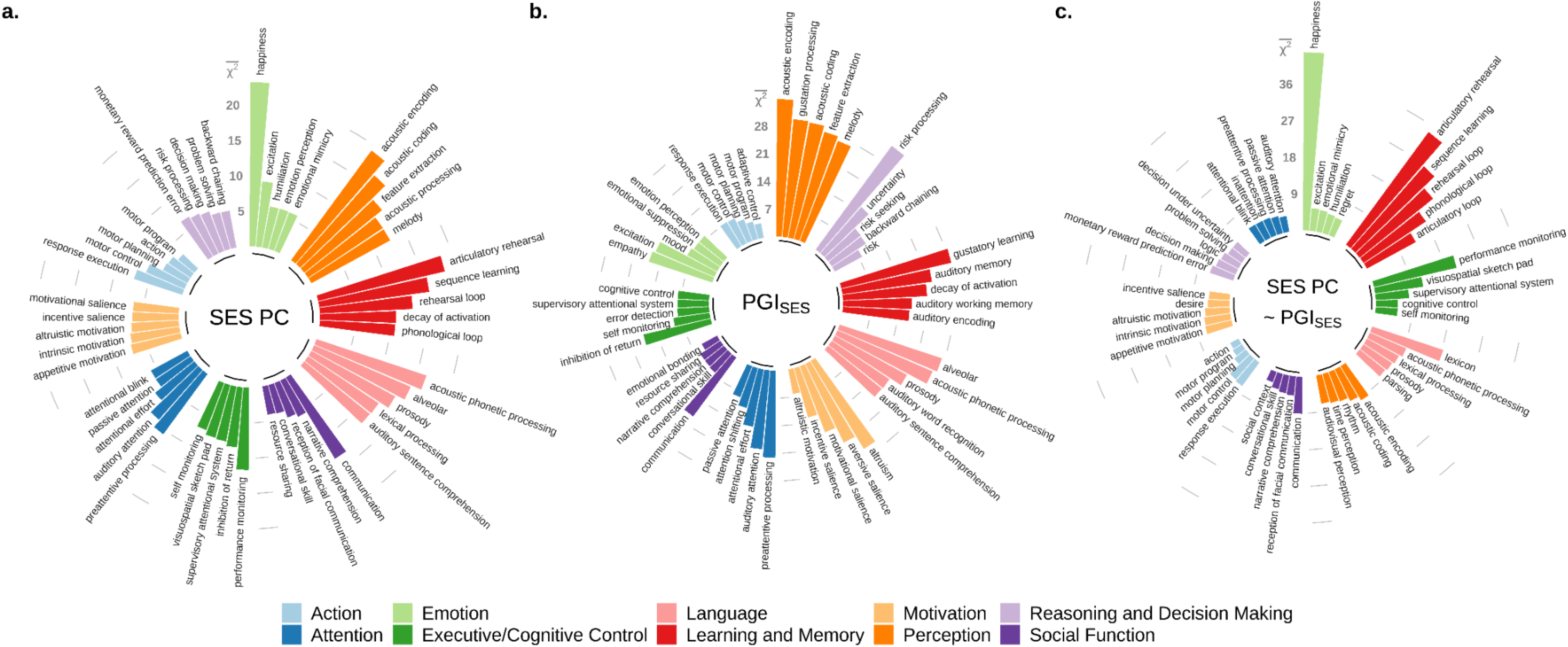
Functional annotation of brain regions associated with socioeconomic status (SES) 506 cognitive concepts, belonging to 11 categories, were taken from Cognitive Atlas and their predicted fMRI meta-analysis results were generated by NeuroQuery. For each concept, mean *χ*^2^ was computed with voxels statistically significant at the FWE rate of 5% level in the voxel-based morphometry results respectively for **a.** two principal components (PC) of SES, **b.** the polygenic index for SES (*PGI_SES_*), and **c.** SES controlling for *PGI_SES_*. The results for the top five concepts from each category were plotted.

We sought to strengthen evidence concerning environmental causation through the study of a specific environmental factor. Numerous environmental exposures are associated with SES and are plausible causal contributors to the SES-GMV relation found here. These include prenatal and childhood factors with lifelong effects, as well as adulthood exposures such as chronic life stress, nutritional status, physical exercise, environmental toxins, smoking and other substance use. Experimental research with animals and human research with longitudinal, quasi-experimental or experimental studies show that these are all capable of impacting the brain. On the basis of recent research with the same sample relating mid-life obesity to cognitive and brain aging (*38*), we chose body mass index (BMI) as marker for a set of lifestyle factors that could mediate the SES-GMV relation, including nutrition, physical activity, and obesity, which can impact the brain through their downstream effects on blood pressure, blood lipids, glucose metabolism and inflammation. In addition to the logical point that *PGI*_SES_ controls would account for genetic influences of BMI on the SES-GMV relation, there is also experimental evidence of SES affecting BMI through the environment: increasing SES causes BMI to decrease (*39*).

BMI accounts for an average of 44% of the SES-GMV associations that remain after controlling for *PGI*_SES_ (Fig 3C-D). This effect is not due to neurological disease associated with BMI, such as stroke or neurodegenerative disease, because neurological disease was an exclusionary criterion for our sample. The effect is particularly large in the thalamus and the cerebellum as well as the lateral temporal region and some areas of the vmPFC. Furthermore, for the 91% of the voxels with significant SES-GMV association in the European ancestry sample, at least 50% of the estimated SES-GMV association can be attributed to *PGI_SES_* and BMI combined, with 67% on average.

Taken together, our results suggest that SES is linked with brain anatomy through a combination of genetic and environmental influences. The balance of genetic and environmental influence varies across brain regions. This insight suggests that brain health is more susceptible to SES-related stressors in specific regions, including the cerebellum.

## Acknowledgments

This research was carried out under the auspices of the Brain Imaging and Genetics in Behavioral Research Consortium (https://big-bear-research.org/), using UK Biobank resources under application 11425. The study was supported by funding from an ERC Consolidator Grant to PK (647648 EdGe), an NSF Early Career Development Program grant (1942917) to GN, the School of Arts and Sciences at the University of Pennsylvania to MJF, and the Wisconsin Alumni Research Foundation, provided to PK by the Office of the Vice Chancellor for Research and Graduate Education at the University of Wisconsin–Madison. We thank James Gee and Philip Cook of the Penn Image Computing and Science Lab for valuable advice on image analysis. Steven Hyman kindly provided comments on an earlier draft of this manuscript. GN thanks Carlos and Rosa de la Cruz for ongoing support.

## Author contributions

HK, GN, MJF, and PDK designed the research plan. GN, MJF, and PDK oversaw the study. HK analyzed the data. HK, MJF, and PDK wrote the paper and supplementary note. GA, AD, DB, and CCR provided helpful advice and feedback on the study design. All authors contributed to and critically reviewed the manuscript.

## Data and materials availability

Data and materials are available via UKB at http://www.ukbiobank.ac.uk. The analysis code used in this study is publicly available at https://osf.io/kg29c.

## Supplementary Note

### 1. Study overview

In this study, we aim to answer two research questions following a pre-registered analysis plan (https://osf.io/kg29c/):

1. Are there robust associations between socio-economic status (SES) and brain anatomy?
2. How much of the association between SES and brain anatomy is due to common genetic factors that are linked to SES?

To this end, we conducted voxel-based morphometry (VBM) analysis of grey matter volumes (GMV) on socioeconomic status (SES), using a population sample of 23,931 older adults from the UK Biobank (UKB) that contains brain images, genetic data, and several measures of SES (*23, 24*).

Our phenotypic measures of SES uses all available information on SES that is available in the UKB and fully recognizes the multidimensional nature of SES. In particular, we measure SES as the first two principal components (PC) of available indices of household income, occupations, neighborhood, and education. These PCs were then jointly tested for their association with voxel-level GMVs in a univariate VBM. Permutation testing was used to maintain a familywise error rate of 5%.

To investigate the genetic basis of the estimated SES-GMV associations, we constructed a polygenic index for SES (*PGI*_SES_) derived from multiple genome-wide association studies (effective *N*=849,744) and adjusted for measurement error in *PGI*_SES_ using genetic instrumental variable regression (*35*). We then examined to which extent the estimated SES-GMV associations can be attributed to the shared common genetic architectures of SES and the GMV structure by comparing regression results of GMV on SES before and after controlling for the polygenic index for SES.

Our analysis was carried out under the auspices of the Brain Imaging and Genetics in Behavioral Research Consortium (https://big-bear-research.org/).

### 2. Sample description

We used publicly available data from the UKB, which recruited ≈500,000 participants from the general population of the UK (*23, 40*). Study participants were 40-69 years old at recruitment between 2006-2010. Our study sample originates from 40,681 individuals whose structural T1 MRI images were available in January 2020 (data field 20252). To derive voxel-level grey matter volumes, we processed T1 images from 38,545 genotyped individuals with the Computational Anatomy Toolbox (CAT) 12 for SPM (see Section 3.1 for details). We then applied several filters to ensure data quality and avoid spurious findings.

We excluded:

- 24 individuals with mismatch between genetic (data field 22001) and self reported sex (data field 31)
- 1,818 individuals with clinical diagnoses related to brain pathology (including dementia, Alzheimer’s, Parkinson’s, and chronic degenerative neurological diseases, Guillan-Barré syndrome, multiple sclerosis, other demyelinating diseases, stroke or ischaemic stroke, brain cancer, brain haemorrhage, brain or intracranial abscess, cerebral aneurysm, cerebral palsy, encephalitis, epilepsy, head injury, infections of the nervous system, meningeal cancer, meningioma, meningitis, ALS, neurological injury or trauma, spina bifida, subdural haematoma, subarachnoid haemorrhage, or transient ischaemic attack; see the pre-registered plan for ICD10 codes - https://osf.io/kg29c)
- 2,705 individuals who were morbidly obese (BMI > 35)
- 2,427 individuals who were current heavy drinkers, where heavy drinking is defined as consuming more than 24 drinks per week for males and more than 18 drinks per week for females (*41, 42*)^1^
- 11 individuals with the image-quality rating (computed by CAT12) lower than “C”
- 230 individuals with a sample homogeneity measure (mean voxel correlation^2^) lower than the 1% quantile (0.805 based on all 38,545 individuals)

After applying these exclusion criteria, 31,330 individuals remained in our sample. 7,215 individuals were further excluded due to missing data for the variables listed in Section 3. In order to rule out that our results are influenced by shared family environments among related individuals, we also removed close relatives by randomly dropping one from each pair of siblings or parent-offsprings. Our final sample for the main analysis included *N* = 23,931 individuals. In analyses that employed genetic data, we included *N* = 20,799 individuals of European ancestry from this sample.

### 3. Measures

#### 3.1. Imaging-derived phenotypes (IDPs)

We extracted GMV on the voxel level from T1-weighted structural brain MRI images provided in NIFTI format (data field 20252). The UKB scanned the participants with a Siemens Skyra 3T scanner using a standard 32-channel head coil (Siemens Healthcare, Erlangen, Germany) in three assessment centers (Cheadle, Newcastle, and Reading). The scanning and processing protocols are detailed in the UKB’s brain imaging documentation (https://biobank.ctsu.ox.ac.uk/crystal/crystal/docs/brain_mri.pdf) as well as in publications (*23, 43*).

We first pre-processed the T1 images with the Computational Anatomy Toolbox (CAT) 12 for SPM (www.fil.ion.ucl.ac.uk/spm/software/spm12/). The images were corrected for bias-field inhomogeneities, tissue-segmented, spatially normalized to the MNI space with 1.5mm resolution by linear and non-linear transformations, and were modulated to ensure that the total amount of signal in the original image was preserved during spatial normalization. 8-mm Full-Width-at-Half-Maximum Gaussian kernel was then used to spatially smooth the pre-processed images. More details can be found in our pre-registered analysis plan (https://osf.io/kg29c/) as well as in the recent publications of BIG BEAR consortium (*41, 42*).

Following the standard VBM procedures (see e.g. SPM/CAT12 http://www.neuro.uni-jena.de/cat12/CAT12-Manual.pdf), we decided to exclude all voxels from the VBM analyses that did not contain any or sufficient grey matter. To determine this, we first computed the average of all GMV images and then thresholded the average brain image at 250 GMV intensity units. The resulting binarized image was then applied as a pre-mask to all individual images. After applying a gray matter mask derived from the data and dropping the voxels unlikely to contain any gray matter (*41*), GMV estimates from 504,426 voxels were used in the analysis.

#### 3.2. SES measures

The UKB offers a rich set of SES indicators, including education, income, occupation, and neighborhood quality. In order to make full and efficient use of the data, we took a data-driven approach to measure SES by extracting principal components (PC) that capture overall SES from all SES measures available in the UKB. Our approach can be summarized as follows: We (1) collected every available source of information relevant to SES in the UKB; (2) combined measures or derived new variables when possible or appropriate; (3) extracted PCs that represent a sparse, but accurate overall measure of SES; (4) and jointly tested for neuroanatomical association of these PCs based on an *F*-test.

There are several important reasons that motivated us to use this approach. First, it allowed us to take into full account the multidimensional nature of SES. While each SES dimension tends to share the same direction of correlation, there often are cases that do not agree with such correlation in reality: for instance, a plumber may have less education than a university lecturer, but may earn higher income. Furthermore, the quality of a neighborhood in which an individual lives is an important dimension of SES, but it may be imperfectly correlated with education and income. Such complex aspects of SES cannot be represented by a single SES measure such as education or income alone.

Second, our data-driven approach is useful for efficiently testing for the association between SES and neuroanatomy by summarizing the available measures and thereby decreasing the multiple testing burden and increasing the statistical power of our analyses.

Third, this approach also makes it possible to use the detailed occupation data of the UKB to a fuller extent. Because it is difficult to handle many occupational categories in a single analysis, studies often use an aggregated summary of occupation by classifying occupations into a small number of predefined categories.^3^ Such a predefined classification can discard potentially useful information and may not truly represent different levels of SES. A data-driven approach can efficiently reduce wide categorical data of occupation into lower continuous dimensions, while minimizing information loss.

Fourth, our approach addresses important limitations of educational attainment measures in the study sample. In the UKB, qualifications are reported in only six non-hierarchical categories, some of which cover a wide range of educational levels. Furthermore, participants were allowed to indicate multiple categories without a specific instruction, which led to a large degree of variation in responses. For this reason, we chose not to use years of schooling as often done (*21*), but instead determined the highest qualification for each participant in a data-driven way and used it as a categorical variable.

##### 3.2.1. Available measures of SES in the UK Biobank

We collected and constructed an extensive set of SES measures as described in the table below. We derived some of the variables by relying on external data sources or aggregating several measures. The participants visited the assessment center up to four times and brain images were taken during the third or fourth visit (the fourth visit was for repeated imaging of a subset of participants). While the data used here was primarily collected during the brain imaging visit at the assessment centers, we used the latest available information if a measure was missing from this visit.

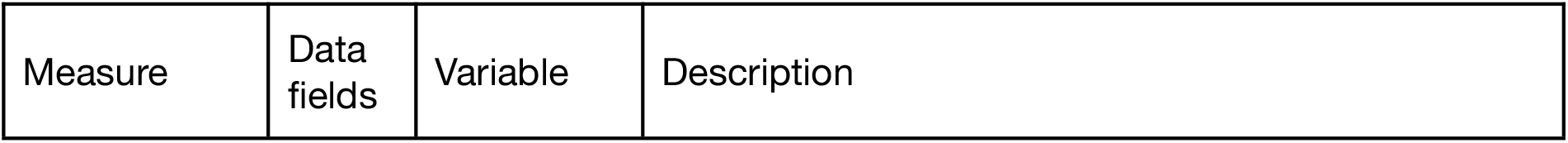
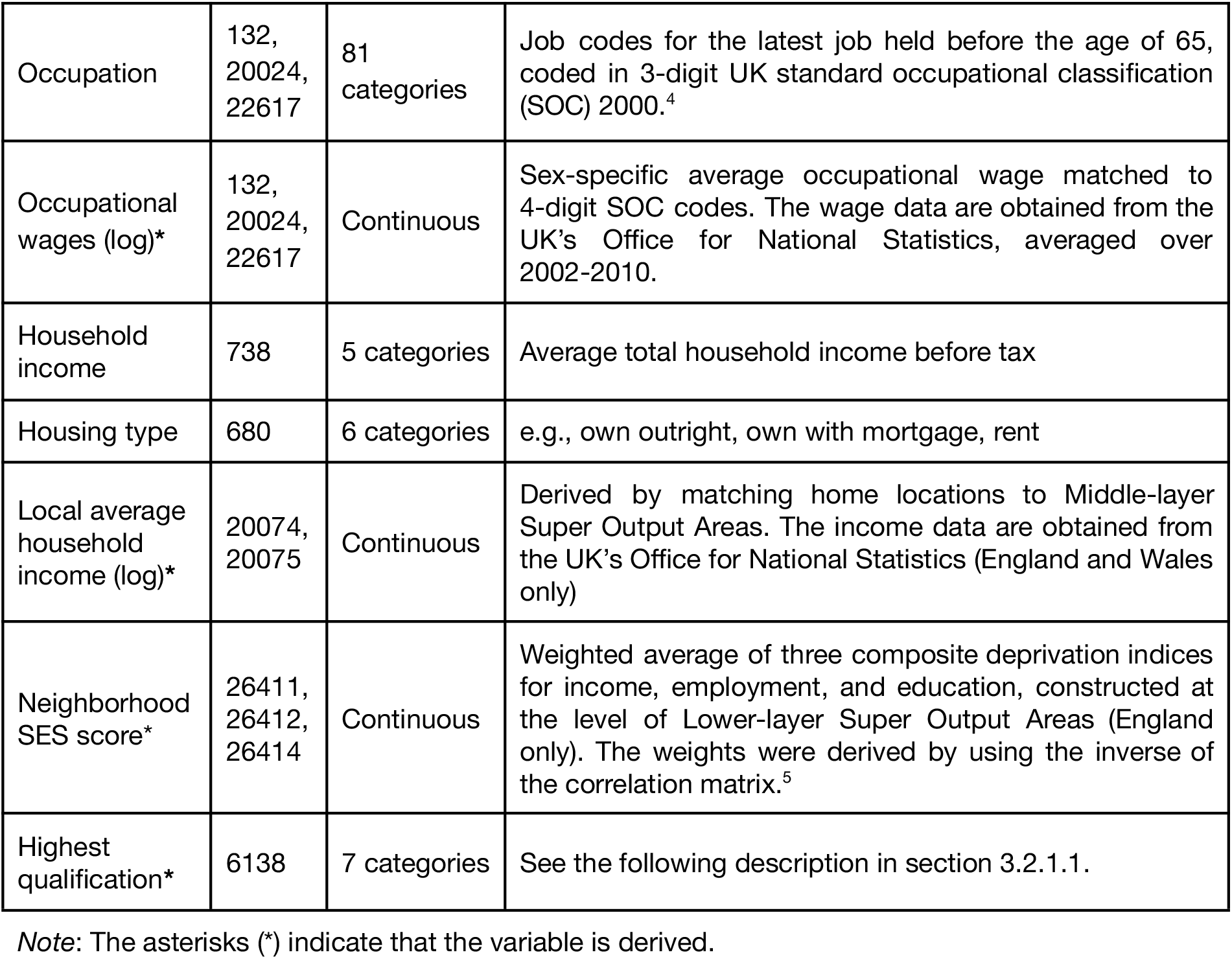

###### 3.2.1.1. Highest qualification

This subsection describes how we derived the highest qualification. During the assessment, participants were asked to choose qualifications that they have from the below options:

1. *College or University degree*
2. *A levels/AS levels or equivalent*
3. *O levels/GCSEs or equivalent*
4. *CSEs or equivalent*
5. *NVQ or HND or HNC or equivalent*
6. *Other professional qualifications eg: nursing, teaching*
7. *None of the above*

Because participants were able to choose multiple qualifications and also because the vocational category (NVQ or HND or HNC or equivalent) covers an extensive range of educational levels, it was not straightforward to determine the highest qualification for qualifications below college degree. Preferably, a better qualification should correspond to a better SES. We therefore used the following procedure to determine the rank of each qualification.

We first created a new categorical qualification variable that treats each combination of multiple choices as a unique response. Using the method described in Section 3.2.2, we extracted the first PC from this variable along with the rest of SES variables listed above. The average of the first PC was then computed for each qualification from a group of people who reported having that qualification. Note that these groups are not mutually exclusive as the individuals can belong to multiple groups. We then determined the SES-rank of qualifications based on these average PC scores. This approach yielded the following ranking:

1 *College or University degree*
2 *A levels/AS levels or equivalent*
6 *Other professional qualifications eg: nursing, teaching*
3 *O levels/GCSEs or equivalent*
5 *NVQ or HND or HNC or equivalent*
4 *CSEs or equivalent*
7 *None of the above*

The highest qualification was chosen for each individual according to this rank, which was then included as a categorical variable in the principal component analysis described below.

##### 3.2.2. Data reduction by principal component analysis

We reduced the dimensions of the data by extracting PCs, which represent overall SES implied by the available indicators. Standard principal component analysis (PCA) is only suitable for non-categorical data. Thus, to account for the fact that we have both non-categorical and categorical SES indicators, we employed a method that is often called factorial analysis of mixed data, which is essentially a generalization of PCA that can handle such mixed data (*45, 46*). This method combines ordinary PCA for non-categorical data with multiple correspondence analysis for categorical data and is implemented in the R package PCAmix (*47*).

Our purpose here was not a factor extraction that finds all relevant factors as typically done, but to exploit only the most meaningful variation in the UKB’s SES data to facilitate efficient discovery of neuroanatomical correlates of SES. For this purpose, it was optimal to use the minimal number of PCs that could sufficiently capture the multidimensional nature of SES. Given this objective, we used the first two PCs (*PC1_SES_* and *PC2_SES_*) as aggregate indicators of SES because these PCs were sufficient to explain the overall SES.

Fig 1B and S2 clearly demonstrate that the first two PCs are both necessary and sufficient to reasonably differentiate major SES groups. The later PCs no longer appear to contribute to distinguishing different SES levels. *PC1*_SES_ mainly distinguishes high and low SES groups and appears to reflect the positive correlation among different SES measures. *PC1*_SES_ mostly loads individual differences in occupation, educational attainment, and income (Fig 1A). On the other hand, *PC2*_SES_ contributes more to explaining the residual variation in the lower SES groups and illustrates more subtle aspects of SES. *PC2*_SES_ primarily reflects occupation and neighborhood qualities that are not strongly linked with educational attainment or income (Fig 1A and S2 and Table S2).

Furthermore, the PCA results reveal the complex nature of SES within lower SES groups. Fig 1B shows that the lower SES a group represents, the more positively correlated the two components are. While the highest SES group even has a lower mean value of *PC2*_SES_ compared to that of the lower SES groups, relatively better-off individuals within the lower SES groups tend to have higher levels of both *PC1*_SES_ and *PC2*_SES_. These observations imply that the dimensions of SES are more complex in the lower SES groups. Overall, these results demonstrate the multidimensional nature of SES, which cannot be sufficiently described by a single SES measure.

Fig S3 plots the eigenvalues of the extracted PCs. *PC1_SES_* represents a dominantly large part of the variation from our SES measures (eigenvalue=2.77). The eigenvalues decrease substantially from *PC2_SES_*, which nonetheless explains an important amount of variation (eigenvalue=1.44). While the eigenvalues of the third and fourth PCs are not very different from that of *PC2_SES_*, these PCs do not appear to explain the meaningful variation in SES as shown earlier.

Prior to the analyses, we standardized *PC1_SES_* and *PC2_SES_* so that they have zero mean and unit variance.

#### 3.3. Control variables

We used the following variables as baseline control variables.

- Age at brain scan (linear, squared, and cubed terms) - field 21003
- Sex - field 31
- Age (linear, squared, and cubed terms) ⨯ Sex
- Total intracranial volume - estimated from CAT12
- Site of acquisition (Cheadle, Reading, or Newcastle) - field 54
- A natural cubic spline function of acquisition date (number of days when the acquisition happened since the acquisitions started) with 3 degrees of freedom^6^ - field 53
- Time of test (in seconds) - field 21862
- Interaction terms of acquisition site with all of the above
- The first 40 PCs of the genetic data - field 22009
- Genotyping array (UK BiLEVE or UK Biobank Axiom array) - field 22000

The acquisition date and time were included as control variables based on a recent paper (*48*), indicating that these variables account for subtle differences in the UKB’s assessment protocols over time. For instance, Fig S1 demonstrates that there is a subtle temporal pattern over time since the UKB started collecting MRI images. We used a natural cubic spline function in order to capture highly non-linear patterns flexibly. The first 40 genetic PCs were also used to control for the genetic population structure and the genotyping array to control for potential confounds in the genetic PCs due to different arrays used.

#### 3.4. Genome-wide association studies and construction of the polygenic index for SES

As a measure of genetic variation associated with SES, we used a polygenic index (PGI) that additively summarizes the effects of more than 1 million genetic markers. The genetic markers used here are single nucleotide polymorphisms (SNP), which are the most common form of genetic variation. A PGI *s*_i_ of individual *i* is a weighted sum of SNPs:

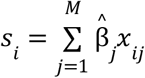

where *x* _*ij*_ represents the genotype of individual *i* for SNP *j* coded as the count of the reference allele. We estimated the weights 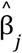 from genome-wide association studies (GWAS), which conduct univariate regressions of an outcome on each SNP across the genome. The resulting estimates were then adjusted for the correlation between the SNPs to obtain the weights 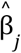.

We constructed a PGI for SES (*PGI_SES_*) by combining multiple GWAS results of SES indicators, which included educational attainment, occupational wages, household income, local average income, and neighborhood score (see further details below). We conducted GWAS on each of these measures with the UKB participants of European ancestry, excluding those in the analysis sample of this study as well as their close relatives (up to the third degree of relatedness^7^). We ran each GWAS with a linear mixed model, estimated by BOLT-LMM (*49*).

*Educational attainment* (years of schooling) was coded in the same way as the recent large-scale GWAS (*21*). *Household income* was coded as the natural log of the midpoint income of each income bracket.^8^ The remaining indicators were the same as described in Section 3.2.1. Except for educational attainment, GWAS was run on male and female samples separately and the male and female results of each measure were meta-analyzed by the meta-analysis version of MTAG (*50*) to account for possible sex heterogeneity in socio-economic outcomes.^9^ The sample sizes for each measure varied from 250,865 to 401,026 participants. For educational attainment, the GWAS result was meta-analyzed with the existing GWAS meta-analysis result of educational attainment (*21*), which excludes the UKB. More details of these GWAS are summarized in Table S4.

Finally, we combined these GWAS results to represent general SES by the common-factor GWAS function of Genomic SEM (*34*). The effective sample size of this common-factor SES GWAS amounts to 849,744 (*51*). We then constructed the PGI for SES for those of European ancestry in the analysis sample (*N* = 20,799). To adjust for the correlation between the SNPs, we used a Bayesian approach called LDpred (*52, 53*) with a reference panel from the Haplotype Reference Consortium (version 1.1) (*54*). The SNPs included in the *PGI_SES_* were limited to the autosomal bi-allelic SNPs established by the International HapMap 3 Consortium (*55*), which are known to work well for phenotype predictions (*21, 56*). The SNPs were also filtered to ensure minor allele frequency > 0.01, the imputation score (INFO) > 0.7, and the missing rate < 0.05. As a result, 1,020,632 SNPs were used for *PGI_SES_*. The *PGI_SES_* was standardized to have zero mean and unit variance.

*PGI_SES_* predicts about *ΔR^2^*=7.1% of the variation of *PC1*_SES_ out-of-sample among individuals of European ancestries, above and beyond the control variables (age, age^2^, age^3^, sex, interactions between sex and the age terms, genotyping array, and the first 40 genetic PCs). On the contrary, *PGI_SES_* barely predicts *PC2*_SES_, explaining *ΔR^2^*=0.02% of its variation.

### 4. Statistical analyses

#### 4.1. Voxel-based Morphometry (VBM) analysis

##### 4.1.1. Baseline analysis

Our baseline analysis estimated the associations between voxel-level GMV and the two SES PCs. For each voxel *j*, we estimated the following regression model via ordinary least square (OLS):

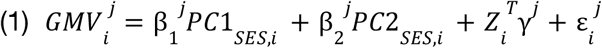

where the GMV of voxel *j* is regressed on the two SES PCs. The vector *Z_i_* include the control variables listed in Section 3.3. 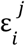 is the error term. The GMV and the SES PCs were standardized to have zero mean and unit variance. An *F*-test was used for each voxel to test whether there is significant association between voxel *j*’s GMV and the SES PCs jointly with the the null hypothesis 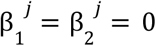. We measured the association size by the variance of interest in GMV explained by the SES PCs beyond the covariates of no interest, *i.e.*, partial 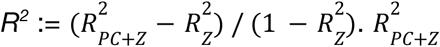 is the *R^2^* from the unrestricted model, which includes the two SES PCs and the covariates of no interest, and 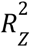 is the *R^2^* from the restricted model, which only includes the covariates of no interest. We also quantified the relative contribution of *PC1*_SES_ in the overall association size by 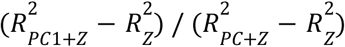. We used permutation testing to correct for multiple hypothesis testing across voxels (see Section 4.1.3 for details).

After the estimation, we anatomically labeled the voxels using the Neuromorphometrics atlas provided in CAT12. For a summary purpose, we also generated cluster-based estimates. Each cluster consists of at least 200 neighboring voxels within the lobe^10^ which are significant at the familywise error rate of 5% in the baseline model. We then repeated the same analysis with mean GMV of these clusters.

##### 4.1.2. Controlling for total intracranial volume (TIV)

Analyses that aim to identify associations between localized GMV and outcomes typically control for TIV, since volumetric brian measures scale with the head size. However, controlling for TIV as a linear covariate has important statistical implications for identifying localized GMV patterns linked to SES, because TIV is positively correlated with both SES and regional GMV. In Fig 2B, GMV of some voxels appear to have negative association with SES when the TIV is included as a control variable in the model. On the contrary, Fig 2A shows that almost all the voxel-level GMV are positively associated with SES when TIV is not controlled for. This result indicates that the absolute GMV-SES association is unlikely to be negative in any brain region.

To formally illustrate this point, consider a VBM model for SES with only the TIV as a covariate without loss of generality:

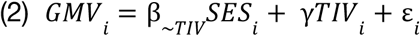

Where *GMV_i_* is the GMV of some voxel and β_∼*TIV*_ denotes the association between the voxel’s GMV and SES while TIV is accounted for. Each variable is standardized to have zero mean and unit variance without loss of generality. γ corresponds to the association between the GMV and the TIV, conditional on SES. The linear dependence between the TIV and SES can be described as: *E*[*TIV*_i_ |*SES*_i_] = λ*SES*_i_. If we denote β as the coefficient of SES from the regression of the GMV on SES without the TIV as a covariate, β_∼*TIV*_ can be written as:

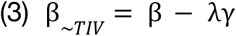

Therefore, if both λ and γ are positive and large, β_∼*TIV*_ can be negative even when β is positive. Our data suggests that this is indeed the case: With the baseline model^11^, we estimated λ̂ = 0. 10 for *PC1_SES_* and λ̂ = 0. 01 for *PC2*_SES_. γ̂ was on average 0.46 with the minimum=0.11 (right exterior cerebellum) and the maximum=0.72 (left gyrus rectus). Since estimates of β are positive for the vast majority of the voxels, one cannot conclude that the absolute GMV-SES association is truly negative even when estimates of β_∼*TIV*_ are negative. Instead, such negative estimates are evidence that λγ is large relative to β and that the GMV-SES association is essentially very small or non-existent for these regions.

Therefore, caution is warranted when interpreting the results when TIV was adjusted for as a covariate. For this reason, we reported the VBM results both with or without TIV included as a covariate. Furthermore, given the above, our results suggest that SES is associated with greater gray matter across almost all brain regions investigated, despite small exceptions with negative estimates after adjusting for the TIV. Note that TIV was always included as a control variable unless otherwise stated.

##### 4.1.3. Multiple testing correction

To correct for multiple testing across voxels, we used permutation testing to determine a *p*-value threshold that controls the familywise error (FWE) rate of 5% (*57*). Following a comprehensive simulation study that examined several permutation approaches for the brain-imaging (*58*), we applied the method developed by Freedman and Lane (1983) to construct an empirical distribution of test statistics (*59*). Consider an *N* × *M* matrix *Y* where column *j* is a length-*N* vector of voxel *j*’s GMV with *M* the number of the voxels. Each column was first residualized of the covariates of no interest (*Z*_i_). Matrix *Y* was then permuted row-wise so that the correlation structure among the voxels was preserved. We then regressed each of the permuted GMV on the non-permuted, original regressors and recorded the maximum *F*-statistic. We repeated this process 5,000 times to form a distribution of the maximum *F*-statistics. We used the *p*-value computed from the 95th percentile of this distribution (*F* = 13.04) as the *p*-value threshold for 5% FWE-corrected significance level, which corresponds to *p* = 2. 193 × 10^-6^ (uncorrected).

While in principle the permutation testing has to be performed for each different analysis, the resulting *p*-value thresholds differed only marginally and the threshold for the baseline model was the most conservative. Therefore, we used the 2. 193 × 10^-6^ threshold for every voxel-based analysis.

##### 4.1.4. Stratified analysis of high and low SES groups

To investigate potential heterogeneity across different SES groups, we conducted the same VBM analysis separately on high and low SES groups. High and low SES groups were defined by National Statistics Socio-economic Classification of the UK (*32*): high SES group holds a managerial, administrative, or professional occupation and low SES group holds intermediate, routine, or manual occupation (*N_high_* = 15,611, *N_low_* = 8,320).

##### 4.1.5. VBM of Individual SES measures

To gain additional insight into the neuroanatomical correlates of SES, we conducted additional VBM analyses on each of the five individual numerical SES measures used to construct the SES PGI, described in Section 3.4. Note that the main purpose of these analyses was not to discover novel neuroanatomical correlates from each SES measure, but rather to compare neuroanatomical correlates across these measures.

#### 4.2. Estimating the overall association between SES and GMV structure

Our VBM results demonstrate that the association between SES and an individual GMV IDP is small and does not exceed partial *R*^2^ of 1% with TIV adjusted for. One might then ask how large the brainwide association between SES and the gray matter structure is if we can aggregate individual SES-GMV association estimates from individual voxels. Estimating the overall association is not an easy task because of the high dimension of the voxel-level GMV data and the strong spatial correlation among the voxels. We addressed these challenges by constructing a brainwide GMV score for SES with a machine learning technique. We used a stacked block ridge regression approach inspired by a recent whole-genome regression method (*33*). This approach allows us to tackle the high dimension issue by stacked regressions and the spatial correlation by the use of ridge regressions without excessive computational burden. Ridge regressions also ensure that we only capture linear relationships between SES and the GMV structure.

We constructed a brainwide GMV score for each SES PC in two steps:

1. Voxels were first partitioned into blocks of 10,000 adjacent voxels. For each block, we ran a ridge regression of each SES PC on its 10,000 voxel-level GMVs with arbitrarily-chosen varying shrinkage parameters: {100, 100^2^, 100^3^}. We then computed predictions for each SES PC for each value of the shrinkage parameters, resulting in 3 predictors for each SES PC from each block. This resulted in 153 predictors from 51 blocks partitioned from 504,426 voxels.
2. After collecting the predictors from all the blocks, a ridge regression was run on them together again. The prediction from this regression was used as a brainwide GMV score.

Both steps were implemented in 5-fold nested cross-validation: In the outer loop, the sample was split into 20% test set and 80% training set, the latter of which was again split into 20% validation set and 80% training set in the inner loop. In the inner loop, cross-validation was used to tune the shrinkage parameter for the step-2 ridge regression. The outer loop was used to train the final model and obtain predictions for the test set given the obtained value of the shrinkage parameter from the inner loop. We ensured that no information from the test set was used in the model training.

To measure the overall association between each SES PC and the GMV structure, we used a change in *R^2^* after including the corresponding brianwide GMV score to the regression. The covariates used were age, age^2^, age^3^, sex, interactions between sex and the age terms, TIV, genotyping array, and the top 40 genetic principal components. We computed confidence intervals with 1,000 bootstrapped samples.

Of note, we do not claim here that this approach is the best way of constructing a brainwide score or estimating the brainwide association. The primary goal of this analysis is to demonstrate that SES is associated with GMV structure to a substantial degree.

#### 4.3. Incorporating genetics

##### 4.3.1. VBM with PGI

Using *PGI_SES_*, we conducted the following additional VBM analyses: (1) VBM of SES PCs only with individuals of European Ancestry (2) VBM of *PGI_SES_* (3) VBM of the SES PCs controlling for *PGI_SES_*. These VBMs were carried out in the same way as the baseline analysis detailed in Section 4.1. We then examined which GMV voxels are significantly associated with the SES PCs and/or the PGI and examined changes in SES-GMV associations before and after the PGI was controlled for. Note that we measured partial *R^2^* of the PCs for VBM of the SES PCs controlling for *PGI_SES_* as 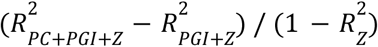 to be able to compare it with partial *R^2^* from VBM of SES PCs. In addition to probing the difference in statistical significance after the PGI was controlled for, we directly tested whether controlling for the PGI significantly altered the SES-GMV association.

##### 4.3.2. Testing differences in SES-GMV associations with and without PGI as a control variable

We used a Wald test to examine whether there was a significant difference in the SES-GMV association before and after the *PGI_SES_* was controlled for. More specifically, consider a model where *PGI_SES_* is added to the model (1) and also set up an auxiliary regression of the PGI on the SES PCs and the covariates for each voxel:

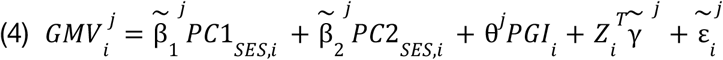

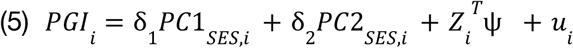

Using vector notations: 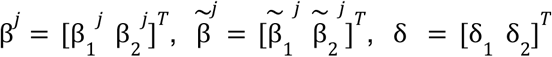, which are all length-2 vectors, it can be shown:

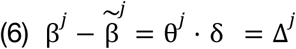

Therefore, the vector Δ*^j^* represents the difference in the SES-GMV association for voxel *j* due to controlling for *PGI_SES_*. Δ^*j*^ can be estimated as the product of estimates of 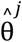 and 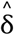 from the model (4) and (5), respectively. A Wald test was then used to test the null Δ ^j^= 0 with the test statistic: 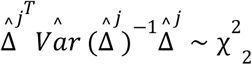, where 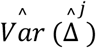 was approximated by the delta method: 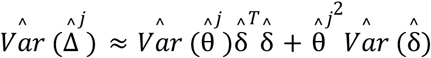. Note that this analysis is statistically equivalent to a mediation analysis with *PGI*_SES_ being a mediator (*60*). We conducted this test only for the voxels whose GMV was significantly associated with the PCs. Then, the multiple testing was corrected for using Bonferroni correction (the corrected 5% threshold = 1. 46 × 10^-6^ with 34,188 tests).

##### 4.3.3. Measuring differences in SES-GMV associations with and without PGI as a control variable

To represent the relative size of Δ^j^ in relation to partial *R^2^*, we used the relative change in the net variation explained by the SES PCs after adding *PGI*_SES_ to the model with the covariates of no interest: 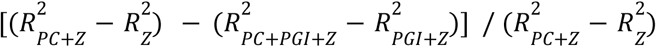. This measure is bounded between 0 and 1 as long as the sign of the coefficients for *PC1*_SES_ and *PC2*_SES_ do not change after controlling for *PGI*_SES_. This expression can be interpreted as the percent change in the SES-GMV associations due to controlling for *PGI*_SES_ and essentially the part of the SES-GMV association that can be attributed to *PGI*_SES_. Note that, because *PC2*_SES_ is barely predicted by *PGI_SES_* and even barely heritable (Table S5), the percent change in SES-GMV association after controlling for *PGI*_SES_ is essentially due to the change in *PC1*_SES_-GMV association. We can therefore rewrite the earlier expression as:

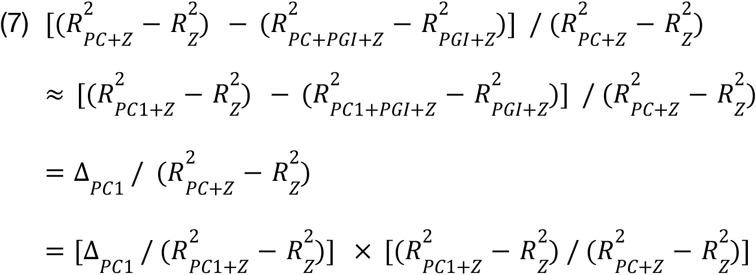

where 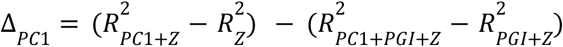, the change in the net variance explained by *PC1*_SES_ after controlling for *PGI_SES_*. Hence, the percent change in SES-GMV association is roughly the product of the percent change in *PC1*_SES_-GMV association 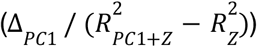 and the relative contribution of *PC1_SES_* in the overall SES-GMV association 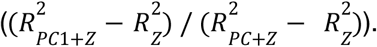 A larger share of SES-GMV association can be attributed to *PGI_SES_* if genetic factors linked to SES play a bigger role for *PC1*_SES_-GMV association and/or if *PC2*_SES_ contributes relatively less to the overall SES-GMV association.

##### 4.3.4. Measurement error correction for PGI

*PGI*_SES_ is a noisy proxy of true linear effects of common genetic variants that are linked to SES because GWAS estimates of individual SNP effects are obtained from finite sample sizes. The difference between the true PGI and the available PGI can be viewed as the classic measurement error, which leads to an attenuation bias in the coefficient estimate for the *PGI*_SES_. Nonetheless, it is still possible to account for the linear effects of common genetic variants that the true *PGI*_SES_ would capture under reasonable assumptions. We addressed this attenuation bias by using genetic instrumental variable (GIV) regression (*35*). The essential idea is that the true *PGI*_SES_ can be recovered from a noisy *PGI*_SES_(^1^) by using another *PGI*_SES_^(2)^ as an instrumental variable that was derived from a different GWAS sample. The crucial assumption here is that the noise in *PGI*_SES_^(1)^ and *PGI*_SES_^(2)^ is uncorrelated to each other. GIV regression can address the measurement error in *PGI*_SES_ to the extent that this assumption holds.

To obtain *PGI*_SES_^(1)^ and *PGI*_SES_^(2)^, we randomly split the UKB GWAS sample into two subsamples (*N*=105,517∼170,945) such that each subsample has the same male-female ratio and no individuals in one subsample are related to anyone in the other subsample with more than the third degree of relatedness. With each subsample, GWAS was run for the five numerical SES measures and the results were combined with Genomic SEM as described in Section 3.4. Then, *PGI*_SES_^(1)^ and *PGI*_SES_^(2)^ were constructed from one of the two independent GWAS subsample results in the main imaging sample.

Using *PGI*_SES_^(1)^ and *PGI*_SES_^(2)^, we fitted the model (4) by the GIV estimation, which is two-stage least squares (TSLS).

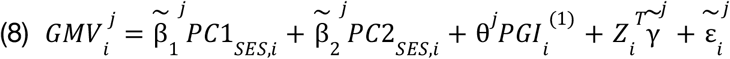

Where *PGI_i_*^(1)^ is the PGI estimated from the first subsample. The first-stage equation can be written as:

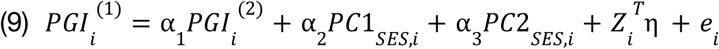

where the PGI estimated from the second subsample, *PGI_i_* ^(2)^, is used as an instrument for *PGI*_i_^(1)^. We obtained the TSLS estimates by fitting the following equation:

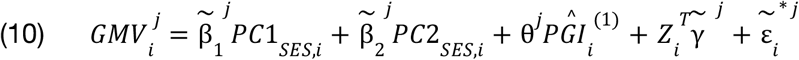

where 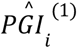 is the fitted value from the equation (8). The statistical inference was then conducted but in the standard TSLS framework to test the association between the GMV and SES for each voxel conditional on *PGI*_SES_ (*61*).

We computed Partial *R^2^*’s based on adding or excluding 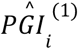 in model (10) instead of the unadjusted PGI. Similarly, we measured the difference in SES-GMV association after controlling for the PGI by GIV as 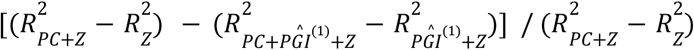

#### 4.4. Functional annotations

We connected our anatomical findings to known functional localizations by leveraging Cognitive Atlas and the extrapolatable meta-analysis tool NeuroQuery (*36, 37*). We first took the 518 cognitive concepts from Cognitive Atlas which were categorized into 11 functional categories.^12^ Then, for each concept, we generated a meta-analyzed Z-score brain map using NeuroQuery. This toolbox allows users to generate a predictive MRI-derived spatial distribution for any term, based on very large-scale meta-analyses containing mostly functional MRI studies. We excluded 12 concepts containing a term for which NeuroQuery failed to generate a brain map. As a result, 506 concepts remained. For each concept-associated brain-map, we calculated mean χ^2^ with voxels statistically significant at the FWE rate of 5% level in the VBM results. We used these mean χ^2^ scores as a summary measure of associations between a given functional concept and the regions linked to SES. The full results are reported in Table S17.

### 5. Interpretation

#### 5.1. Brain, SES, and genetics

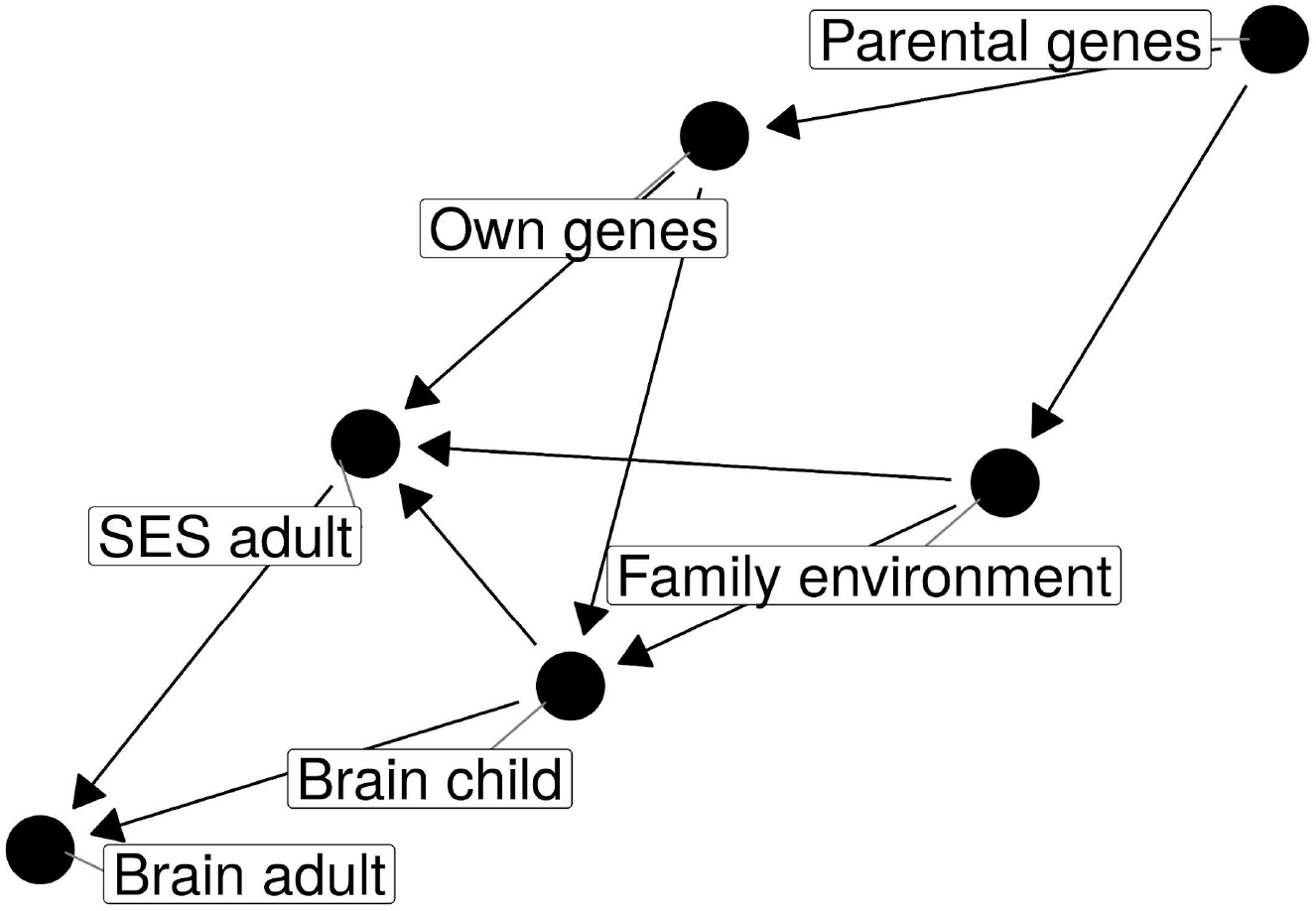

To aid interpretation of the association between SES and brain anatomy observed in late adulthood, the figure above describes a simple model that illustrates how adulthood brain anatomy can be linked to SES, family environments, and genetics. The model is depicted in a directed acyclic graph (DAG), a popular graphical framework for identifying confounding variables (*62–64*). The model does not attempt to include all possibly relevant factors and mediating pathways. Rather, its purpose is to identify what effects are potentially captured in the estimated GMV-SES association in relation to genetics and family environments.

It is important to note that each arrow in the DAG represents a unidirectional causal relationship between two variables (nodes). For instance, the arrow from “*SES adult”* to “*Brain adult”* only indicates the environmental effect of adult SES on the adult brain. A path is a set of one or more arrows that connects multiple nodes. A path can be either open or closed. An open path channels statistical associations, which can be closed by conditioning on a variable in the middle. A path can be closed due to a collider, which is a variable that receives two arrows. Conditioning on a collider opens up a closed path, which induces a collider bias.

Though fairly simple, the model is capable of describing key relevant pathways. First, child brain development is determined by genetics and family environments (“*Own genes* → *Brain child”* and “*Family environment* → *Brain child”*). Second, SES in adulthood is a function of genetics, family environments, and child brain development (“*Own genes* → *SES adult”,* “*Family environment* → *SES adult”,* and *“Brain child* → *SES adult”*). Third, the transition to the (late) adulthood brain is partly influenced by adult SES (“*Brain child* → *SES adult* → *Brain adult”* and *“Brain child* → *Brain adult”*). Therefore, the model describes the roles of both genetics and family environments in causing differences in SES and the brain. Furthermore, the feedback between SES and the brain is illustrated by the path: “*Brain child* → *SES adult* → *Brain adult”.* One could extend the model by distinguishing late and early adulthood phases and including another feedback effect. Such an extension, however, will not provide additional key insights as long as socioeconomic mobility is limited during adulthood.

Another important feature is that the model recognizes so-called genetic nurture effects (*65*). Childhood family environments shaped by the parents are known to be associated with the genes of the parents (“*Parental genes* → *family environment”*), which are passed on to their child (“*Parental genes* → *Own genes”)*. These links induce statistical associations between own genetics and family environments (“*Own genes ← Parental Genes* → *family environment”*). This fact statistically blurs the common dichotomy between genetics and family environments.

In this study, we regressed voxel-level GMV on an adult SES measure with a goal to estimate the SES-GMV association. If our aim were to estimate the causal effect of adult SES environments on the GMV structure (*i.e*., “*SES adult* → *Brain adult”*), a resulting regression estimate will clearly be biased due to the open confounding paths, which transmit statistical associations. Therefore, the estimated SES-GMV associations in this study are expected to encompass the direct environmental effect of adult SES on adult brain and all the effects due to the open paths, which can be summarized as follows:

1. Environmental effects of adult SES on adult brain: *SES adult* → *Brain adult*
2. Brain causing SES: *SES adult ← Brain child → Brain adult*
3. Genetic effects: *SES adult ← Own genes → Brain child → Brain adult*
4. Family environment effects: *SES adult ← Family environment → Brain child → Brain adult*
5. Genetic nurture effects on brain*: SES adult ← Own genes ← Parental genes → Family environment → Brain child → Brain adult*
6. Genetic nurture effects on SES: *SES adult ← Family environment ← Parental genes → Own genes → Brain child → Brain adult*

Notably, the DAG above demonstrates that one needs to account for either childhood brain measures (*i.e.*, lifetime longitudinal data) or measures of both family environments and genetics in order to identify the causal effect of the adult SES on the brain (“*SES adult* → *Brain adult”)*, assuming the absence of no other unobserved confounders.

#### 5.2. Interpretation of the polygenic index for SES

While statistical analysis using *PGI*_SES_ is straightforward, careful interpretations are required. Most importantly, the remaining associations between the GMV and SES after conditioning on *PGI_SES_* cannot entirely be interpreted as environmental effects of SES on the brain anatomy because *PGI_SES_* only captures noisy estimates of the effects of measured common genetic variants. It does not include the potential effects of structural or rare genetic variants that are not (or only partly) captured by the observed common genetic variants. Nonetheless, the GMV-SES association that is robust to controlling for *PGI_SES_* can point to regions of the brain that are more likely to be affected by environmental factors linked with SES.

To interpret the results, we first need to probe what effects are likely to be summarized in *PGI_SES_.* On the basis of the DAG presented above, the GWAS of SES will capture the direct genetic effects on SES (“*Own genes → Brain child → SES adult”* and “*Own genes → SES adult”*) as well as the effects due to confounders, namely genetic nurture effects (“*Own genes ← Parental genes → Family environment → SES adult”* and *“Own genes ← Parental genes → Family environment→ Brain child → SES adult”*). All of these effects will therefore be incorporated in *PGI_SES_.* Furthermore, it is important to note that the paths via the adult brain will not be captured in *PGI_SES_* due to the adult brain being a collider: “*Own genes → Brain child → Brain adult ← SES adult”*.

These observations lead to the following interpretations for the SES-GMV association estimates conditional on *PGI_SES_*. First and most importantly, *PGI_SES_* is expected to capture a part of the SES-GMV association due to different family environments and parental SES. A PGI captures the association between a phenotype and genetic variants, rather than causal effects of genetic variants. For this reason, *PGI_SES_* will contain genetic nurture effects as described above. Studies have shown that such genetic nurture effects tend to be larger for socio-economic phenotypes (*65, 66*). Therefore, *PGI_SES_* is likely to overstate the genetic effects associated with SES.

Second, what we effectively control for by controlling for *PGI_SES_* is the shared genetic architecture between SES and developmental neuroanatomy that is captured by the measured genetic variants and their estimated linear associations with SES. Hence, controlling for *PGI_SES_* is not necessarily equivalent to controlling for the entire common genetic variants behind the GMV-SES association. More specifically, in light of the DAG, *PGI_SES_* will account for the following genetic effects on SES: “*Own genes → SES adult”* and *“Own genes → Brain child → SES adult”,* the latter of which works via the child brain. On the other hand*, PGI_SES_* will not account for the genetic effects on the adult brain that do not work through adult SES: “*Own genes → Brain child → Brain adult*”. In fact, in order to account for the underlying genetic effects in the SES-GMV association, it would be required to construct a *PGI* for a brain IDP conditional on adult SES. However, it is currently difficult to construct such a *PGI* with sufficient predictive power due to a limited sample size available for conducting a required GWAS. Moreover, such a *PGI* will need to be constructed for each IDP representing a sufficiently narrow region.

Despite these challenges for interpretation, *PGI_SES_* is still useful for identifying brain regions likely to be more susceptible to the influence of socio-economic environments than that of genetic factors. If the estimated SES-GMV association is relatively less attenuated after controlling for *PGI_SES_*, the observed SES-GMV association is likely to be a result of environmental effects of SES rather than genetic factors. One reason is because *PGI_SES_* tends to overestimate the effects of common genetic variants on SES. Also, at least for healthy individuals, it is highly unlikely that the SES-GMV association is dominantly driven by rare or structural genetic variants with only negligible contribution from common genetic variants associated with SES.

### 6. Supplementary analyses

#### 6.1. Heritability and genetic correlation

We estimated SNP-based heritability of SES, TIV, and the brainwide GMV scores as well as their pairwise genetic correlation, using genomic-relatedness-based restricted maximum likelihood (GREML) estimation (*67, 68*). The method estimates the genetic contribution to the phenotypic variance based on a linear mixed model, where the genetic effects are modelled as random. Its extension to a bivariate model estimates genetic correlation between two phenotypes.

We randomly dropped one of a pair of individuals with estimated relatedness greater than 0.05, which resulted in *N* = 20,447 (*69*). We used a slightly pruned set of the SNPs used to construct *PGI*_SES_ with the following pruning parameters: window size = 1,000 variant counts, step size = 5, *r*^2^ = 0.95. As a result, 452,190 SNPs were included. As covariates, we included age, age^2^, age^3^, sex, interaction terms between the sex and age terms, genotyping array indicator, and top 40 genetic PCs. The estimation was implemented in BOLT-REML (*70*).

The results are reported in Table S5. TIV and the GMV score for *PC1_SES_* were both partly heritable (*h^2^* = 0.41, *SE* = 0.02; and *h^2^* = 0.28, *SE* = 0.02, respectively). *PC1_SES_* was moderately heritable (*h^2^* = 0.16, *SE* = 0.02) and positively genetically correlated with TIV (r_g_ = 0.37, SE = 0.06). Furthermore, *PC1_SES_* had a moderate genetic correlation with the values of the brainwide GMV score that we constructed for *PC1_SES_* (*r_g_* = 0.57, *SE* = 0.06). Similar estimates for *PC2_SES_* and GMV score for *PC2_SES_* were either smaller or had much larger standard errors (*h^2^* = 0.05, *SE* = 0.02 for *PC2_SES_*; *h^2^* = 0.14, *SE* = 0.02 for GMV score; *r_g_* = 0.18, *SE* = 0.10 with TIV; *r_g_* = 0.34, *SE* = 0.18 with the GMV score). Overall, these results demonstrate that the genetic architectures of SES and brain structure are partly overlapping.

#### 6.2. Testing differences in residual SES-GMV associations due to BMI

As presented in Figure 3C and 3D, the remaining SES-GMV associations after controlling for *PGI_SES_* can be substantially attributed to individual differences in BMI. Here we formally tested whether this is the case statistically. In other words, we tested whether there is a statistically significant change in at least one of the coefficients for *PC1_SES_* and *PC2_SES_* after accounting for BMI in addition to *PGI_SES_*. The testing procedure was analogous to the one conducted for *PGI_SES_*, which is described in Section 4.3.2, except that GIV regression was used to estimate each model. As it was done for *PGI_SES_*, we conducted this test only for the voxels that had significant association with the PCs and then the multiple testing was corrected for using Bonferroni correction. As a result, we found that 84.4% of 34,188 voxels tested had a significant change in at least one of the coefficients for *PC1_SES_* and *PC2_SES_* after controlling for BMI in addition to *PGI_SES_*. This result confirms that BMI can indeed explain the remaining SES-GMV associations after adjusting for *PGI_SES_*.

Note that, to measure the contribution of BMI in explaining the remaining SES-GMV associations after controlling for *PGI_SES_*, we again used the relative change in the net variation explained by the SES PCs. Hence, following the same logic, we computed the contribution of BMI as:

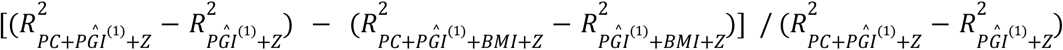

#### 6.3. Heterogeneity

Sex and age are two important factors for both SES and neuroanatomy. Therefore, we tested whether the SES-GMV associations are heterogeneous with respect to *i)* different sex (sex interaction) and *ii)* different ages (age interaction). We examined each aspect of heterogeneity separately by using the voxel clusters and including the interaction terms with *PC1_SES_* and *PC2_SES_*. The interaction terms were then tested jointly with *F*-tests.

The results are reported in Table S23-24. The SES-GMV associations were generally larger for men, with the largest difference found in the biggest cluster from the prefrontal cortex. One exception was found in a small cluster in the cerebellum, where the SES-GMV association was larger for women. However, none of the regions would survive the brainwide multiple testing correction. The SES-GMV associations also tended to increase with age, while the age interaction estimates were not large enough to be statistically significant even at the uncorrected 5% level, except for one cluster from the anterior insular and the frontal operculum. These null results for age interaction may be due to the survival effect because the majority of the participants were older than 60.

#### 6.4. Controlling for alcohol consumption

Our baseline analyses implicitly adjusted for heavy drinking by excluding heavy drinking individuals. A recent study has shown that even moderate alcohol consumption is associated with reduction in GMV even when educational attainment is adjusted for (*42*). Since alcohol drinking behavior is known to be related to SES, it may be hypothesized that the alcohol consumption is a factor that constitutes the observed SES-GMV associations. However, because individuals with high SES tend to consume a greater amount of alcohol (*71*), controlling for the alcohol consumption is expected to only increase estimates for the SES-GMV associations.

Our data confirms that this is indeed the case. In a cluster-based analysis, we controlled for the alcohol consumption (the number of drinks per week) with linear and square terms. The results show that the SES-GMV associations measured in partial *R^2^* increased by up to 31%, but only marginally in general (Table S25). Therefore, our positive estimates for the SES-GMV associations cannot be directly attributed to the alcohol consumption. Rather, when not adjusted for, the alcohol intake is a factor that reduces the GMV difference between high and low SES individuals.

### 7. Supplementary discussion

Our results quantify the extent to which SES could be an underlying cause or a confounding factor in brain imaging studies concerned with cognitive health or behavior. Furthermore, the modest effect sizes we found for specific brain regions imply that large sample sizes are required to identify robust neuroanatomical associations with SES. For example, a sample size of *N* > 1,200 or *N* > 2,800 is required for obtaining 90% statistical power to detect even the largest effect we found (partial *R^2^* = 1.3%) at *p*-values of 0.01 or 2⨉10^-6^, respectively. The effect sizes reported in this paper are far smaller than typically reported in smaller-scale studies in the past. Due to their small sample size, some of the past reports may be inflated estimates that suffer the winner’s curse (*11*).

Importantly, small effect sizes for individual voxels do not necessarily imply that the association between SES and brainwide GMV structure is also negligible. The brainwide GMV score we constructed was able to predict almost 5% of the *out-of-sample* variation of SES. The predictive accuracy of the score can be further improved with a larger training sample. Furthermore, this result suggests that the overall association between SES and GMV will be far from being modest if it is estimated in-sample.

Of note, the modest effect sizes can partly be due to the fact that the UKB participants generally hold a higher SES than the general British population (*72*). As Fig S6 suggests, SES-GMV associations are likely to be larger for lower SES individuals and therefore our results may underestimate the true SES-GMV association size for the general population. Furthermore, in order to identify robust SES-GMV association, we excluded individuals who were clinically diagnosed with a brain disease, morbidly obese, and heavy-drinking. Such sample exclusion criteria can attenuate the estimates since these traits are known or likely to be negatively associated with both SES and GMV. Fig S12 demonstrates that this was indeed the case.

---

**Fig. S1.**
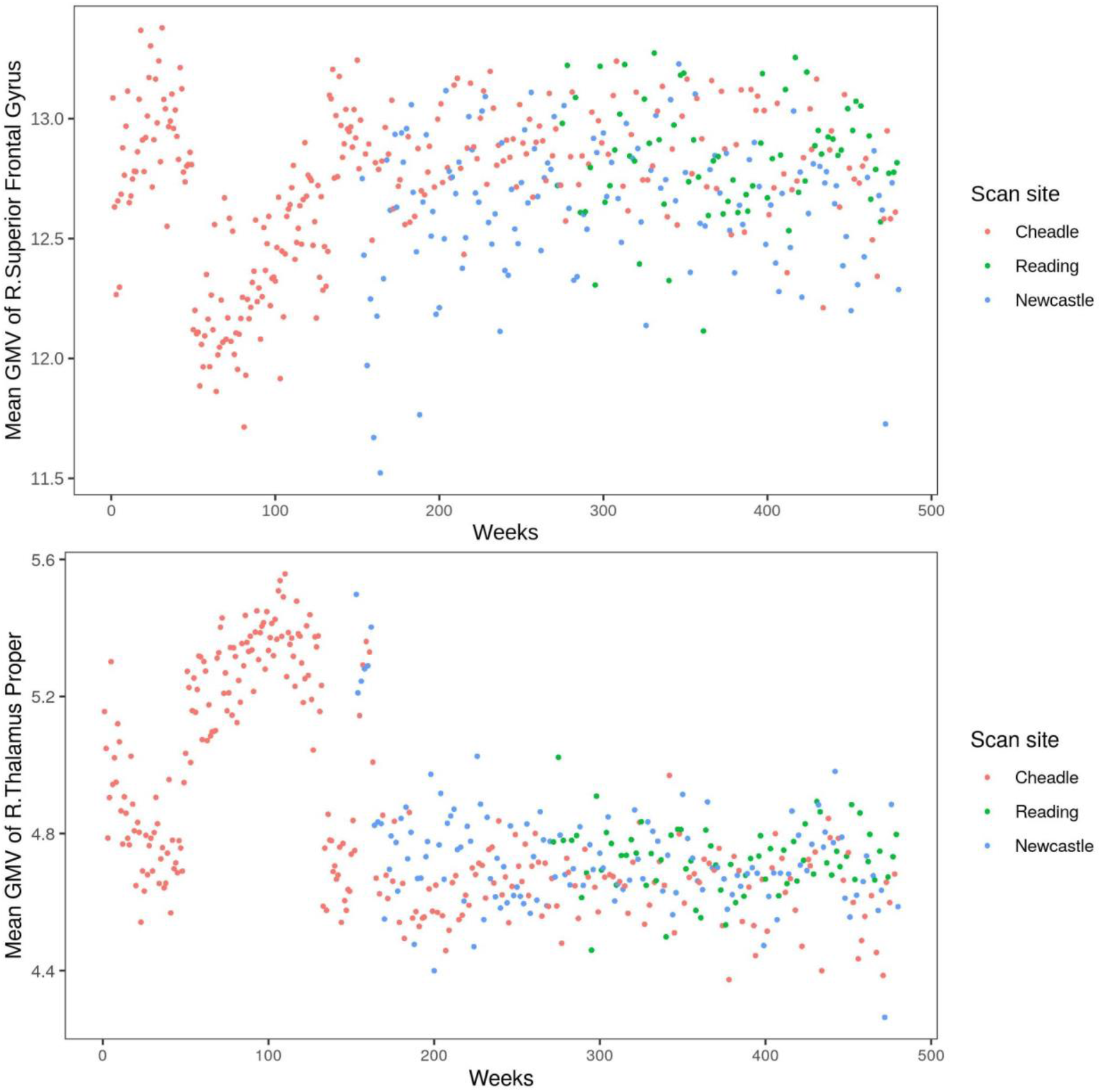
Average grey matter volume over acquisition dates by week. Site-specific weekly averages of grey matter volume are plotted over acquisition dates for the right superior frontal gyrus and the right thalamus. These regions are selected as examples and there are other regions showing similar patterns.

**Fig. S2.**
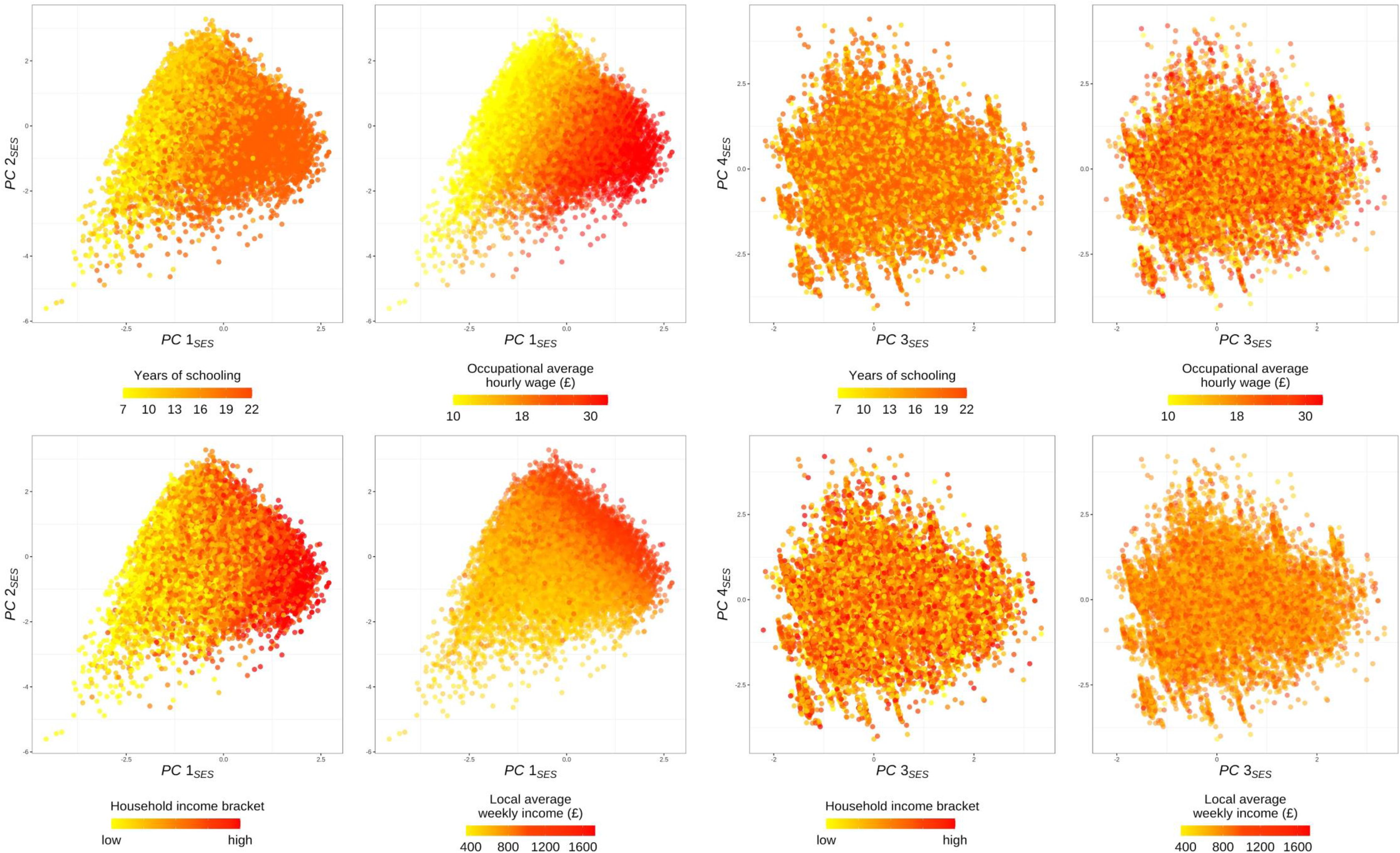
Scatter plots of top principal components (PC) for socioeconomic status (SES) Top four PCs for SES are plotted with color indicating different levels of selected socioeconomic measures.

**Fig. S3.**
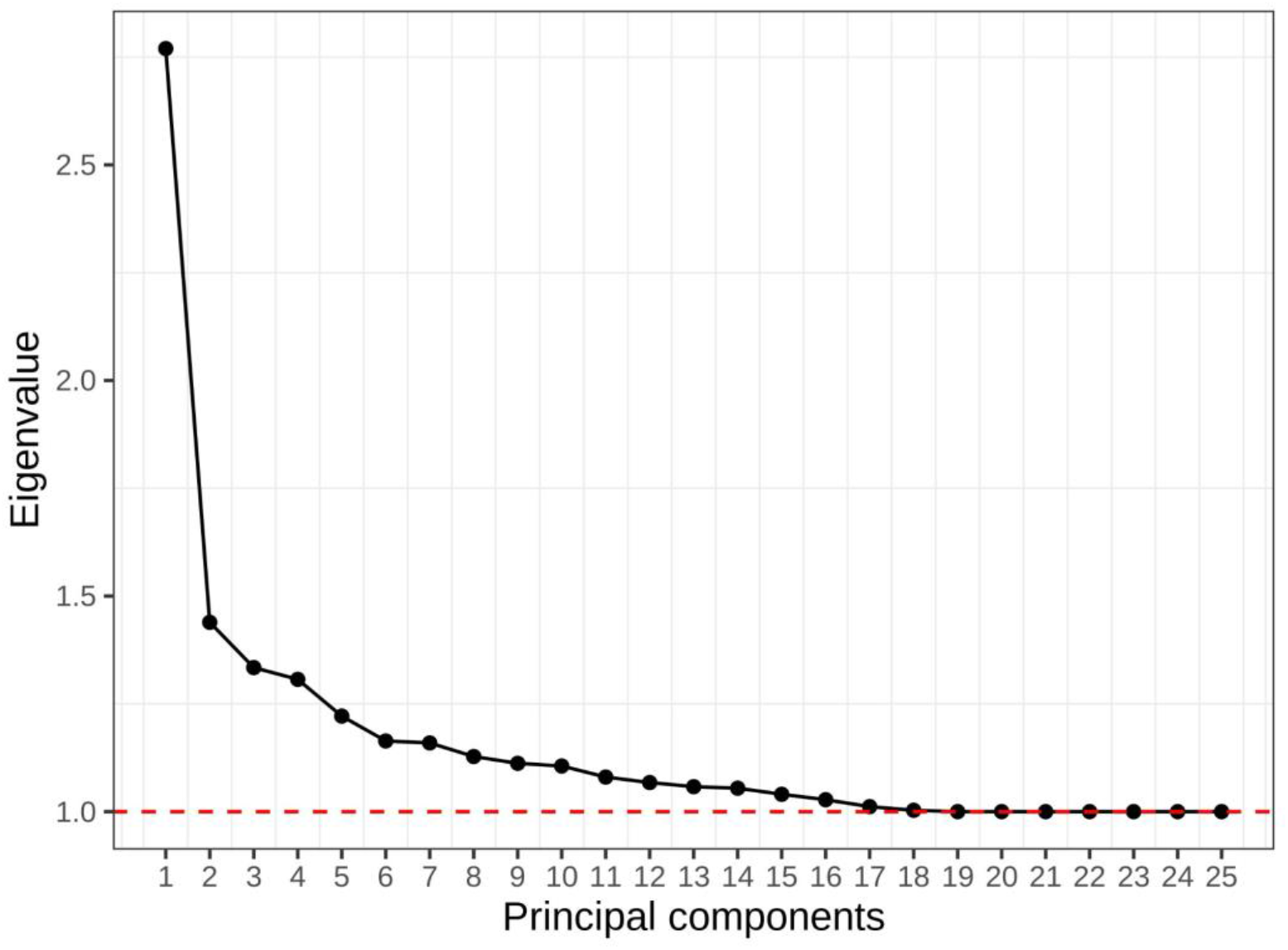
Eigenvalues from the principal component analysis of socioeconomic indicators.

**Fig. S4.**
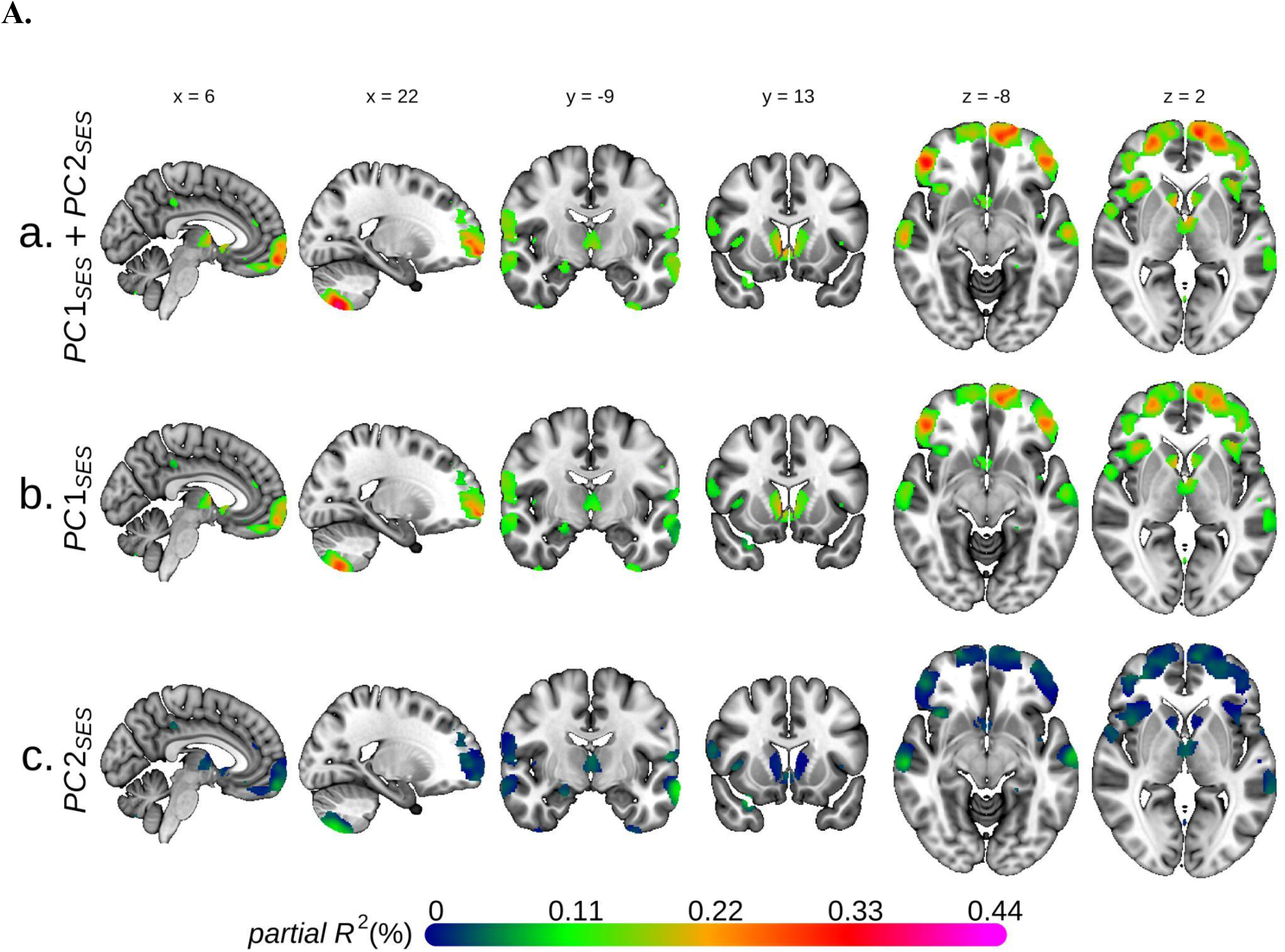
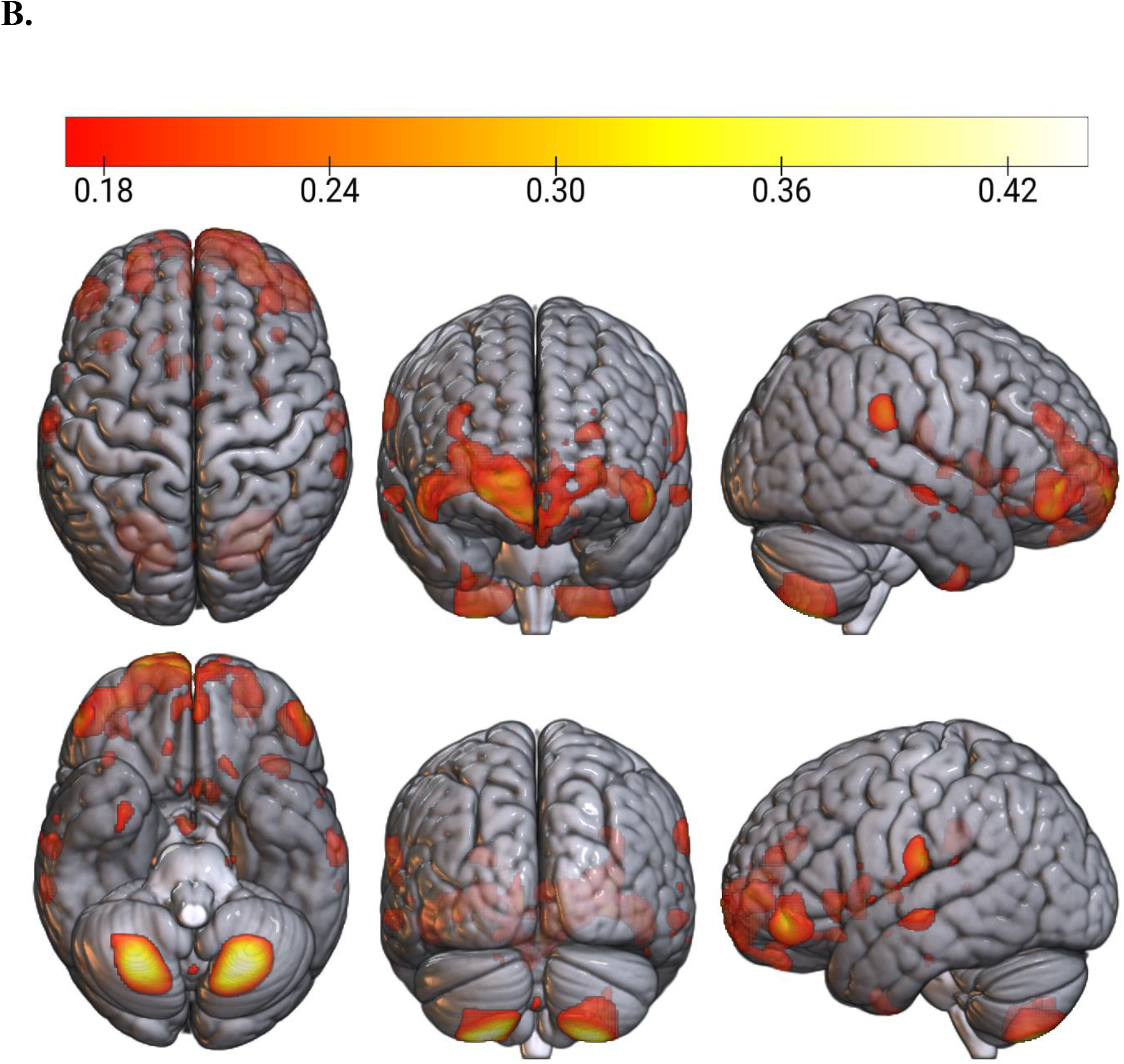
Voxel-based morphometry of grey matter volume and socioeconomic status (SES) Univariate voxel-based morphometry results on the two principal components (PC) for SES. **A.** Each figure plots the association sizes measured in partial *R^2^* for only the voxels significant at FWE rate of 5%. MNI coordinates are indicated. **B.** Partial *R^2^* (%) for two SES PC is plotted for only the voxels that were significant at FWE rate of 5% and had partial *R^2^*>0.17%.

**Fig. S5.**
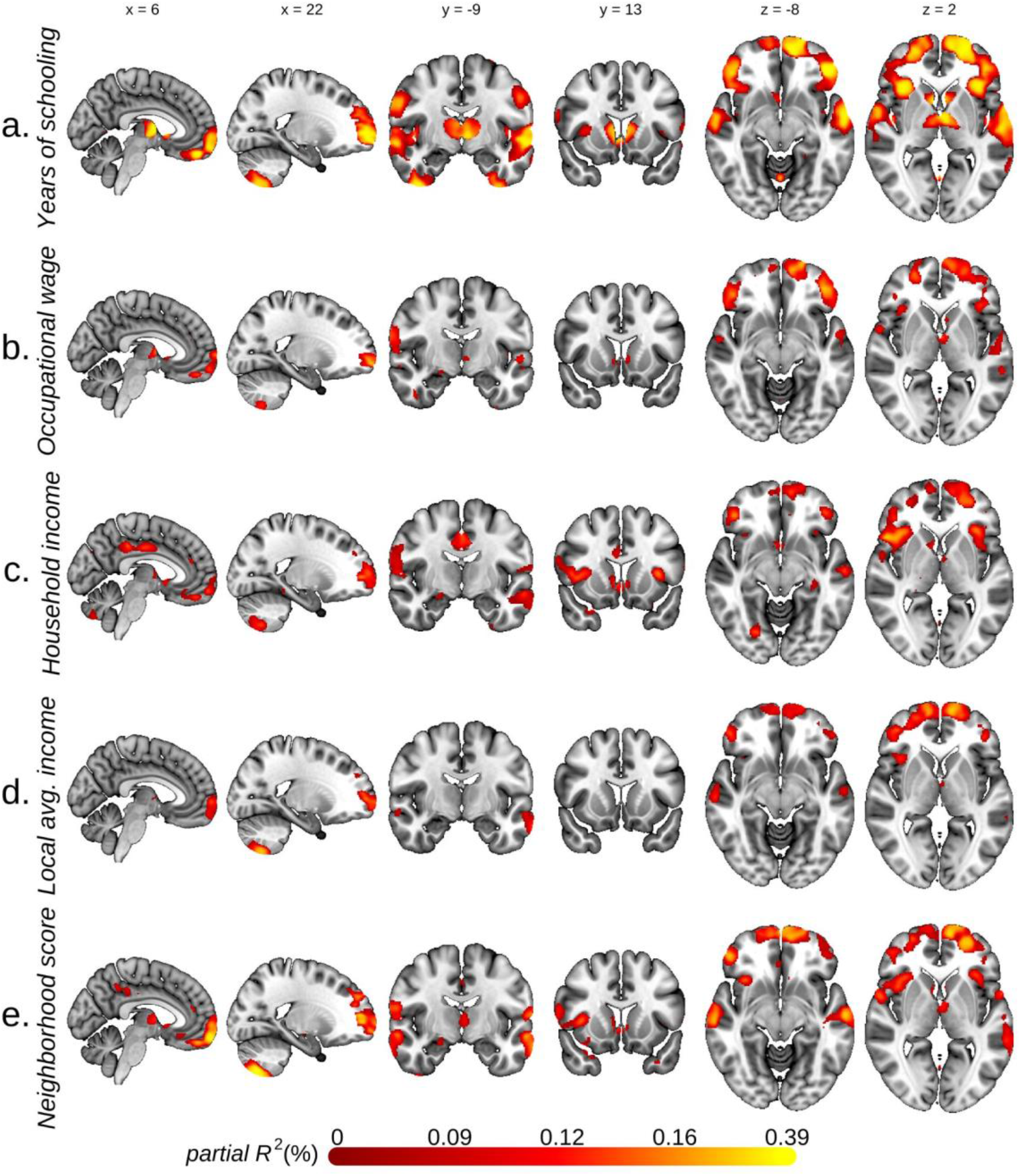
Voxel-based morphometry of grey matter volume and various socioeconomic measures. Univariate voxel-based morphometry results, with grey matter volume as the dependent variable and each of the five socioeconomic measures as the explanatory variable. Each figure plots the association sizes measured in partial *R^2^* for only the voxels significant at FWE rate of 5%. MNI coordinates are indicated. *N* = 30,954 (a), 27,307 (b), 29,839 (c), 29,006 (d), 28,405 (e)

**Fig. S6.**
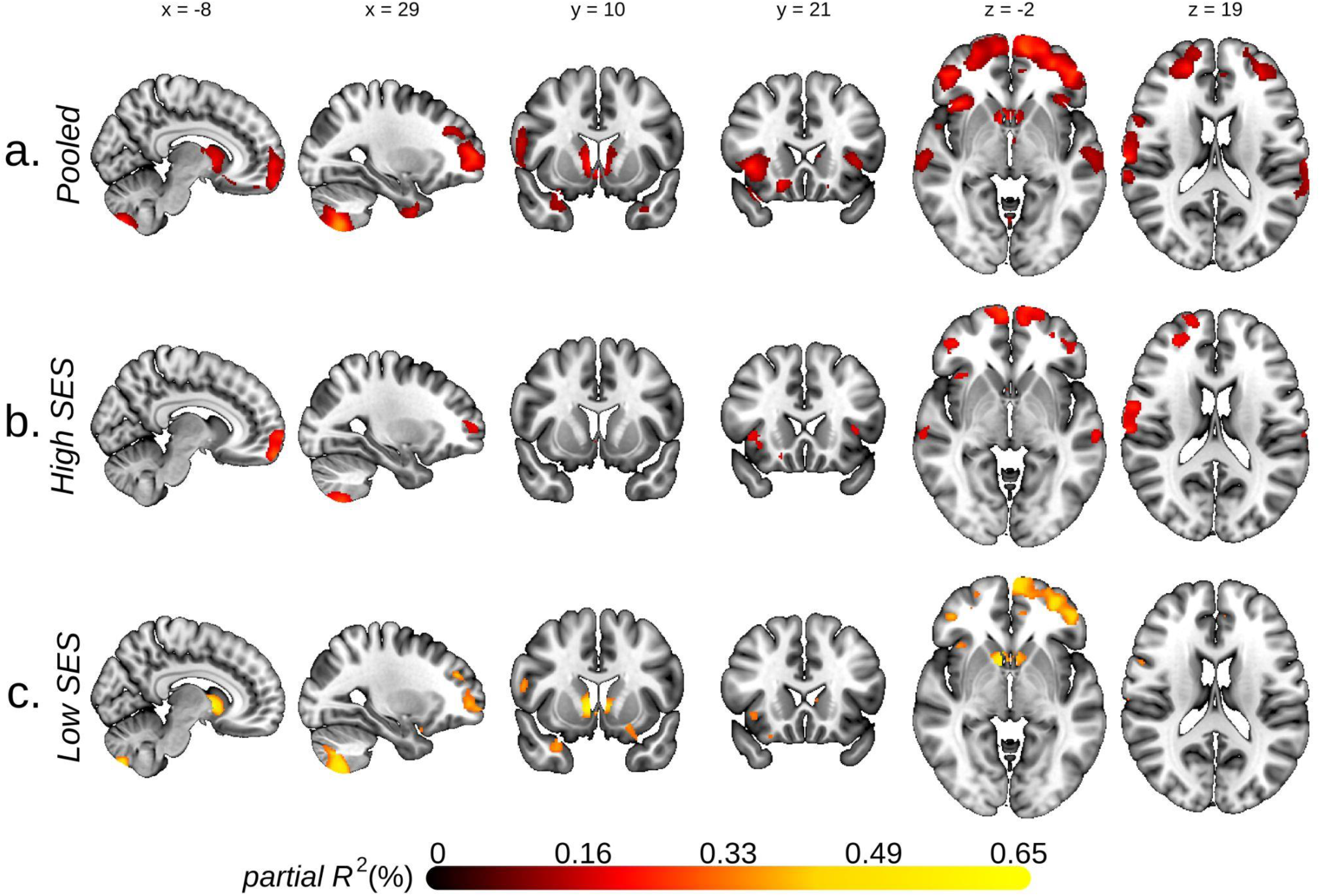
Stratified analyses on low and high socioeconomic status groups. Results from baseline voxel-based morphometry analysis conducted separately on low and high socioeconomic status (SES) groups as well as the pooled sample. Each figure plots the association sizes measured in partial *R^2^* for only the voxels significant at FWE rate of 5%. High and low SES groups were defined by National Statistics Socio-economic Classification (high if holding a managerial, administrative, or professional occupation). MNI coordinates are indicated.

**Fig. S7.**
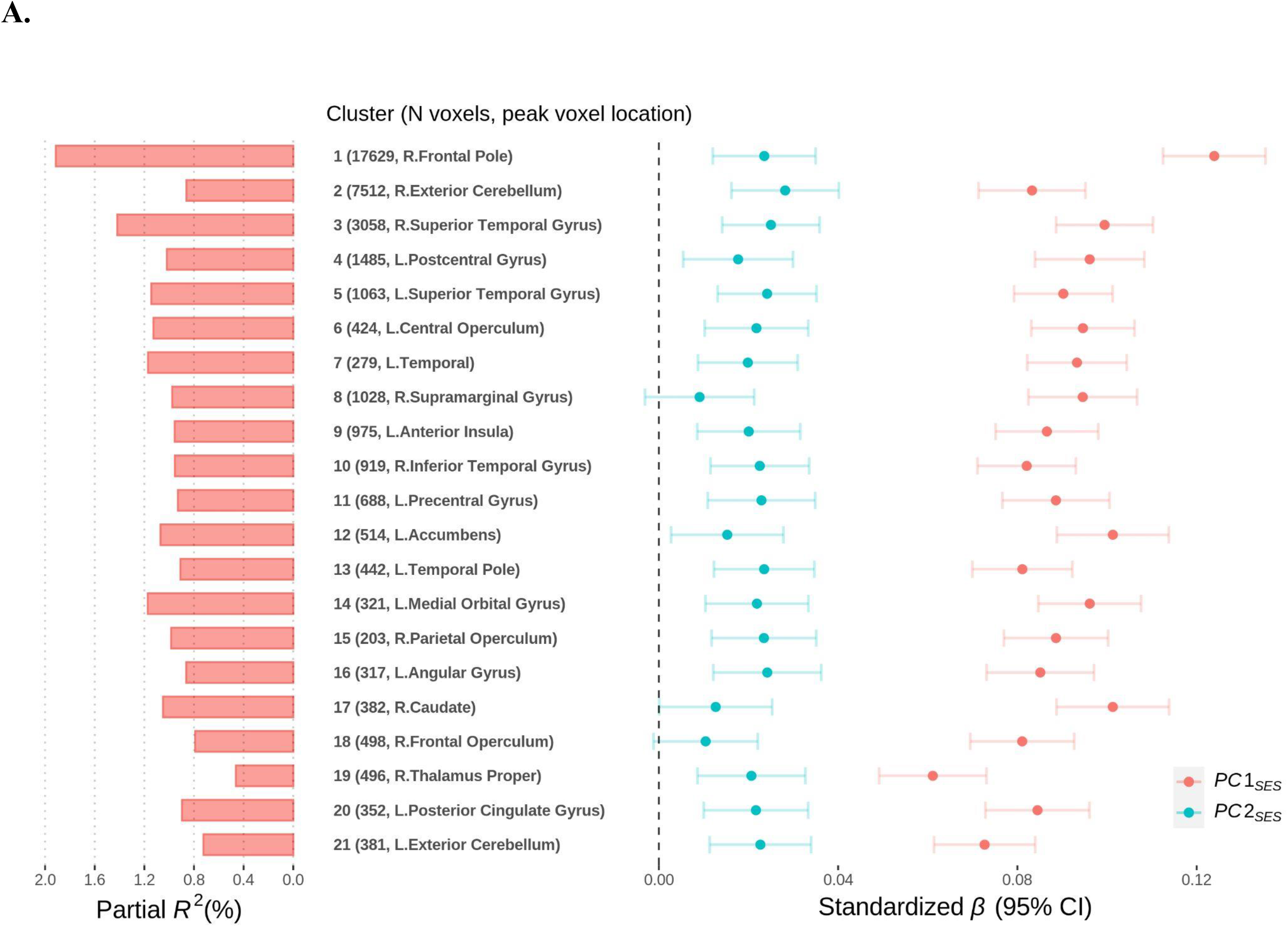
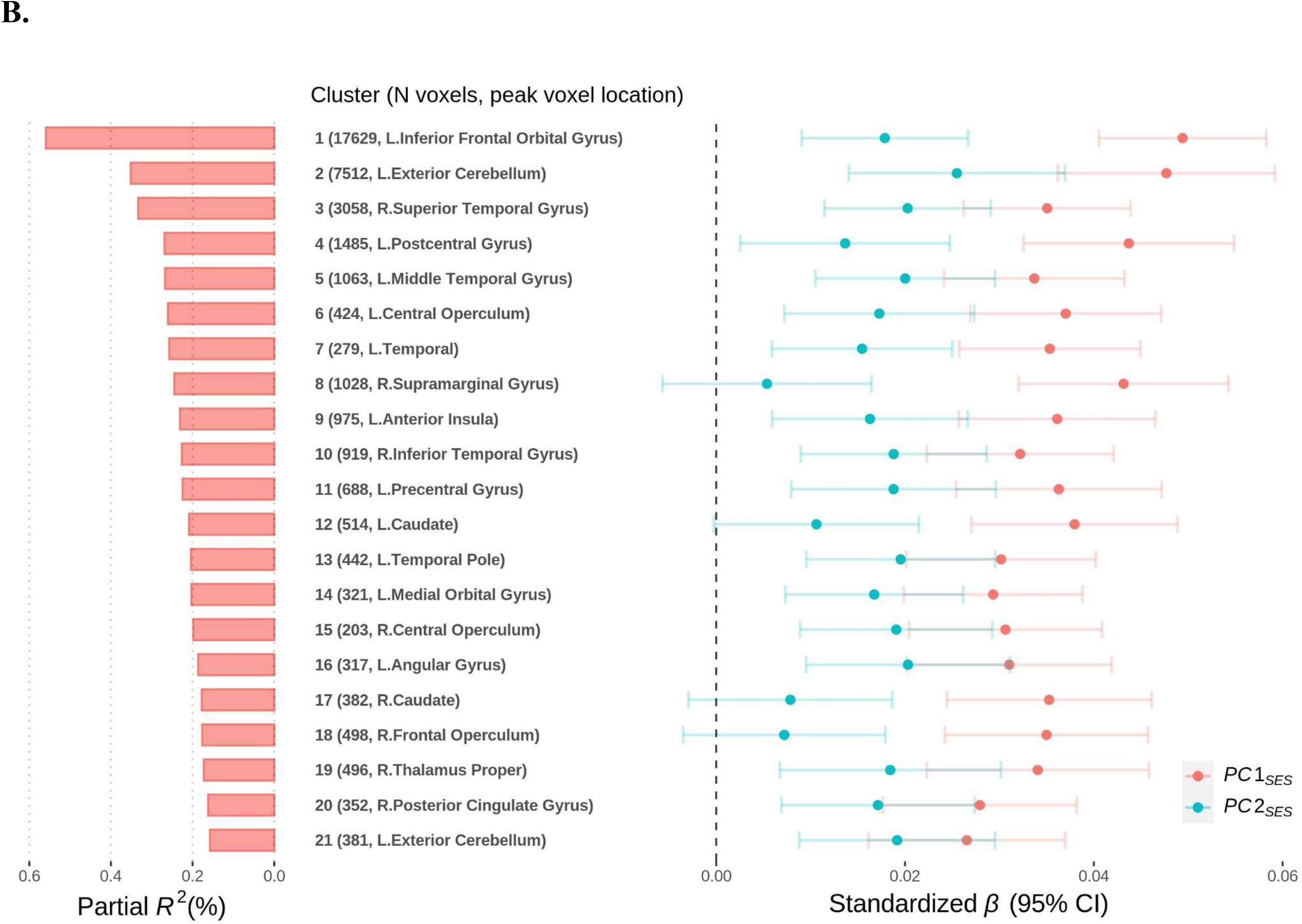
Standardized effect sizes of associations between socioeconomic status (SES) and grey matter volume (GMV) in voxel clusters. Results from regressing GMV in each cluster on *PC1*_SES_ and *PC2*_SES_. Total intracranial volume was not controlled for in **A.** but in **B.** On the left, partial *R^2^* from both *PC1*_SES_ and *PC2*_SES_ are reported and, on the right, the standardized coefficient estimates are plotted with their uncorrected 95% confidence intervals. The clusters were formed with at least 200 voxels showing significant associations at FWE rate of 5% level in the baseline TIV-adjustd voxel-based morphometry (VBM) results on *PC1*_SES_ and *PC2*_SES_. For each cluster, the anatomical location of the peak voxel from the VBM results is indicated. See Table S8 for more information about the clusters.

**Fig. S8.**
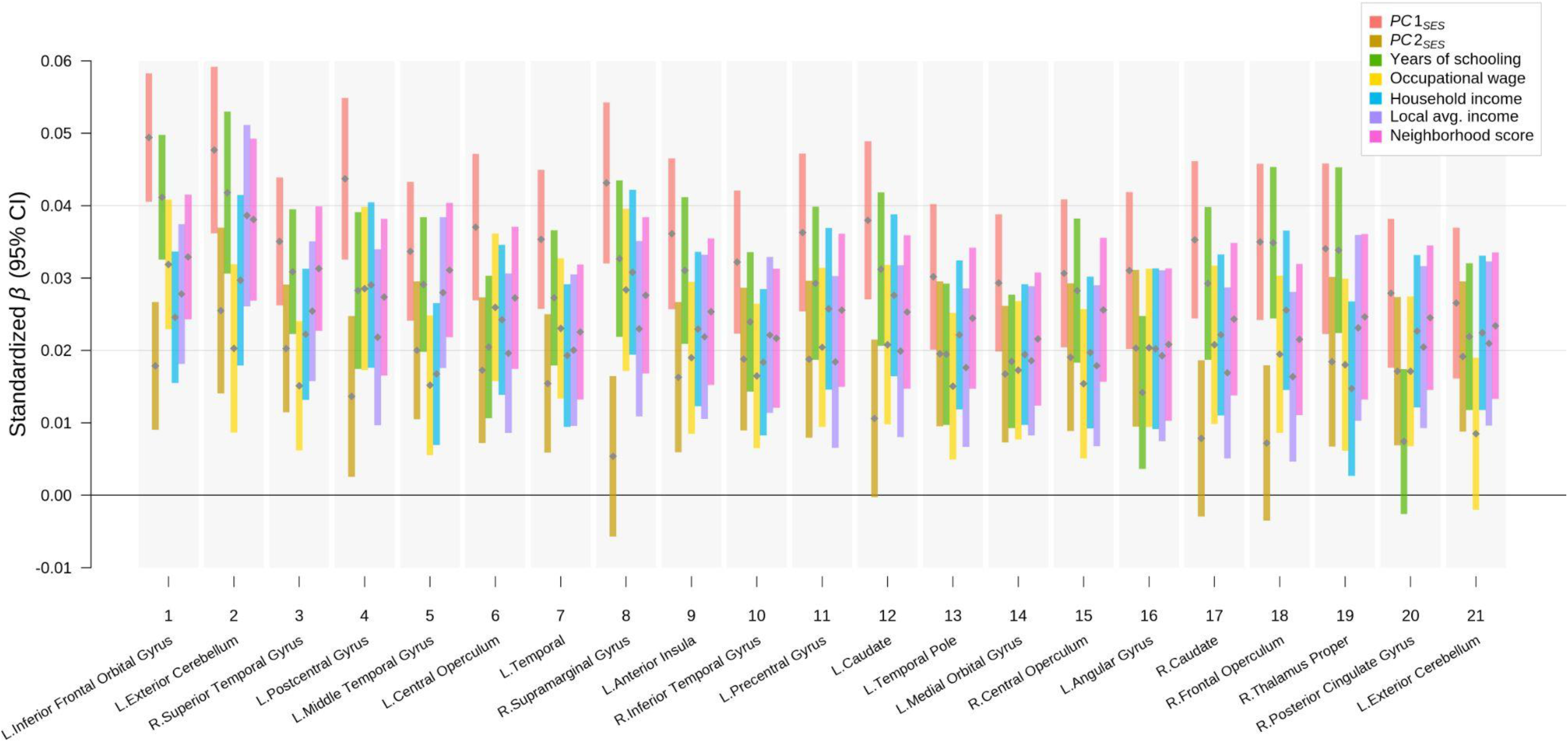
Standardized effect sizes of associations between various socioeconomic measures and grey matter volume (GMV) in voxel clusters. Results from regressing GMV in each cluster on each of the socioeconomic measures separately. The standardized coefficient estimates (grey points) are plotted with their uncorrected 95% confidence intervals (color bars). The clusters were formed with at least 200 voxels showing significant associations at FWE rate of 5% level in the baseline voxel-based morphometry (VBM) results on *PC1*_SES_ and *PC2*_SES_. The clusters are ordered by the strength of joint associations with *PC1*_SES_ and *PC2*_SES_. For each cluster, the anatomical location of the peak voxel from the VBM results is indicated at the bottom. See Table S8 for more information about the clusters.

**Fig. S9.**
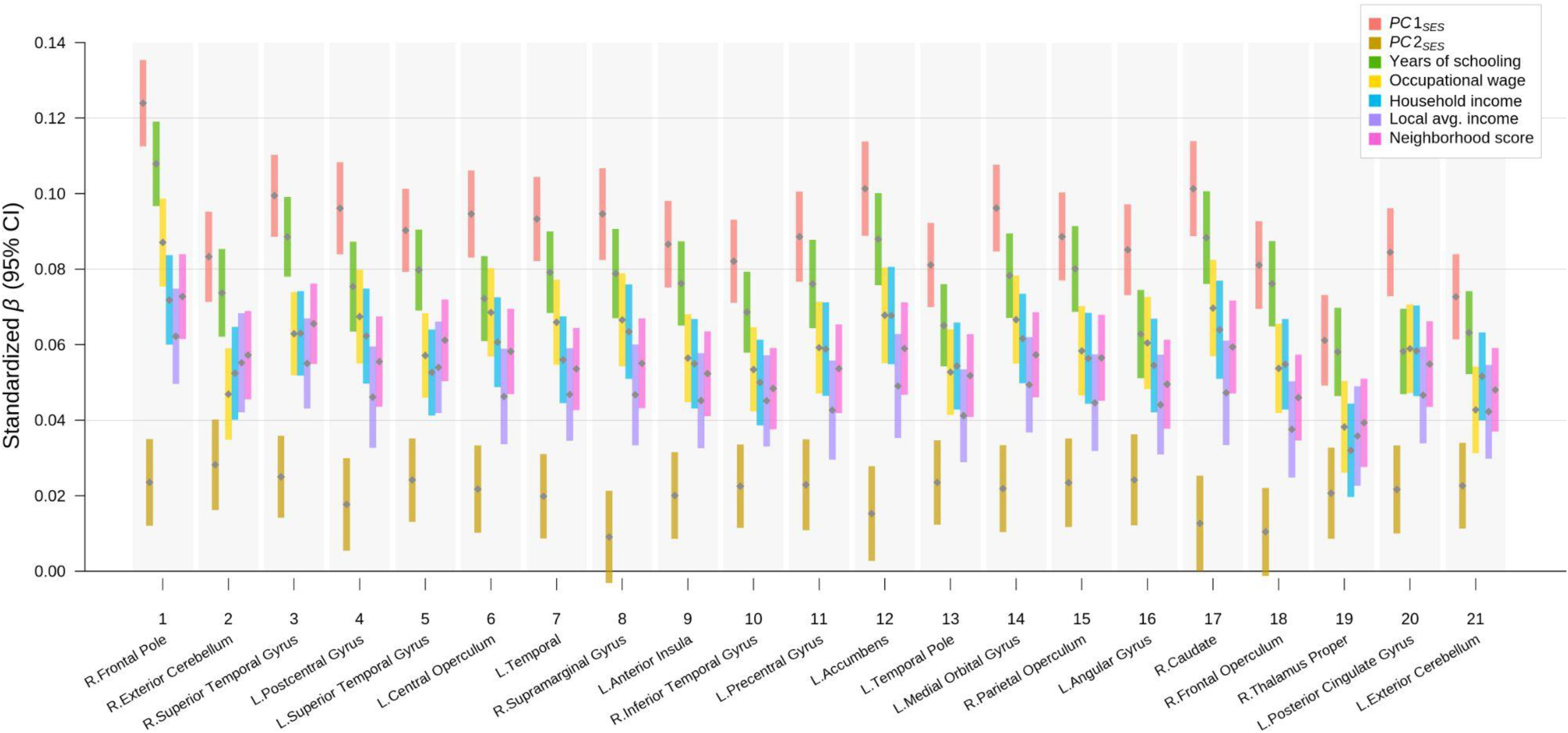
Standardized effect sizes of TIV-unadjusted associations between various socioeconomic measures and grey matter volume (GMV) in voxel clusters. Results from regressing GMV in each cluster on each of the socioeconomic measures separately while not controlling for total intracranial volume (TIV). The standardized coefficient estimates (grey points) are plotted with their uncorrected 95% confidence intervals (color bars). The clusters were formed with at least 200 voxels showing significant associations at FWE rate of 5% level in the baseline voxel-based morphometry (VBM) results on *PC1*_SES_ and *PC2*_SES_. The clusters are ordered by the strength of joint associations with *PC1*_SES_ and *PC2*_SES_. For each cluster, the anatomical location of the peak voxel from the VBM results is indicated at the bottom. See Table S8 for more information about the clusters.

**Fig. S10.**
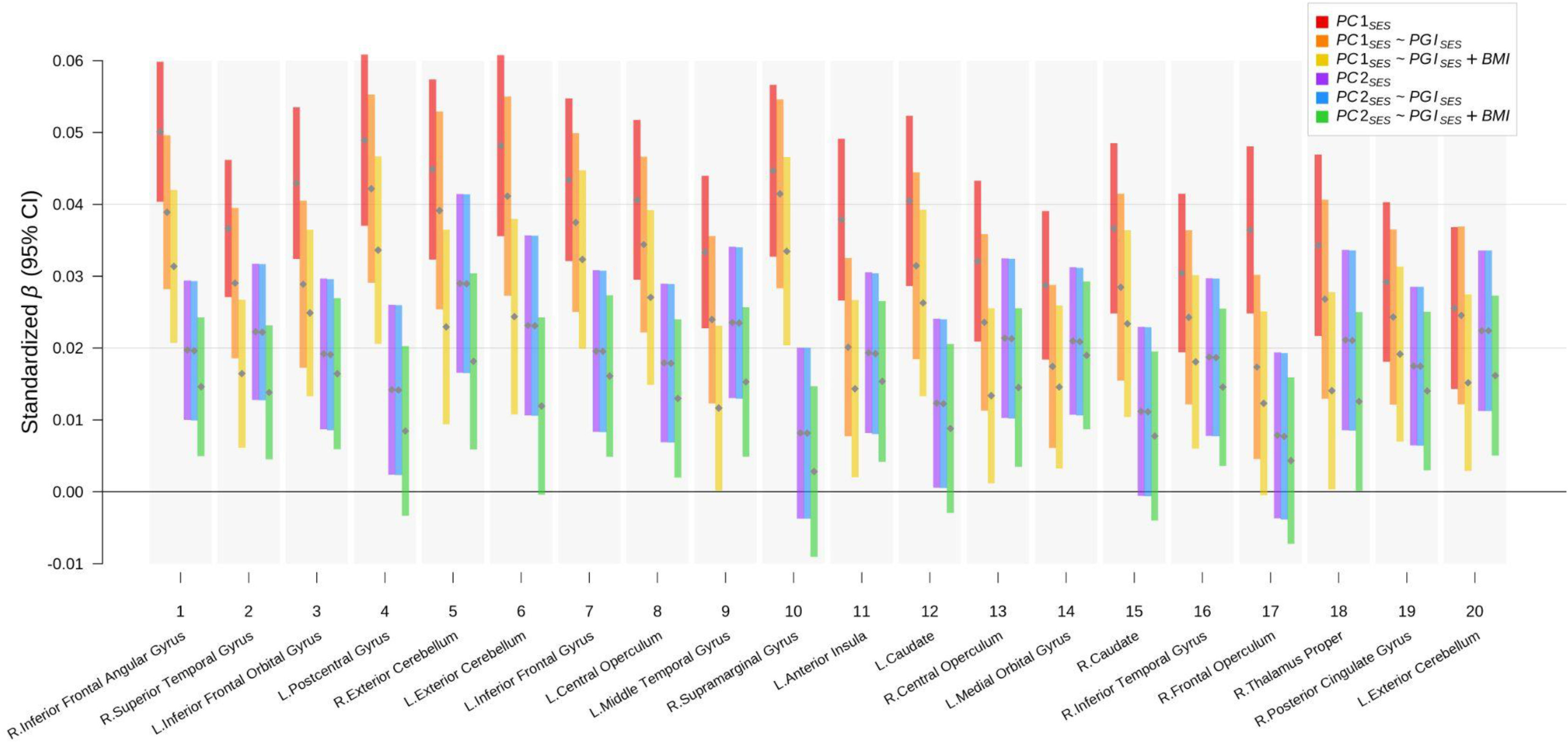
Standardized effect sizes of associations between socioeconomic status (SES) and grey matter volume (GMV) in voxel clusters with additional controls. Results from regressing GMV in each cluster on *PC1*_SES_ and *PC2*_SES_ while additionally controlling for *PGI*_SES_ and BMI. The standardized coefficient estimates (grey points) are plotted with their uncorrected 95% confidence intervals (color bars). The sample was restricted to individuals of European ancestry. Measurement error in *PGI_SES_* is adjusted for with genetic instrument variable (GIV) regression. The clusters were formed with at least 200 voxels showing significant associations at FWE rate of 5% level in the baseline voxel-based morphometry (VBM) results on *PC1*_SES_ and *PC2*_SES_. The clusters are ordered by the strength of joint associations with *PC1*_SES_ and *PC2*_SES_. For each cluster, the anatomical location of the peak voxel from the VBM results is indicated at the bottom. See Table S9 for more information about the clusters.

**Fig. S11.**
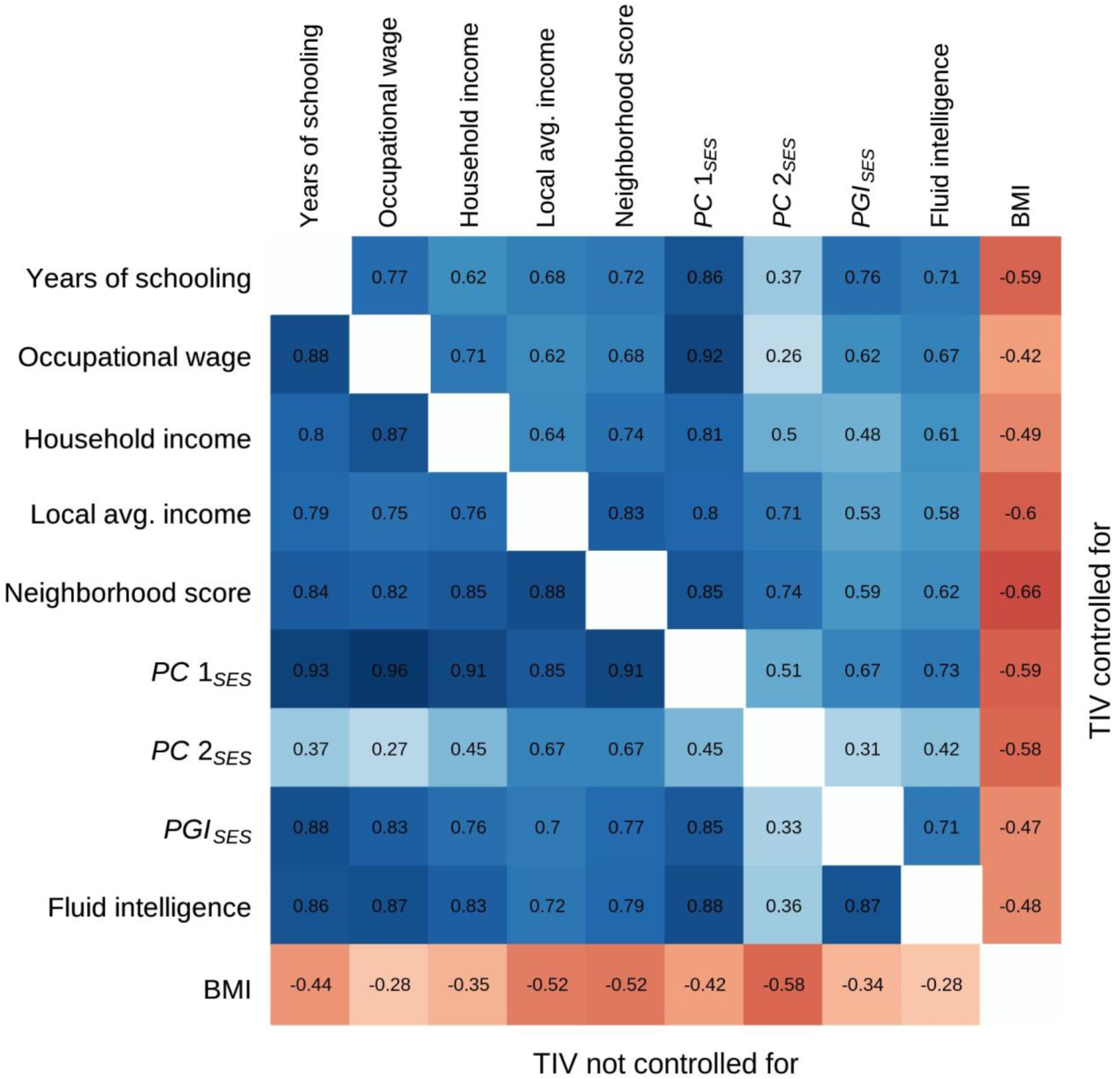
Grey-matter neuroanatomical correlations of various measures. The figure plots pairwise Pearson correlations computed from *T*-statistics of univariate grey-matter VBM results for each measure. The upper triangle reports the correlations of the *T*-statistics from the VBM analyses that adjusted for total intracranial volume (TIV), while the lower triangle from those that did not adjust for TIV.

**Fig. S12.**
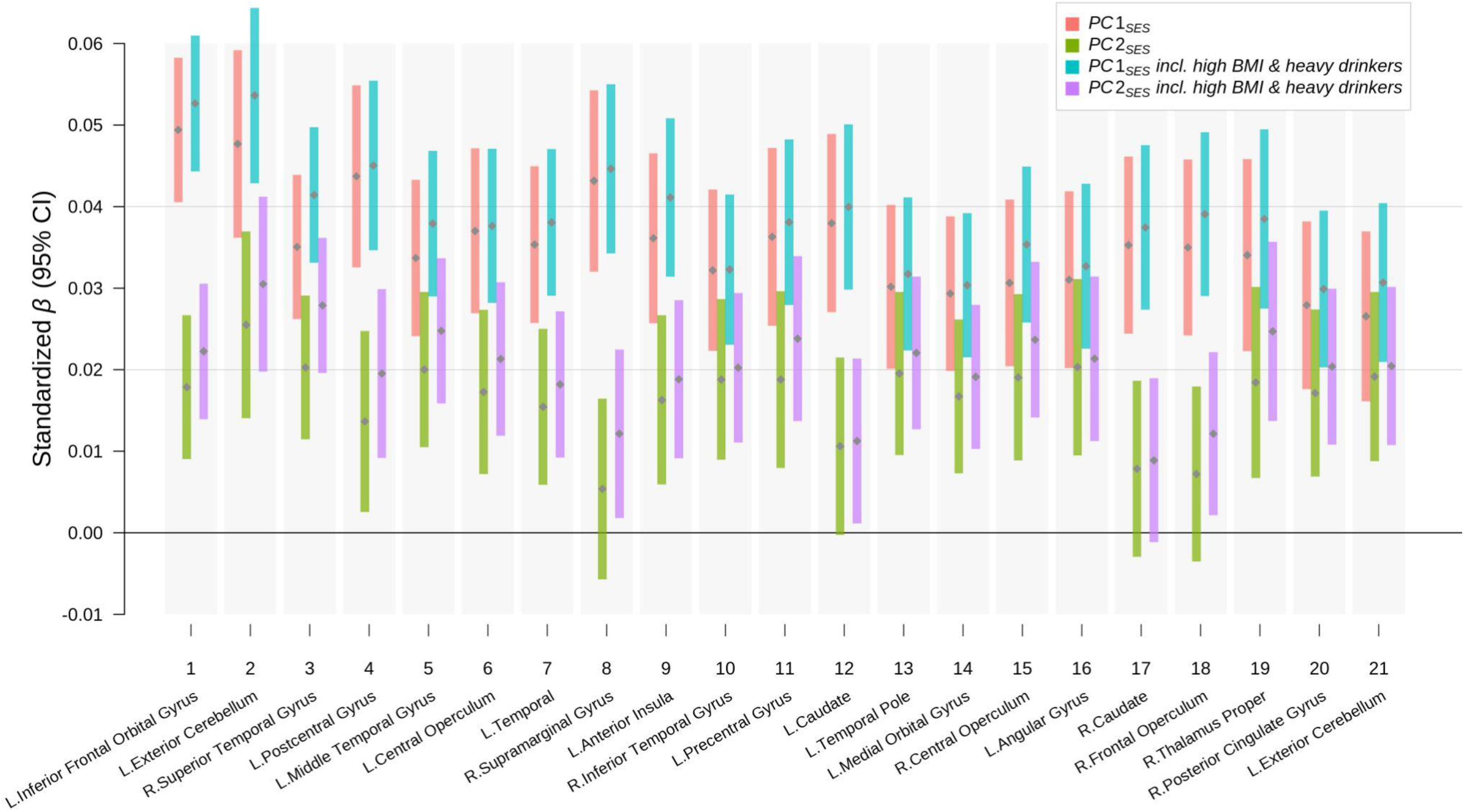
Standardized effect sizes of associations between socioeconomic status and grey matter volume (GMV) in voxel clusters with and without morbidly obese and heavy drinking individuals. Results from regressing GMV in each cluster on *PC1*_SES_ and *PC2*_SES_ with and without morbidly obese and heavy drinking individuals. The standardized coefficient estimates (grey points) are plotted with their uncorrected 95% confidence intervals (color bars). The clusters were formed with at least 200 voxels showing significant associations at FWE rate of 5% level in the baseline voxel-based morphometry (VBM) results on *PC1*_SES_ and *PC2*_SES_. The clusters are ordered by the strength of joint associations with *PC1*_SES_ and *PC2*_SES_. For each cluster, the anatomical location of the peak voxel from the VBM results is indicated at the bottom. See Table S8 for more information about the clusters.

**Fig. S13.**
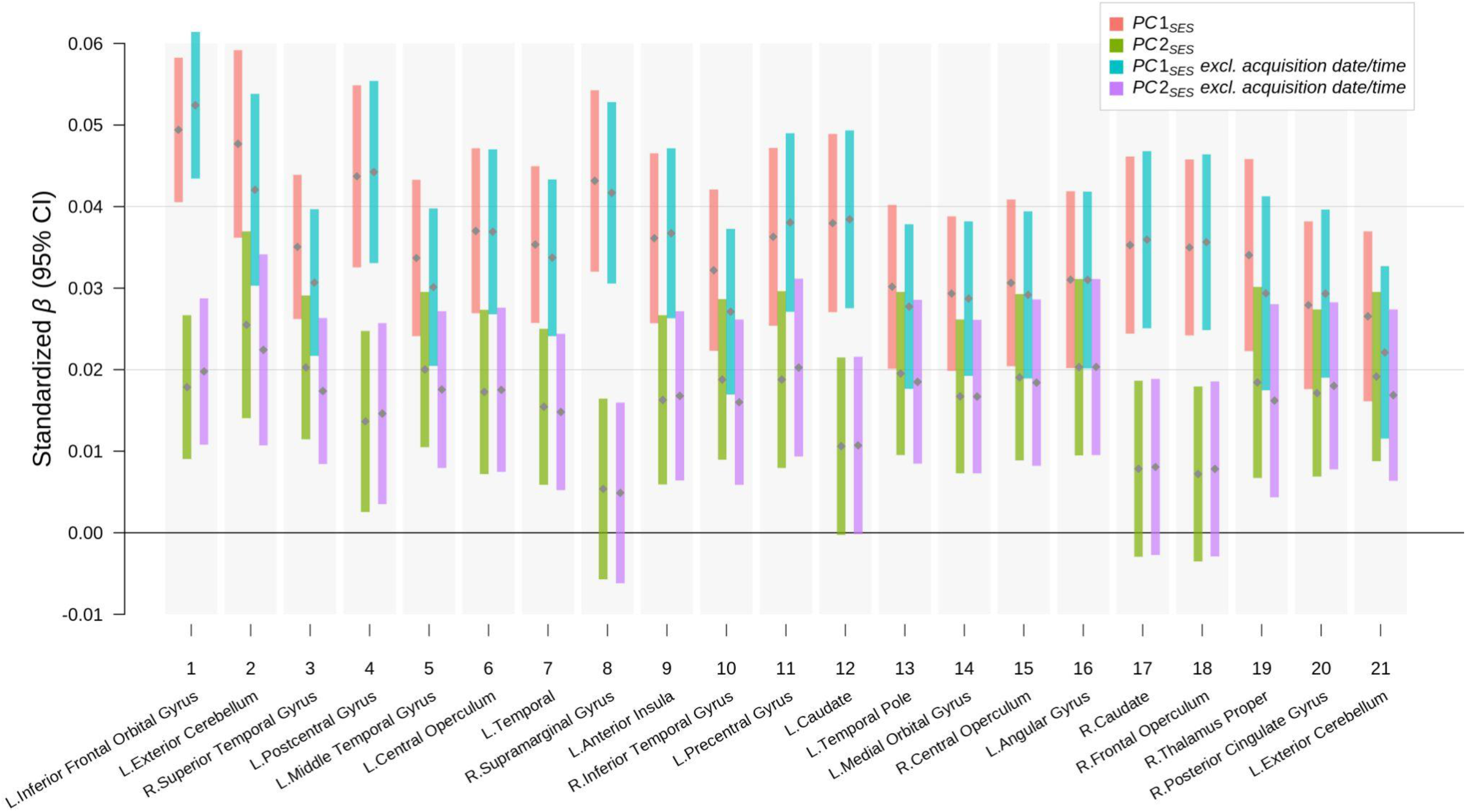
Standardized effect sizes of associations between socioeconomic status and grey matter volume (GMV) in voxel clusters with and without acquisition date and time as control variables. Results from regressing GMV in each cluster on *PC1*_SES_ and *PC2*_SES_ with and without the acquisition date and time as control variables. The standardized coefficient estimates (grey points) are plotted with their uncorrected 95% confidence intervals (color bars). The clusters were formed with at least 200 voxels showing significant associations at FWE rate of 5% level in the baseline voxel-based morphometry (VBM) results on *PC1*_SES_ and *PC2*_SES_. The clusters are ordered by the strength of joint associations with *PC1*_SES_ and *PC2*_SES_. For each cluster, the anatomical location of the peak voxel from the VBM results is indicated at the bottom. See Table S8 for more information about the clusters.

**Fig. S14.**
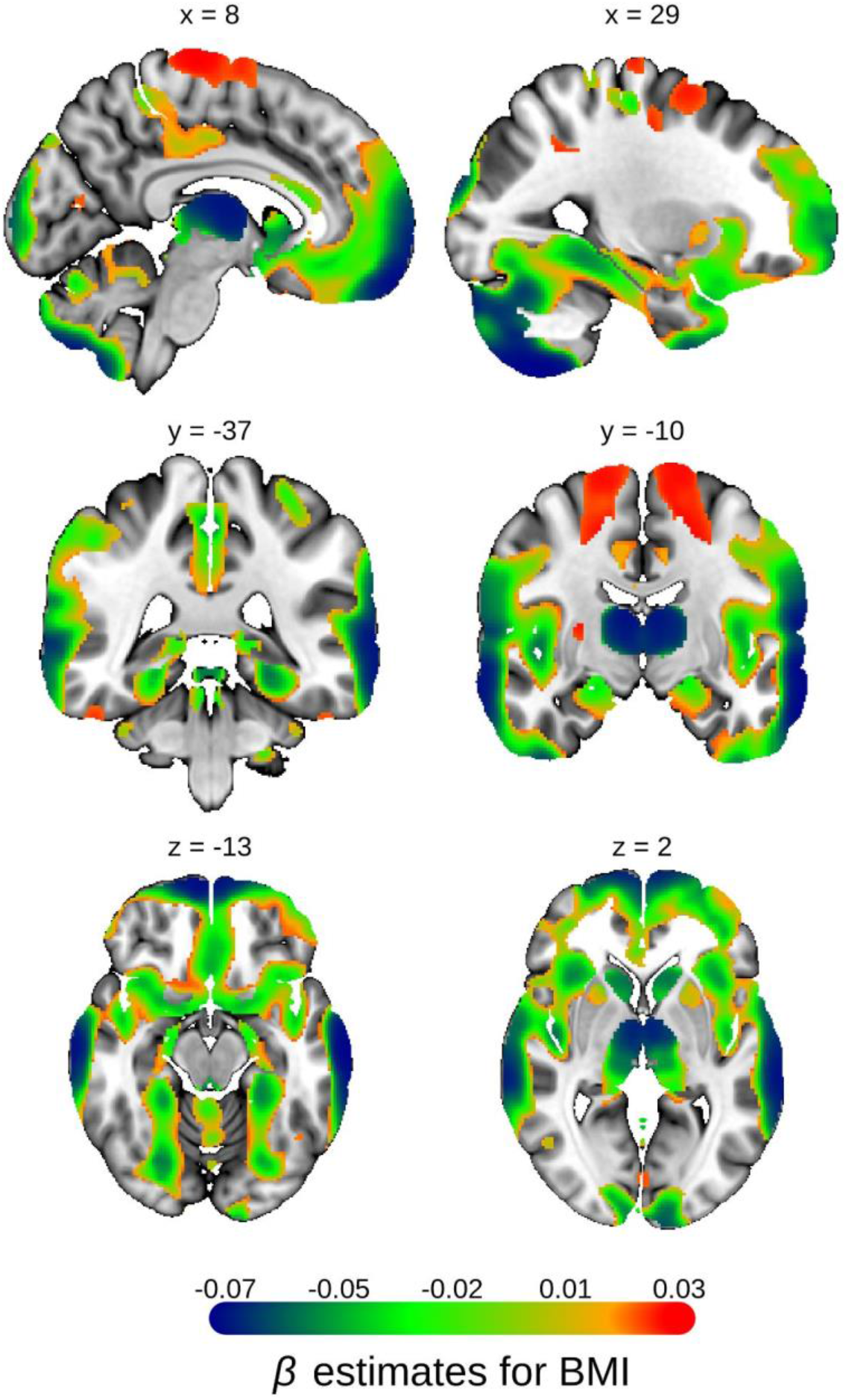
Voxel-based morphometry (VBM) of body mass index (BMI) Univariate VBM results on BMI, with grey matter volume (GMV) as the dependent variable. The Beta estimates are plotted for voxels significant at FWE rate of 1% level with partial *R^2^*>0.02%. The Beta estimates are reported in the standard deviation unit of GMV. MNI coordinates are indicated.

**Table S1.**
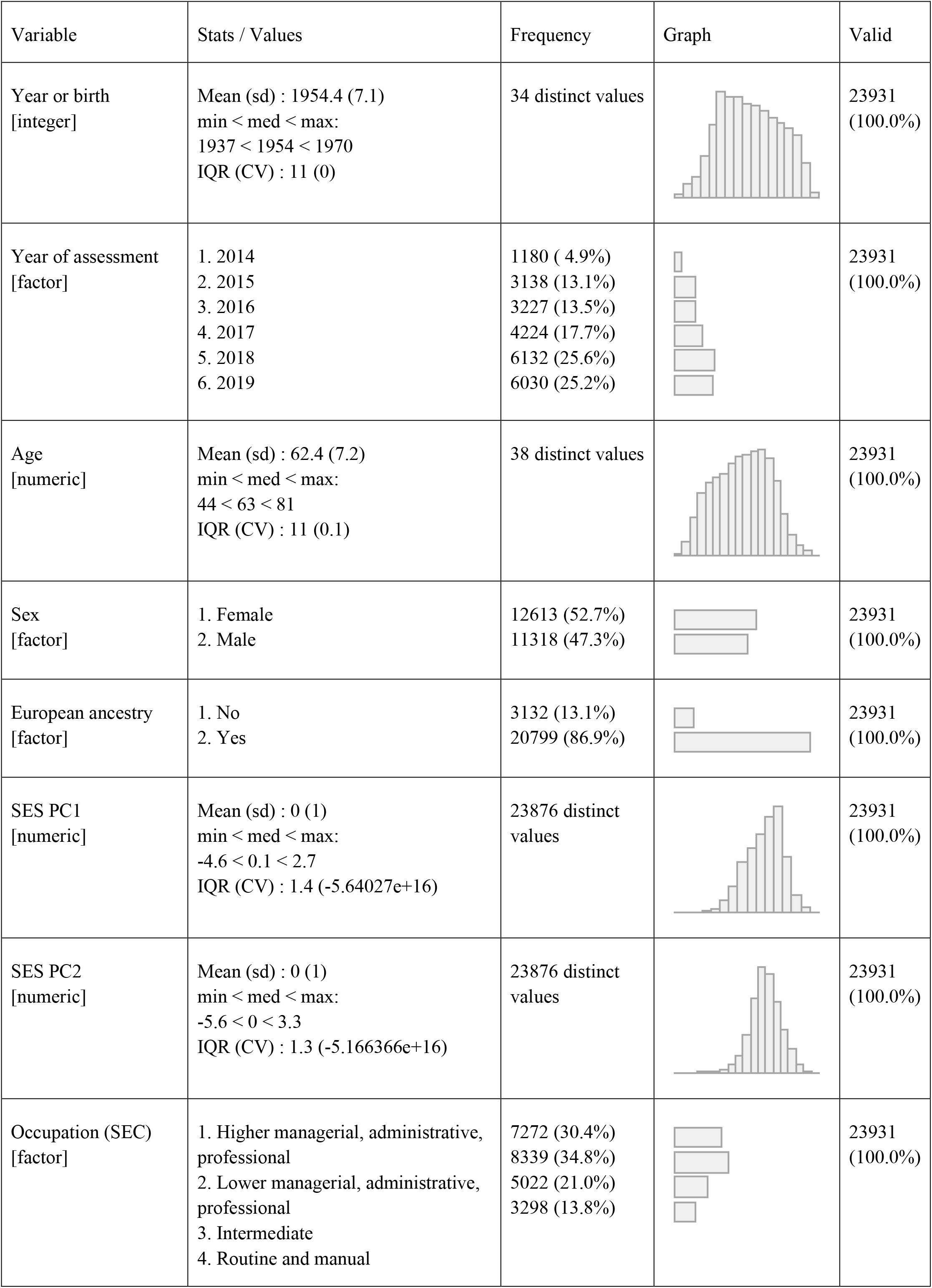
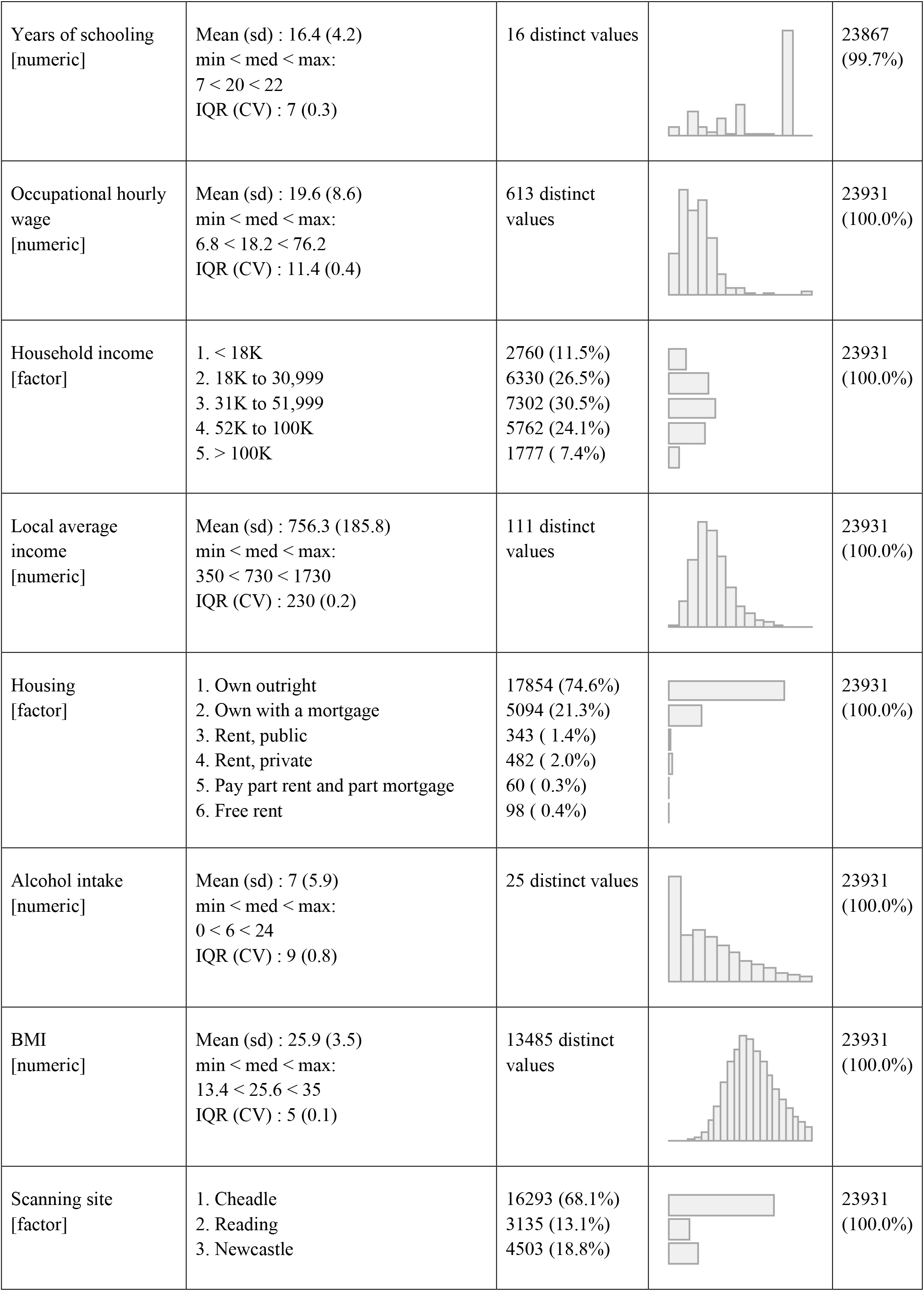
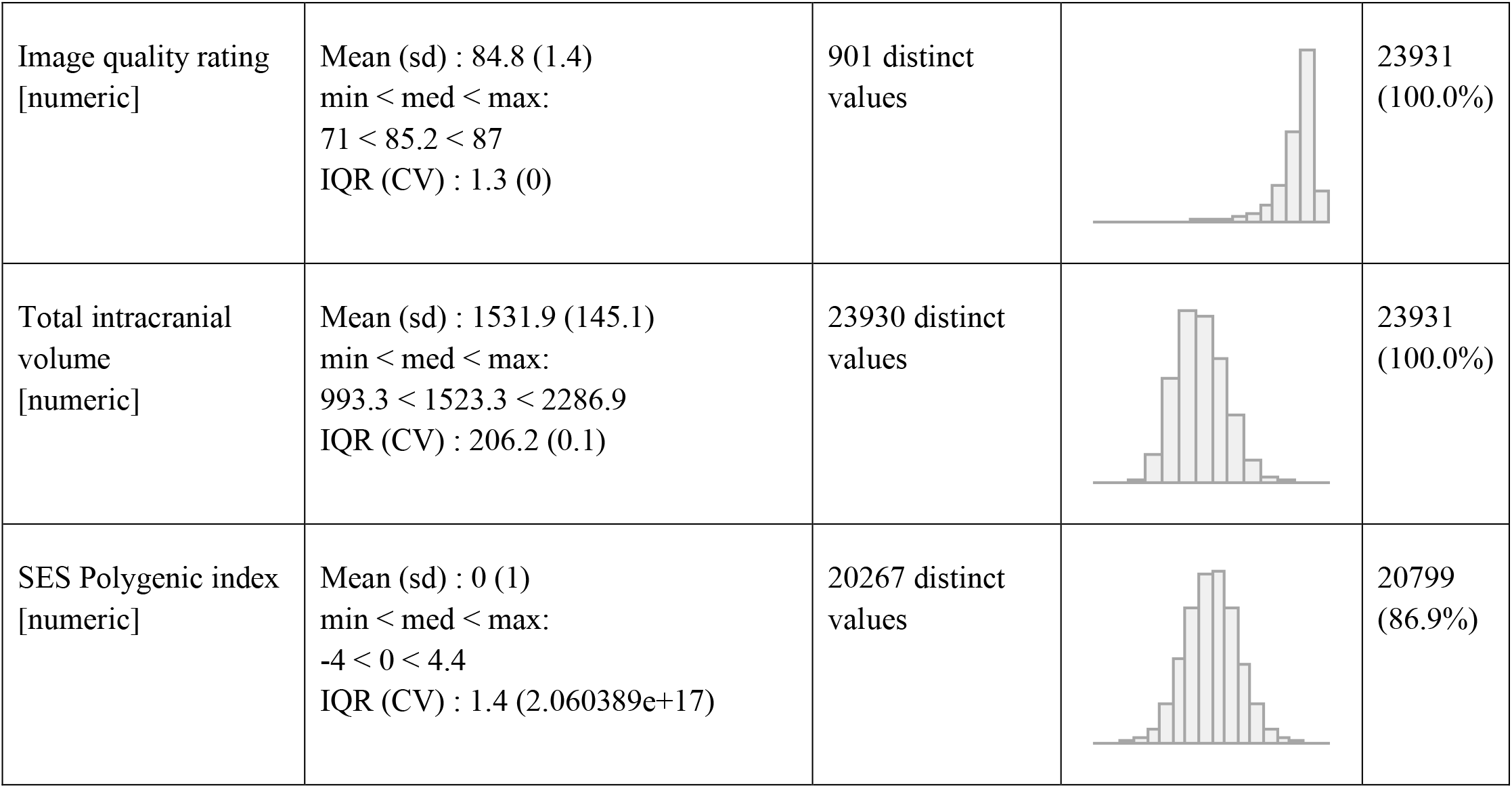
Descriptive statistics

## Footnotes

1 The sum of data fields 1568,1578,1588,1598, and 1608.

2 We first estimated a correlation matrix with elements being correlation estimates between individuals computed from voxel-level GMV. The mean voxel correlation corresponds to column-wise (or equivalently row-wise) averages of this matrix.

3 One example is using the UK’s National Statistics Socio-economic Classification, which reduces the occupation data to 3 or 8 classes.

4 While the UKB offers the occupation codes in 4-digits, we used 3-digit information to “blur” the data on purpose in order to address possible classification errors for such finely-defined occupation groups. Also, there were some 4-digit level occupation groups where too few individuals were observed, which can add more noise than signal. On the contrary, for occupational wages, we used 4-digit SOC codes because the wage data from the national statistics was used and classification errors should have less impact.

5 This method takes a weighted average of multiple outcomes such that outcomes highly correlated with each other are assigned less weight, while outcomes receive more weight if they are less correlated and therefore represent new information. See (*44*) for details.

6 The analysis plan specified 5 degrees of freedom for this, but we used 3 degrees of freedom due to rank deficiency.

7 This corresponds to everyone in the relatedness table reported by the UKB (minimum kinship coefficient = 0.04). See (*24*) for more details.

8 For the lowest and highest brackets, which are open-ended, 3/4 times the upper bound and 4/3 times the lower bound were used as the midpoint, respectively.

9 MTAG was used especially because it is robust to the relatedness between the samples. MTAG can be viewed as a generalization of the conventional inverse-variance-weighted meta-analysis.

10 Limbic, cerebellum, insular, frontal, parietal, occipital, and temporal

11 Here, we did not interact TIV with the site of acquisition for simplicity when obtaining these estimates. There was no much difference in TIV due to images taken in different acquisition sites.

12 These concepts were taken from https://www.cognitiveatlas.org/concepts/categories/all on 2 July 2021.

