## Supplementary notes and figures for "Human brain anatomy reflects separable genetic and environmental components of socioeconomic status"

### Supplementary Note

1. Study overview

2. Sample description

3. Measures
   - 3.1. Imaging-derived phenotypes (IDPs)
   - 3.2. SES measures
     - 3.2.1. Available measures of SES in the UK Biobank
       - 3.2.1.1. Highest qualification
     - 3.2.2. Data reduction by principal component analysis
   - 3.3. Control variables
   - 3.4. Genome-wide association studies and construction of the polygenic index for SES

4. Statistical analyses
   - 4.1. Voxel-based Morphometry (VBM) analysis
     - 4.1.1. Baseline analysis
     - 4.1.2. Controlling for total intracranial volume (TIV)
     - 4.1.3. Multiple testing correction
     - 4.1.4. Stratified analysis of high and low SES groups
     - 4.1.5. VBM of Individual SES measures
   - 4.2. Estimating the overall association between SES and GMV structure
   - 4.3. Incorporating genetics
     - 4.3.1. VBM with PGI
     - 4.3.2. Testing differences in SES-GMV associations with and without PGI as a control variable
     - 4.3.3. Measuring differences in SES-GMV associations with and without PGI as a control variable
     - 4.3.4. Measurement error correction for PGI
   - 4.4. Functional annotations

5. Interpretation
   - 5.1. Brain, SES, and genetics
   - 5.2. Interpretation of the polygenic index for SES

6. Supplementary analyses
   - 6.1. Heritability and genetic correlation
   - 6.2. Testing differences in residual SES-GMV associations due to BMI
   - 6.3. Heterogeneity
   - 6.4. Controlling for alcohol consumption

| Measure | Data fields | Variable | Description |
|---|---|---|---|

[3] One example is using the UK's National Statistics Socio-economic Classification, which reduces the occupation data to 3 or 8 classes.

| | | | |
|---|---|---|---|
| Occupation | 132, 20024, 22617 | 81 categories | Job codes for the latest job held before the age of 65, coded in 3-digit UK standard occupational classification (SOC) 2000.[4] |
| Occupational wages (log)* | 132, 20024, 22617 | Continuous | Sex-specific average occupational wage matched to 4-digit SOC codes. The wage data are obtained from the UK's Office for National Statistics, averaged over 2002-2010. |
| Household income | 738 | 5 categories | Average total household income before tax |
| Housing type | 680 | 6 categories | e.g., own outright, own with mortgage, rent |
| Local average household income (log)* | 20074, 20075 | Continuous | Derived by matching home locations to Middle-layer Super Output Areas. The income data are obtained from the UK's Office for National Statistics (England and Wales only) |
| Neighborhood SES score* | 26411, 26412, 26414 | Continuous | Weighted average of three composite deprivation indices for income, employment, and education, constructed at the level of Lower-layer Super Output Areas (England only). The weights were derived by using the inverse of the correlation matrix.[5] |
| Highest qualification* | 6138 | 7 categories | See the following description in section 3.2.1.1. |

*Note*: The asterisks (*) indicate that the variable is derived.

- *3.2.1.1. Highest qualification*

This subsection describes how we derived the highest qualification. During the assessment, participants were asked to choose qualifications that they have from the below options:

$$s_i = \sum_{j=1}^{M} \hat{\beta}_j x_{ij}$$

where $x_{ij}$ represents the genotype of individual *i* for SNP *j* coded as the count of the reference allele. We estimated the weights $\hat{\beta}_j$ from genome-wide association studies (GWAS), which conduct univariate regressions of an outcome on each SNP across the genome. The resulting estimates were then adjusted for the correlation between the SNPs to obtain the weights $\hat{\beta}_j$.

$$(1)\quad GMV_i^j = \beta_1^j PC1_{SES,i} + \beta_2^j PC2_{SES,i} + Z_i^T \gamma^j + \varepsilon_i^j$$

where the GMV of voxel *j* is regressed on the two SES PCs. The vector $Z_i$ include the control variables listed in Section 3.3. $\varepsilon_i^j$ is the error term. The GMV and the SES PCs were standardized

[9] MTAG was used especially because it is robust to the relatedness between the samples. MTAG can be viewed as a generalization of the conventional inverse-variance-weighted meta-analysis.

to have zero mean and unit variance. An *F*-test was used for each voxel to test whether there is significant association between voxel *j*'s GMV and the SES PCs jointly with the the null hypothesis $\beta_1^j = \beta_2^j = 0$. We measured the association size by the variance of interest in GMV explained by the SES PCs beyond the covariates of no interest, *i.e.*, partial $R^2 := (R^2_{PC+Z} - R^2_Z) / (1 - R^2_Z)$. $R^2_{PC+Z}$ is the $R^2$ from the unrestricted model, which includes the two SES PCs and the covariates of no interest, and $R^2_Z$ is the $R^2$ from the restricted model, which only includes the covariates of no interest. We also quantified the relative contribution of $PC1_{SES}$ in the overall association size by $(R^2_{PC1+Z} - R^2_Z) / (R^2_{PC+Z} - R^2_Z)$. We used permutation testing to correct for multiple hypothesis testing across voxels (see Section 4.1.3 for details).

After the estimation, we anatomically labeled the voxels using the Neuromorphometrics atlas provided in CAT12. For a summary purpose, we also generated cluster-based estimates. Each cluster consists of at least 200 neighboring voxels within the lobe[10] which are significant at the familywise error rate of 5% in the baseline model. We then repeated the same analysis with mean GMV of these clusters.

###### *4.1.2. Controlling for total intracranial volume (TIV)*

Analyses that aim to identify associations between localized GMV and outcomes typically control for TIV, since volumetric brian measures scale with the head size. However, controlling for TIV as a linear covariate has important statistical implications for identifying localized GMV patterns linked to SES, because TIV is positively correlated with both SES and regional GMV. In Fig 2B, GMV of some voxels appear to have negative association with SES when the TIV is included as a control variable in the model. On the contrary, Fig 2A shows that almost all the voxel-level GMV are positively associated with SES when TIV is not controlled for. This result indicates that the absolute GMV-SES association is unlikely to be negative in any brain region.

To formally illustrate this point, consider a VBM model for SES with only the TIV as a covariate without loss of generality:

$$(2)\quad GMV_i = \beta_{\sim TIV} SES_i + \gamma TIV_i + \varepsilon_i$$

where $GMV_i$ is the GMV of some voxel and $\beta_{\sim TIV}$ denotes the association between the voxel's GMV and SES while TIV is accounted for. Each variable is standardized to have zero mean and unit variance without loss of generality. $\gamma$ corresponds to the association between the GMV and the TIV, conditional on SES. The linear dependence between the TIV and SES can be described

[10] Limbic, cerebellum, insular, frontal, parietal, occipital, and temporal

as: $E[TIV_i|SES_i] = \lambda SES_i$. If we denote β as the coefficient of SES from the regression of the GMV on SES without the TIV as a covariate, $\beta_{\sim TIV}$ can be written as:

(3) $\beta_{\sim TIV} = \beta - \lambda\gamma$

Therefore, if both λ and γ are positive and large, $\beta_{\sim TIV}$ can be negative even when β is positive. Our data suggests that this is indeed the case: With the baseline model[11], we estimated $\hat{\lambda} = 0.10$ for $PC1_{SES}$ and $\hat{\lambda} = 0.01$ for $PC2_{SES}$. $\hat{\gamma}$ was on average 0.46 with the minimum=0.11 (right exterior cerebellum) and the maximum=0.72 (left gyrus rectus). Since estimates of β are positive for the vast majority of the voxels, one cannot conclude that the absolute GMV-SES association is truly negative even when estimates of $\beta_{\sim TIV}$ are negative. Instead, such negative estimates are evidence that λγ is large relative to β and that the GMV-SES association is essentially very small or non-existent for these regions.

$$(4)\ \ GMV_i^j = \tilde{\beta}_1^j PC1_{SES,i} + \tilde{\beta}_2^j PC2_{SES,i} + \theta^j PGI_i + Z_i^T \tilde{\gamma}^j + \tilde{\varepsilon}_i^j$$

$$(5)\ \ PGI_i = \delta_1 PC1_{SES,i} + \delta_2 PC2_{SES,i} + Z_i^T \psi + u_i$$

Using vector notations: $\beta^j = [\beta_1^j\ \beta_2^j]^T$, $\tilde{\beta}^j = [\tilde{\beta}_1^j\ \tilde{\beta}_2^j]^T$, $\delta = [\delta_1\ \delta_2]^T$, which are all length-2 vectors, it can be shown:

$$(6)\ \ \beta^j - \tilde{\beta}^j = \theta^j \cdot \delta = \Delta^j$$

Therefore, the vector $\Delta^j$ represents the difference in the SES-GMV association for voxel *j* due to controlling for $PGI_{SES}$. $\Delta^j$ can be estimated as the product of estimates of $\hat{\theta}^j$ and $\hat{\delta}$ from the model (4) and (5), respectively. A Wald test was then used to test the null $\Delta^j = 0$ with the test statistic: $\hat{\Delta}^{jT}\ \hat{Var}(\hat{\Delta}^j)^{-1}\hat{\Delta}^j \sim \chi^2_2$, where $\hat{Var}(\hat{\Delta}^j)$ was approximated by the delta method: $\hat{Var}(\hat{\Delta}^j) \approx \hat{Var}(\hat{\theta}^j)\hat{\delta}^T\hat{\delta} + \hat{\theta}^{j2}\ \hat{Var}(\hat{\delta})$. Note that this analysis is statistically equivalent to a mediation analysis with $PGI_{SES}$ being a mediator (*60*). We conducted this test only for the voxels whose GMV was significantly associated with the PCs. Then, the multiple testing was corrected for using Bonferroni correction (the corrected 5% threshold = $1.46 \times 10^{-6}$ with 34,188 tests).

##### *4.3.3. Measuring differences in SES-GMV associations with and without PGI as a control variable*

To represent the relative size of $\Delta^j$ in relation to partial $R^2$, we used the relative change in the net variation explained by the SES PCs after adding $PGI_{SES}$ to the model with the covariates of no interest: $[(R^2_{PC+Z} - R^2_Z) - (R^2_{PC+PGI+Z} - R^2_{PGI+Z})]\ /\ (R^2_{PC+Z} - R^2_Z)$. This measure is bounded between

0 and 1 as long as the sign of the coefficients for $PC1_{\text{SES}}$ and $PC2_{\text{SES}}$ do not change after controlling for $PGI_{\text{SES}}$. This expression can be interpreted as the percent change in the SES-GMV associations due to controlling for $PGI_{\text{SES}}$ and essentially the part of the SES-GMV association that can be attributed to $PGI_{\text{SES}}$. Note that, because $PC2_{\text{SES}}$ is barely predicted by $PGI_{SES}$ and even barely heritable (Table S5), the percent change in SES-GMV association after controlling for $PGI_{\text{SES}}$ is essentially due to the change in $PC1_{\text{SES}}$-GMV association. We can therefore rewrite the earlier expression as:

$$(7)\quad [(R^2_{PC+Z} - R^2_Z) - (R^2_{PC+PGI+Z} - R^2_{PGI+Z})] \,/\, (R^2_{PC+Z} - R^2_Z)$$

$$\approx [(R^2_{PC1+Z} - R^2_Z) - (R^2_{PC1+PGI+Z} - R^2_{PGI+Z})] \,/\, (R^2_{PC+Z} - R^2_Z)$$

$$= \Delta_{PC1} \,/\, (R^2_{PC+Z} - R^2_Z)$$

$$= [\Delta_{PC1} \,/\, (R^2_{PC1+Z} - R^2_Z)] \times [(R^2_{PC1+Z} - R^2_Z) \,/\, (R^2_{PC+Z} - R^2_Z)]$$

where $\Delta_{PC1} = (R^2_{PC1+Z} - R^2_Z) - (R^2_{PC1+PGI+Z} - R^2_{PGI+Z})$, the change in the net variance explained by $PC1_{\text{SES}}$ after controlling for $PGI_{SES}$. Hence, the percent change in SES-GMV association is roughly the product of the percent change in $PC1_{\text{SES}}$-GMV association ($\Delta_{PC1} / (R^2_{PC1+Z} - R^2_Z)$) and the relative contribution of $PC1_{\text{SES}}$ in the overall SES-GMV association ($(R^2_{PC1+Z} - R^2_Z) / (R^2_{PC+Z} - R^2_Z)$). A larger share of SES-GMV association can be attributed to $PGI_{SES}$ if genetic factors linked to SES play a bigger role for $PC1_{\text{SES}}$-GMV association and/or if $PC2_{\text{SES}}$ contributes relatively less to the overall SES-GMV association.

##### *4.3.4. Measurement error correction for PGI*

$PGI_{\text{SES}}$ is a noisy proxy of true linear effects of common genetic variants that are linked to SES because GWAS estimates of individual SNP effects are obtained from finite sample sizes. The difference between the true PGI and the available PGI can be viewed as the classic measurement error, which leads to an attenuation bias in the coefficient estimate for the $PGI_{\text{SES}}$. Nonetheless, it is still possible to account for the linear effects of common genetic variants that the true $PGI_{\text{SES}}$ would capture under reasonable assumptions. We addressed this attenuation bias by using genetic instrumental variable (GIV) regression (*35*). The essential idea is that the true $PGI_{\text{SES}}$ can be recovered from a noisy $PGI_{\text{SES}}^{(1)}$ by using another $PGI_{\text{SES}}^{(2)}$ as an instrumental variable that was derived from a different GWAS sample. The crucial assumption here is that the noise in $PGI_{\text{SES}}^{(1)}$ and $PGI_{\text{SES}}^{(2)}$ is uncorrelated to each other. GIV regression can address the measurement error in $PGI_{\text{SES}}$ to the extent that this assumption holds.

To obtain $PGI_{SES}^{(1)}$ and $PGI_{SES}^{(2)}$, we randomly split the UKB GWAS sample into two subsamples ($N$=105,517~170,945) such that each subsample has the same male-female ratio and no individuals in one subsample are related to anyone in the other subsample with more than the third degree of relatedness. With each subsample, GWAS was run for the five numerical SES measures and the results were combined with Genomic SEM as described in Section 3.4. Then, $PGI_{SES}^{(1)}$ and $PGI_{SES}^{(2)}$ were constructed from one of the two independent GWAS subsample results in the main imaging sample.

Using $PGI_{SES}^{(1)}$ and $PGI_{SES}^{(2)}$, we fitted the model (4) by the GIV estimation, which is two-stage least squares (TSLS).

$$(8)\quad GMV_i^j = \tilde{\beta}_1^j PC1_{SES,i} + \tilde{\beta}_2^j PC2_{SES,i} + \theta^j PGI_i^{(1)} + Z_i^T \tilde{\gamma}^j + \tilde{\varepsilon}_i^j$$

where $PGI_i^{(1)}$ is the PGI estimated from the first subsample. The first-stage equation can be written as:

$$(9)\quad PGI_i^{(1)} = \alpha_1 PGI_i^{(2)} + \alpha_2 PC1_{SES,i} + \alpha_3 PC2_{SES,i} + Z_i^T \eta + e_i$$

where the PGI estimated from the second subsample, $PGI_i^{(2)}$, is used as an instrument for $PGI_i^{(1)}$. We obtained the TSLS estimates by fitting the following equation:

$$(10)\qquad GMV_i^j = \tilde{\beta}_1^j PC1_{SES,i} + \tilde{\beta}_2^j PC2_{SES,i} + \theta^j \widehat{PGI}_i^{(1)} + Z_i^T \tilde{\gamma}^j + \tilde{\varepsilon}_i^{*j}$$

where $\widehat{PGI}_i^{(1)}$ is the fitted value from the equation (8). The statistical inference was then conducted but in the standard TSLS framework to test the association between the GMV and SES for each voxel conditional on $PGI_{SES}$ (*61*).

We computed Partial $R^2$'s based on adding or excluding $\widehat{PGI}_i^{(1)}$ in model (10) instead of the unadjusted PGI. Similarly, we measured the difference in SES-GMV association after controlling for the PGI by GIV as $[(R^2_{PC+Z} - R^2_Z) - (R^2_{PC+\widehat{PGI}^{(1)}+Z} - R^2_{\widehat{PGI}^{(1)}+Z})] \,/\, (R^2_{PC+Z} - R^2_Z)$

#### 4.4. Functional annotations

We connected our anatomical findings to known functional localizations by leveraging Cognitive Atlas and the extrapolatable meta-analysis tool NeuroQuery (*36*, *37*). We first took the 518 cognitive concepts from Cognitive Atlas which were categorized into 11 functional categories.[12]

[12] These concepts were taken from https://www.cognitiveatlas.org/concepts/categories/all on 2 July 2021.

Then, for each concept, we generated a meta-analyzed Z-score brain map using NeuroQuery. This toolbox allows users to generate a predictive MRI-derived spatial distribution for any term, based on very large-scale meta-analyses containing mostly functional MRI studies. We excluded 12 concepts containing a term for which NeuroQuery failed to generate a brain map. As a result, 506 concepts remained. For each concept-associated brain-map, we calculated mean $\chi^2$ with voxels statistically significant at the FWE rate of 5% level in the VBM results. We used these mean $\chi^2$ scores as a summary measure of associations between a given functional concept and the regions linked to SES. The full results are reported in Table S17.

### 5. Interpretation

#### 5.1. Brain, SES, and genetics

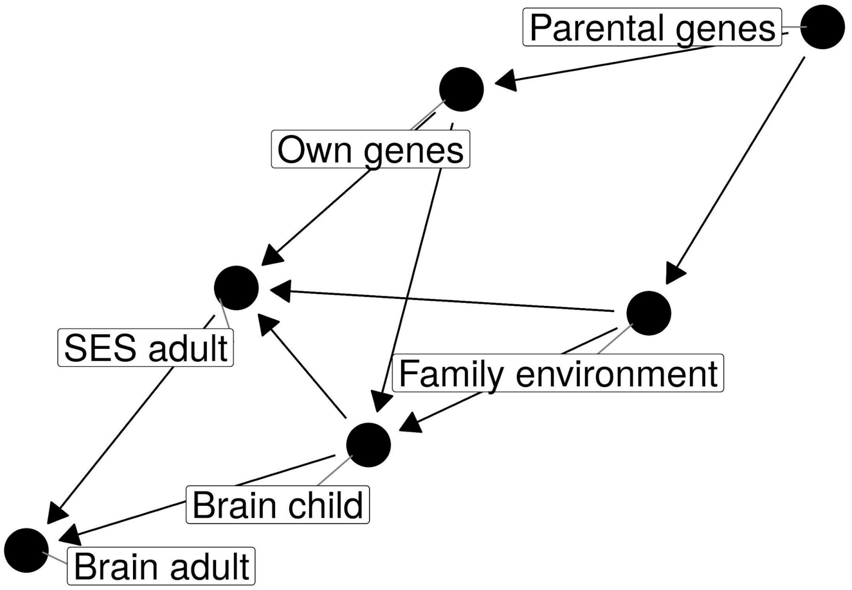


To aid interpretation of the association between SES and brain anatomy observed in late adulthood, the figure above describes a simple model that illustrates how adulthood brain anatomy can be linked to SES, family environments, and genetics. The model is depicted in a directed acyclic graph (DAG), a popular graphical framework for identifying confounding variables (*62–64*). The model does not attempt to include all possibly relevant factors and mediating pathways. Rather, its purpose is to identify what effects are potentially captured in the estimated GMV-SES association in relation to genetics and family environments.

$$[(R^2_{PC+P\hat{G}I^{(1)}+Z} - R^2_{P\hat{G}I^{(1)}+Z}) - (R^2_{PC+P\hat{G}I^{(1)}+BMI+Z} - R^2_{P\hat{G}I^{(1)}+BMI+Z})] \; / \; (R^2_{PC+P\hat{G}I^{(1)}+Z} - R^2_{P\hat{G}I^{(1)}+Z})$$

##### 6.3. Heterogeneity

Sex and age are two important factors for both SES and neuroanatomy. Therefore, we tested whether the SES-GMV associations are heterogeneous with respect to *i)* different sex (sex interaction) and *ii)* different ages (age interaction). We examined each aspect of heterogeneity separately by using the voxel clusters and including the interaction terms with $PC1_{SES}$ and $PC2_{SES}$. The interaction terms were then tested jointly with $F$-tests.

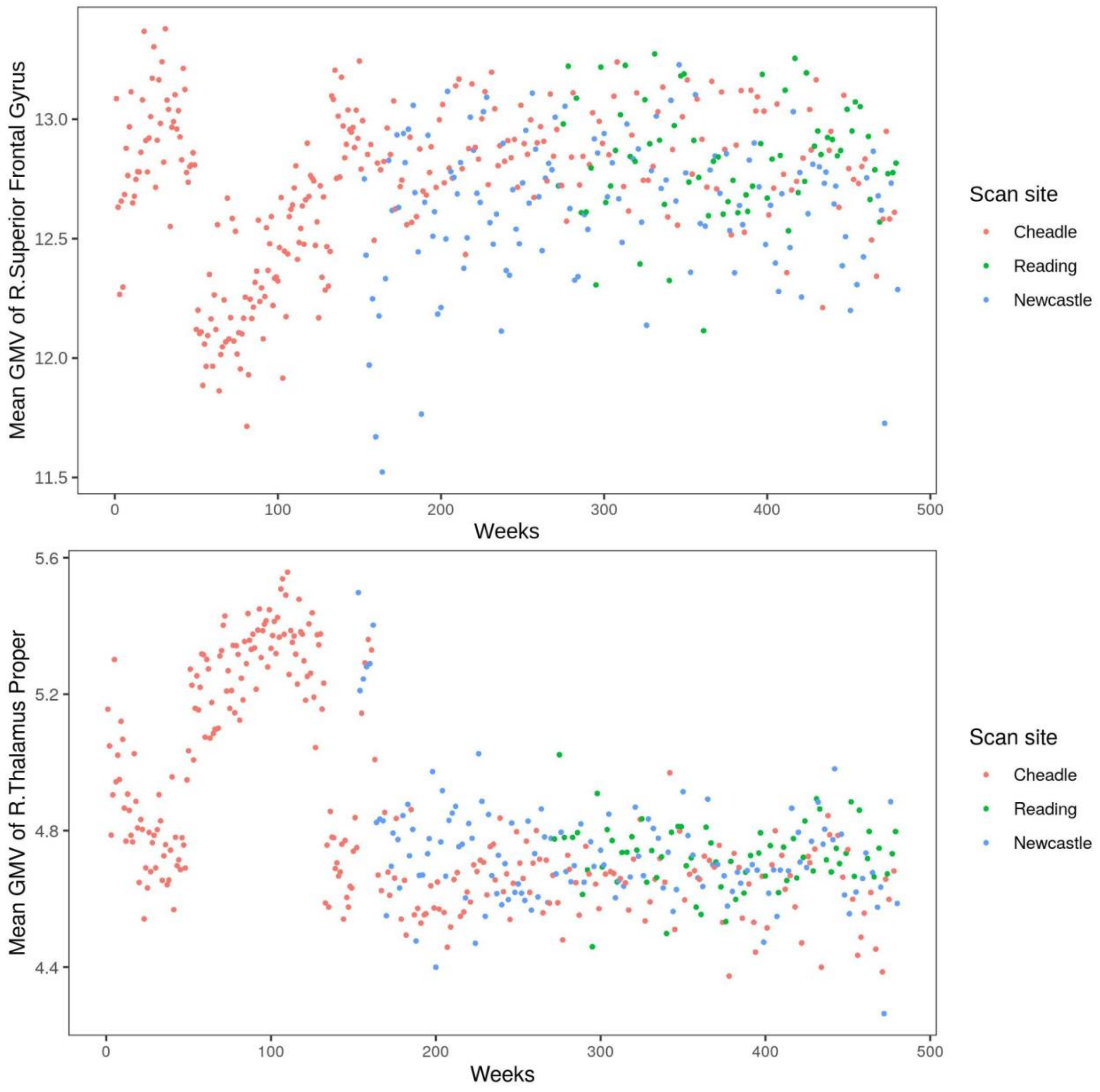


**Fig. S1. Average grey matter volume over acquisition dates by week**

Site-specific weekly averages of grey matter volume are plotted over acquisition dates for the right superior frontal gyrus and the right thalamus. These regions are selected as examples and there are other regions showing similar patterns.

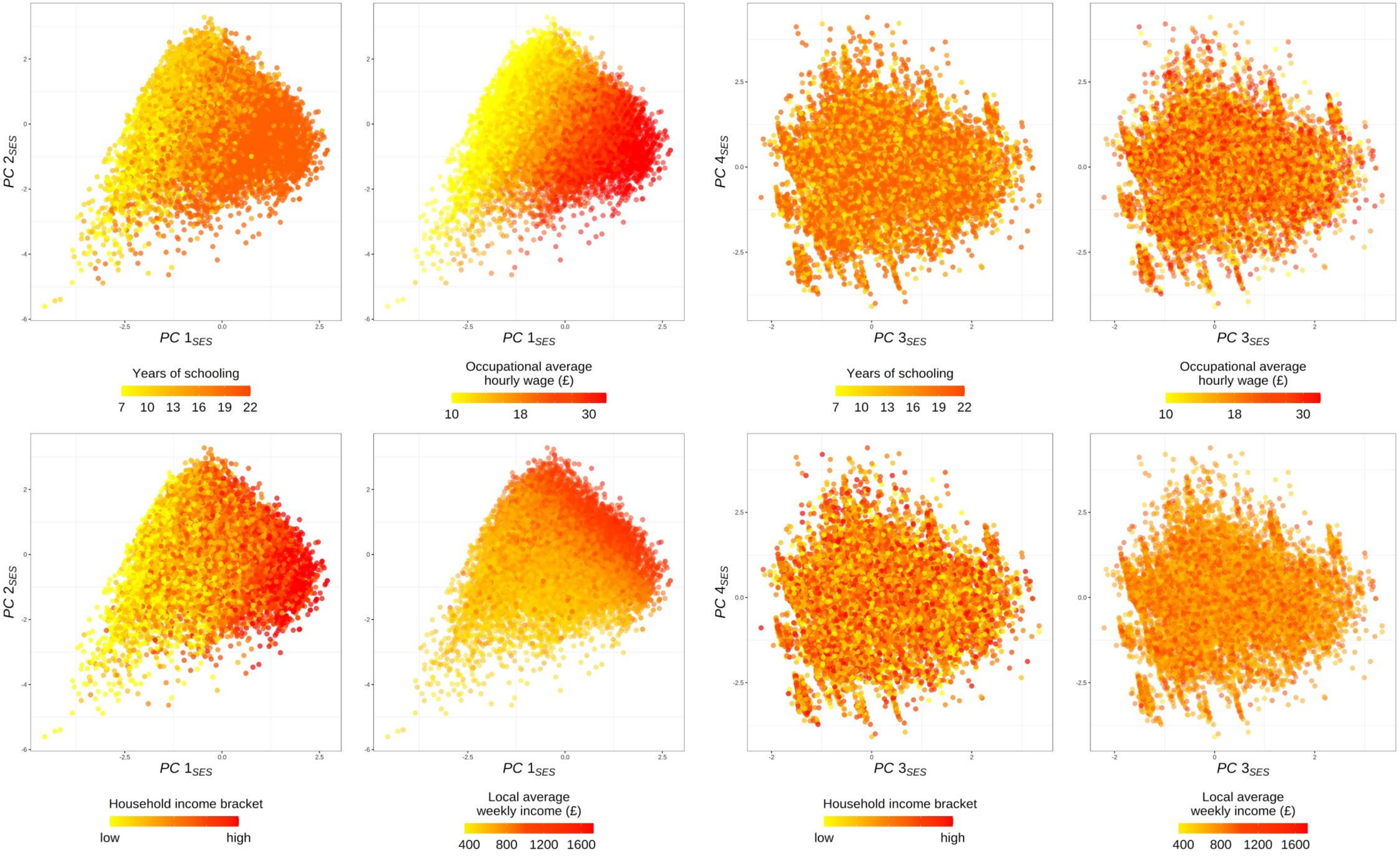


**Fig. S2. Scatter plots of top principal components (PC) for socioeconomic status (SES)**

Top four PCs for SES are plotted with color indicating different levels of selected socioeconomic measures.

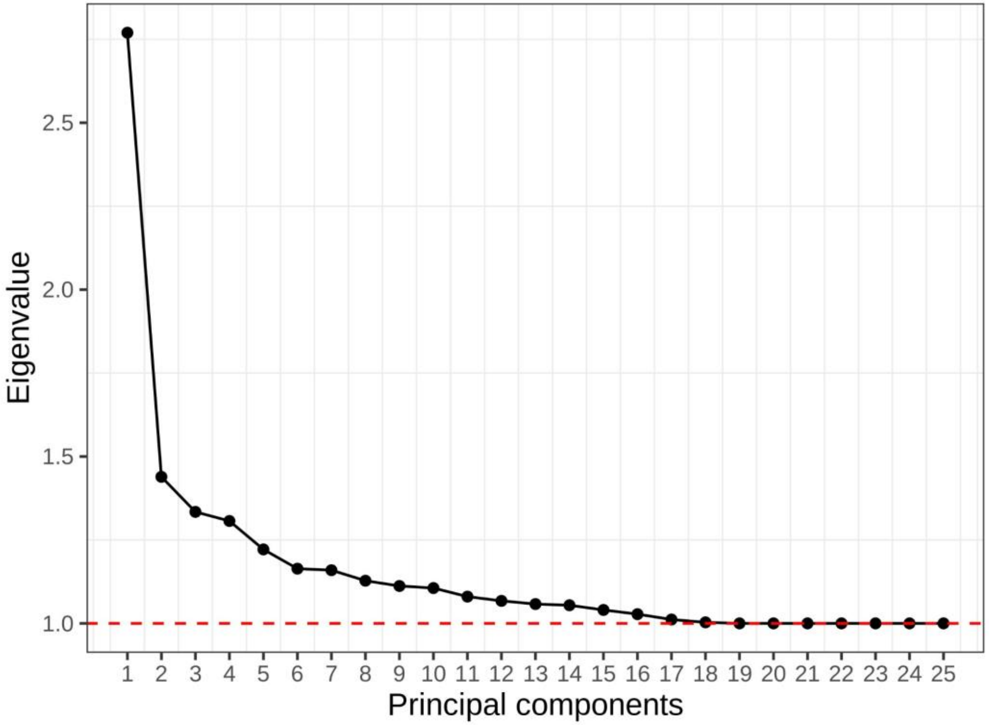


**Fig. S3. Eigenvalues from the principal component analysis of socioeconomic indicators**

**A.**

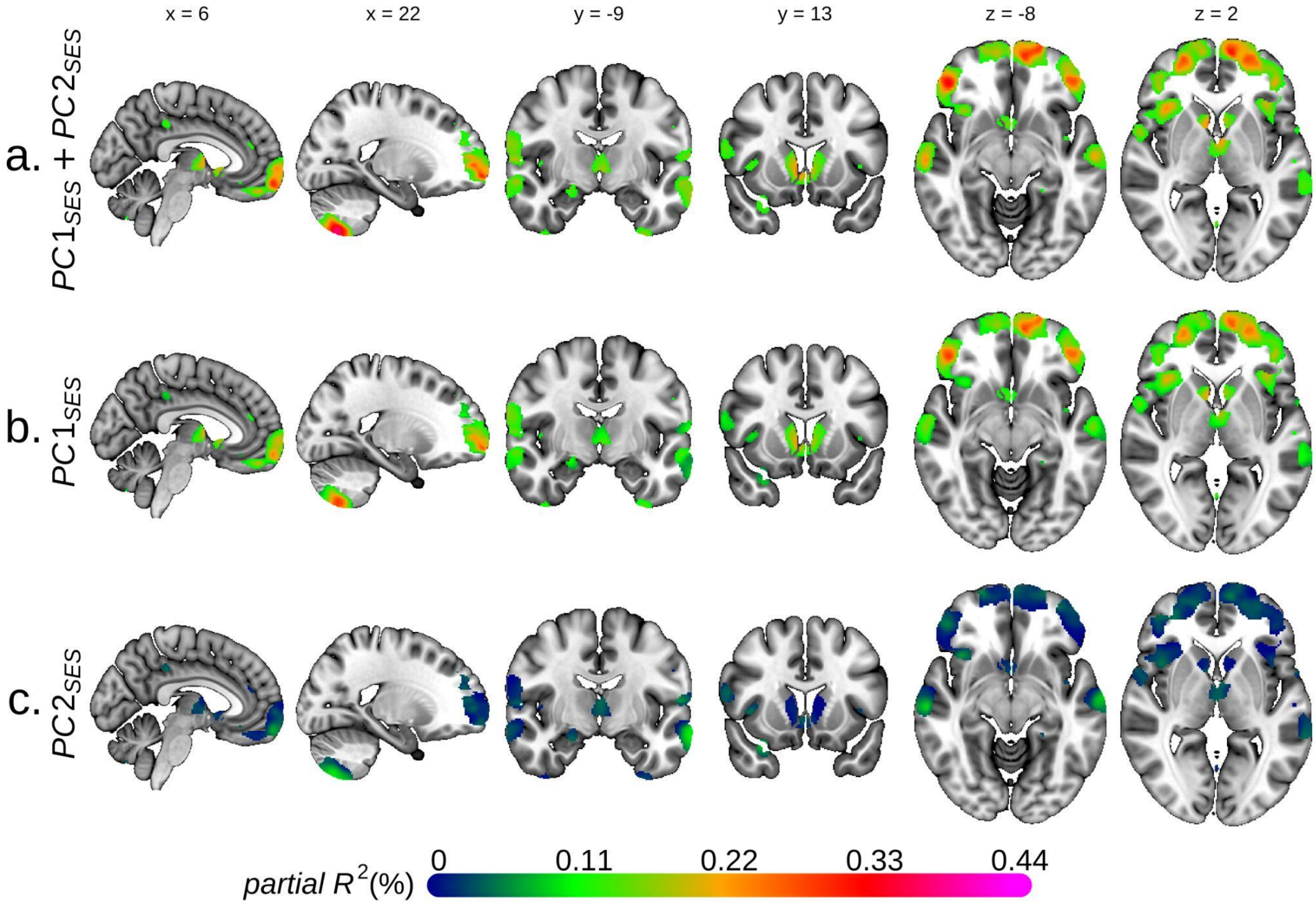

x = 6
x = 22
y = -9
y = 13
z = -8
z = 2
a. $PC1_{SES} + PC2_{SES}$
b. $PC1_{SES}$
c. $PC2_{SES}$
partial $R^2$(%)
0
0.11
0.22
0.33
0.44

**B.**

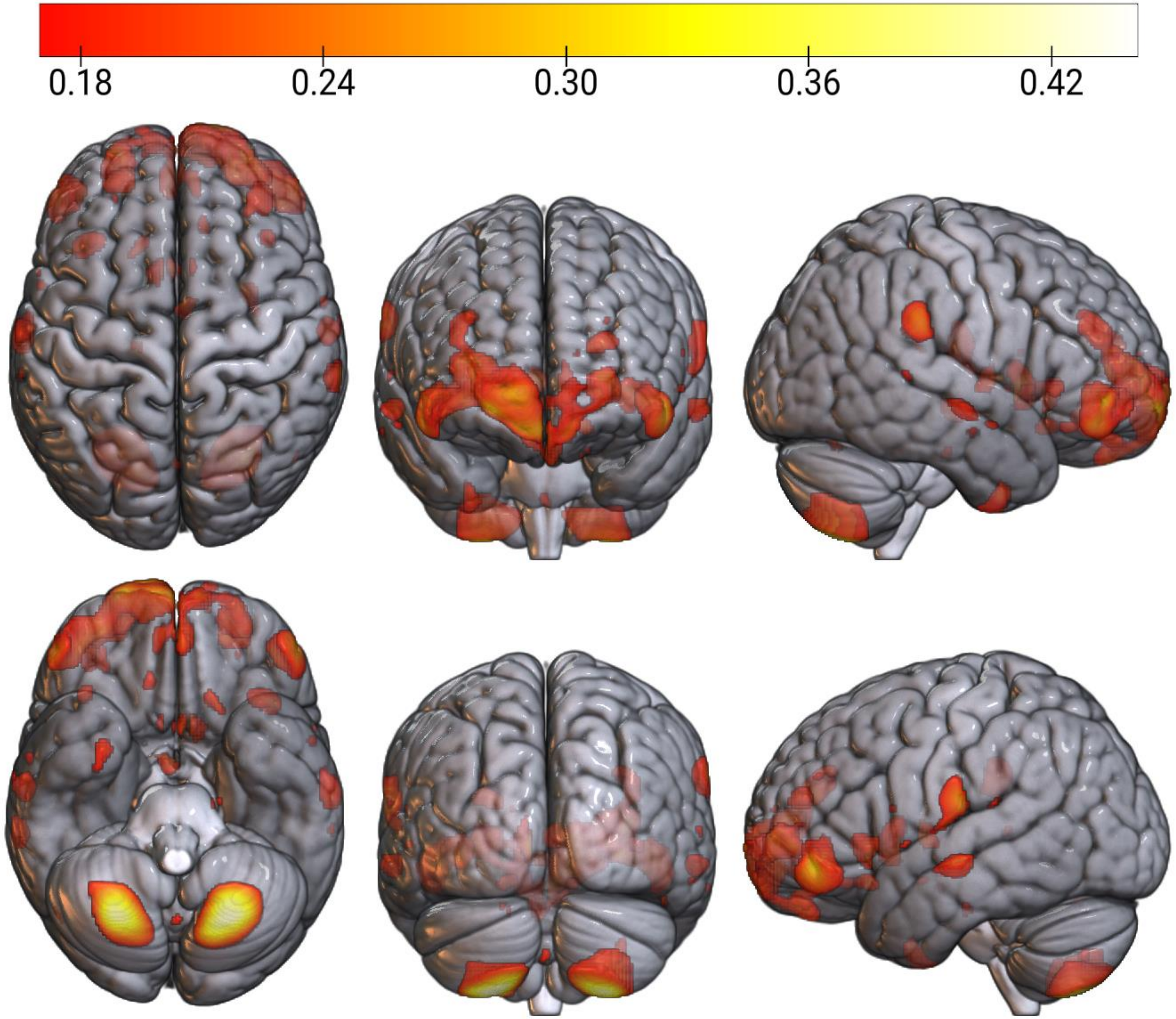


**Fig. S4. Voxel-based morphometry of grey matter volume and socioeconomic status (SES)**

Univariate voxel-based morphometry results on the two principal components (PC) for SES.

**A.** Each figure plots the association sizes measured in partial $R^2$ for only the voxels significant at FWE rate of 5%. MNI coordinates are indicated.

**B.** Partial $R^2$ (%) for two SES PC is plotted for only the voxels that were significant at FWE rate of 5% and had partial $R^2$>0.17%.

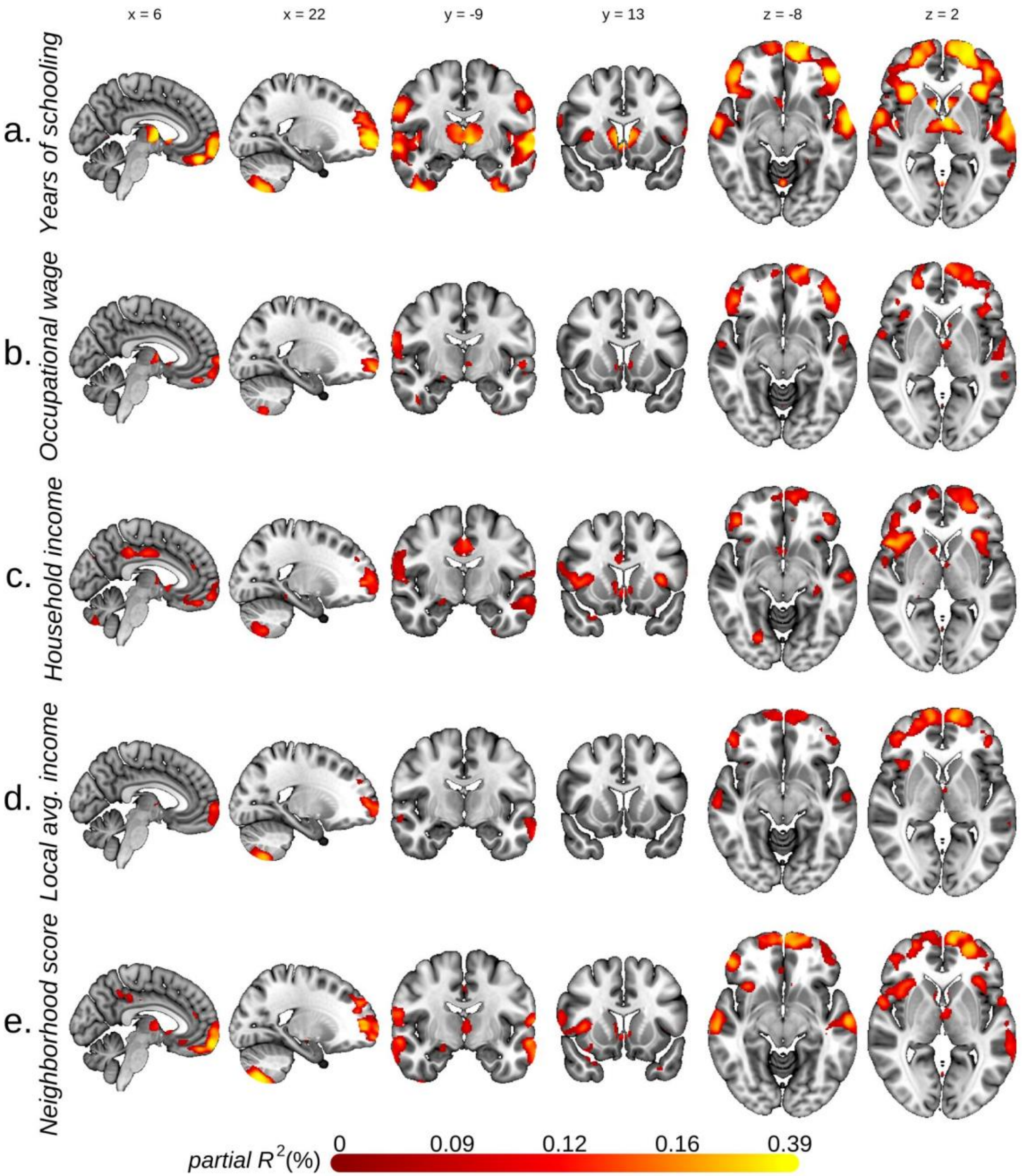


**Fig. S5. Voxel-based morphometry of grey matter volume and various socioeconomic measures**

Univariate voxel-based morphometry results, with grey matter volume as the dependent variable and each of the five socioeconomic measures as the explanatory variable. Each figure plots the association sizes measured in partial $R^2$ for only the voxels significant at FWE rate of 5%. MNI coordinates are indicated. $N$ = 30,954 (a), 27,307 (b), 29,839 (c), 29,006 (d), 28,405 (e)

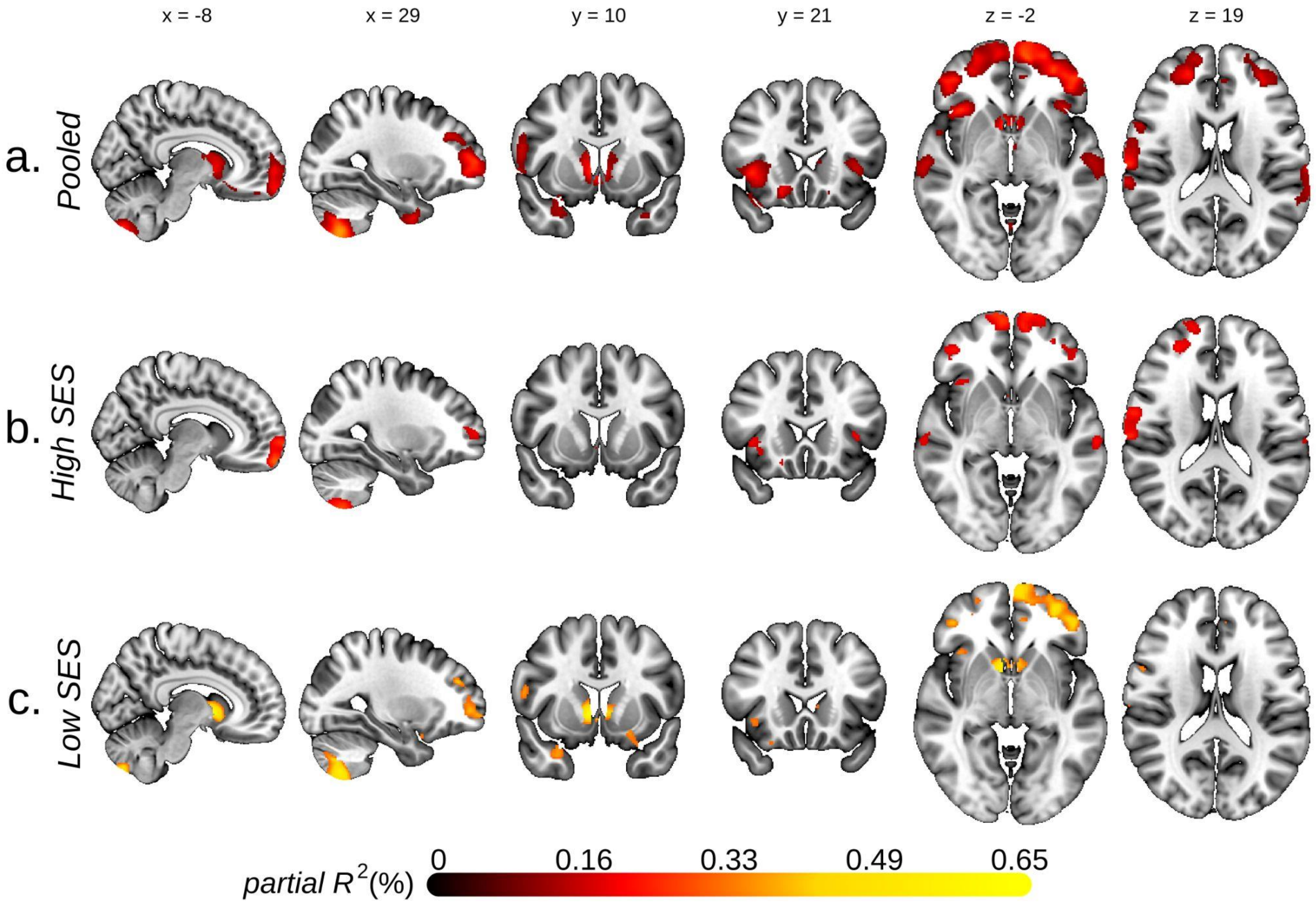


**Fig. S6. Stratified analyses on low and high socioeconomic status groups**

Results from baseline voxel-based morphometry analysis conducted separately on low and high socioeconomic status (SES) groups as well as the pooled sample. Each figure plots the association sizes measured in partial $R^2$ for only the voxels significant at FWE rate of 5%. High and low SES groups were defined by National Statistics Socio-economic Classification (high if holding a managerial, administrative, or professional occupation). MNI coordinates are indicated.

**A.**

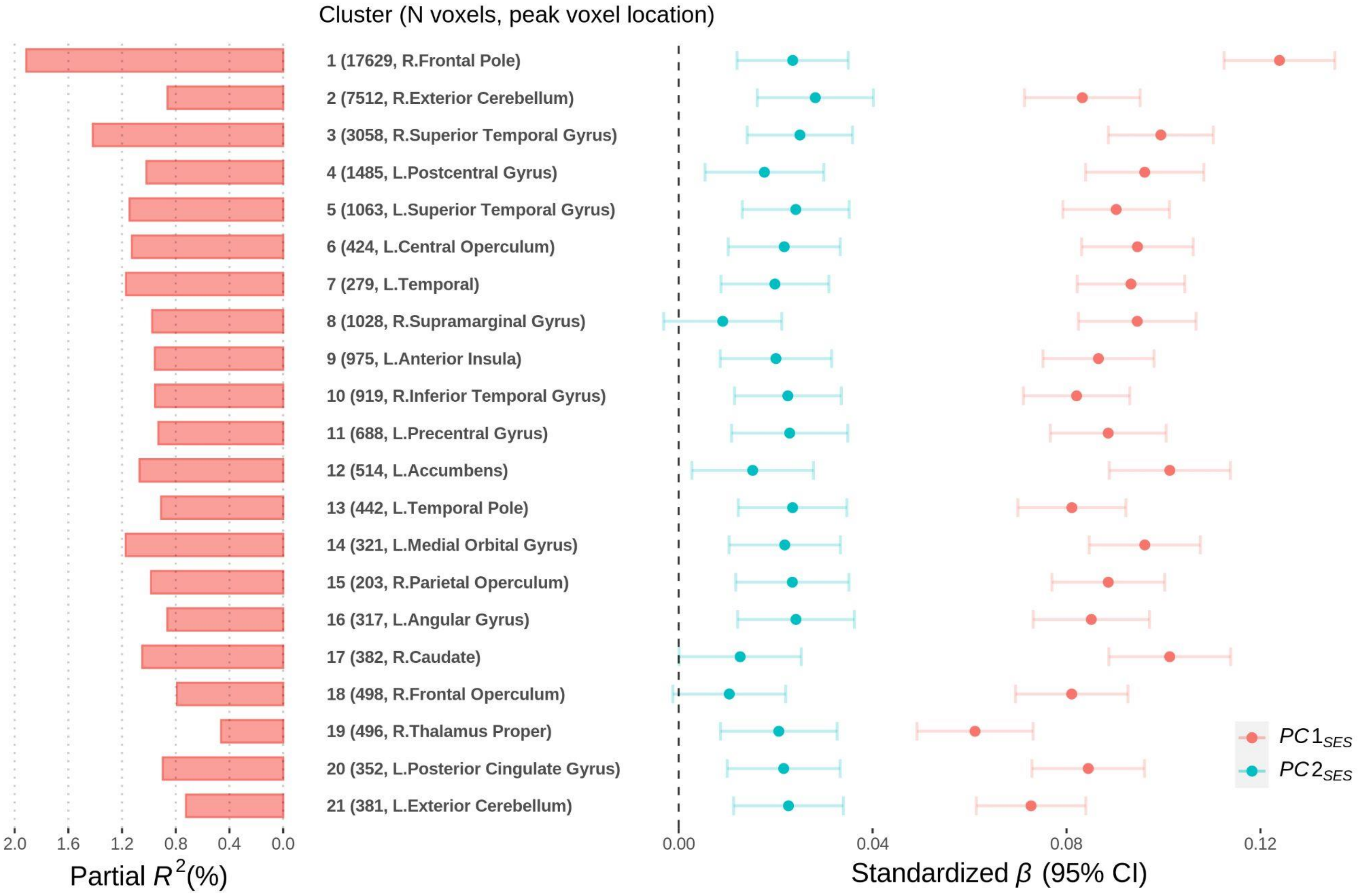

Cluster (N voxels, peak voxel location)
1 (17629, R.Frontal Pole)
2 (7512, R.Exterior Cerebellum)
3 (3058, R.Superior Temporal Gyrus)
4 (1485, L.Postcentral Gyrus)
5 (1063, L.Superior Temporal Gyrus)
6 (424, L.Central Operculum)
7 (279, L.Temporal)
8 (1028, R.Supramarginal Gyrus)
9 (975, L.Anterior Insula)
10 (919, R.Inferior Temporal Gyrus)
11 (688, L.Precentral Gyrus)
12 (514, L.Accumbens)
13 (442, L.Temporal Pole)
14 (321, L.Medial Orbital Gyrus)
15 (203, R.Parietal Operculum)
16 (317, L.Angular Gyrus)
17 (382, R.Caudate)
18 (498, R.Frontal Operculum)
19 (496, R.Thalamus Proper)
20 (352, L.Posterior Cingulate Gyrus)
21 (381, L.Exterior Cerebellum)
2.0 1.6 1.2 0.8 0.4 0.0
Partial $R^2$(%)
0.00 0.04 0.08 0.12
Standardized $\beta$ (95% CI)
$PC1_{SES}$
$PC2_{SES}$

**B.**

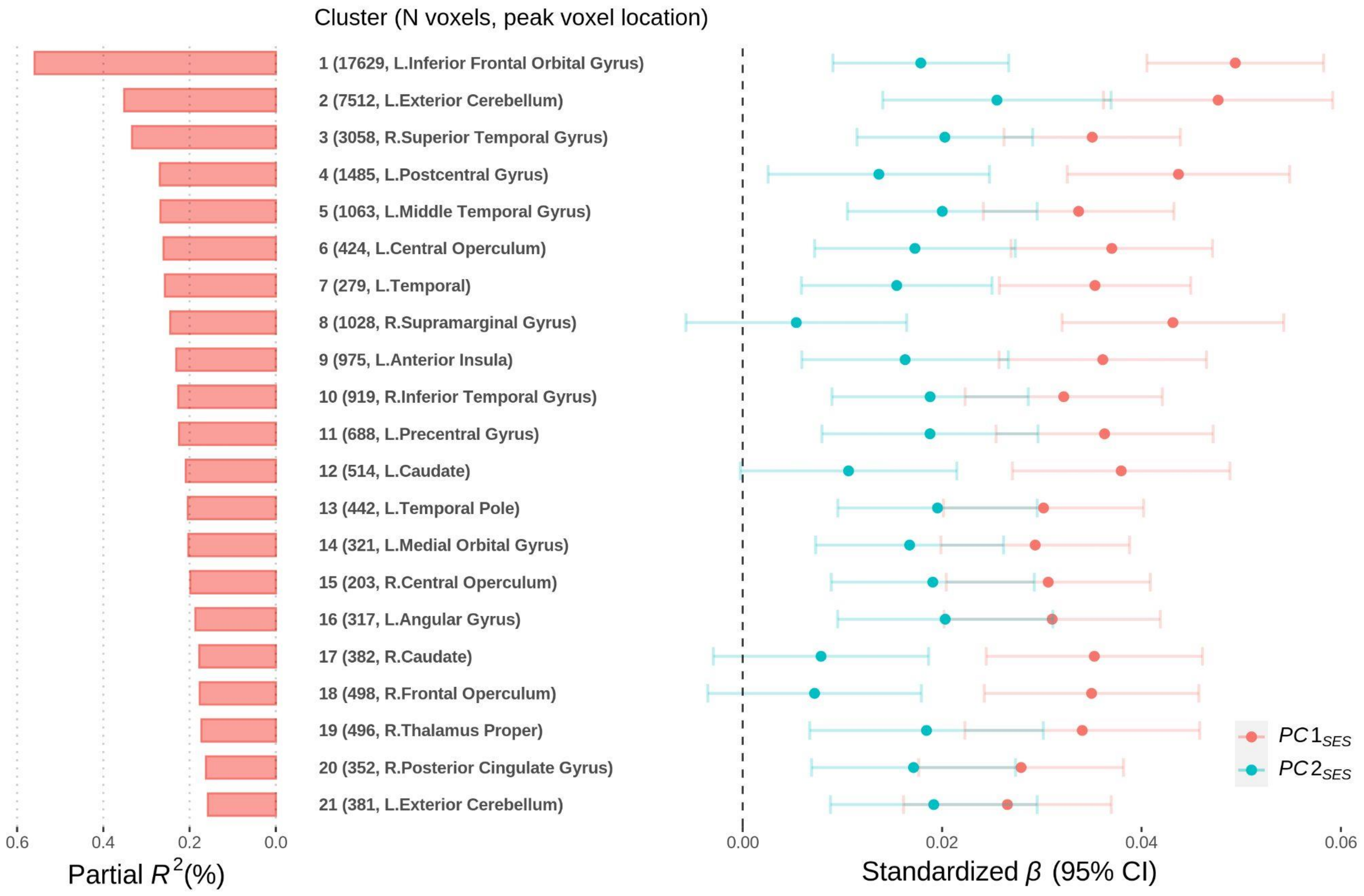


**Fig. S7. Standardized effect sizes of associations between socioeconomic status (SES) and grey matter volume (GMV) in voxel clusters**

Results from regressing GMV in each cluster on $PC1_{SES}$ and $PC2_{SES}$. Total intracranial volume was not controlled for in **A.** but in **B.** On the left, partial $R^2$ from both $PC1_{SES}$ and $PC2_{SES}$ are reported and, on the right, the standardized coefficient estimates are plotted with their uncorrected 95% confidence intervals. The clusters were formed with at least 200 voxels showing significant associations at FWE rate of 5% level in the baseline TIV-adjustd voxel-based morphometry (VBM) results on $PC1_{SES}$ and $PC2_{SES}$. For each cluster, the anatomical location of the peak voxel from the VBM results is indicated. See Table S8 for more information about the clusters.

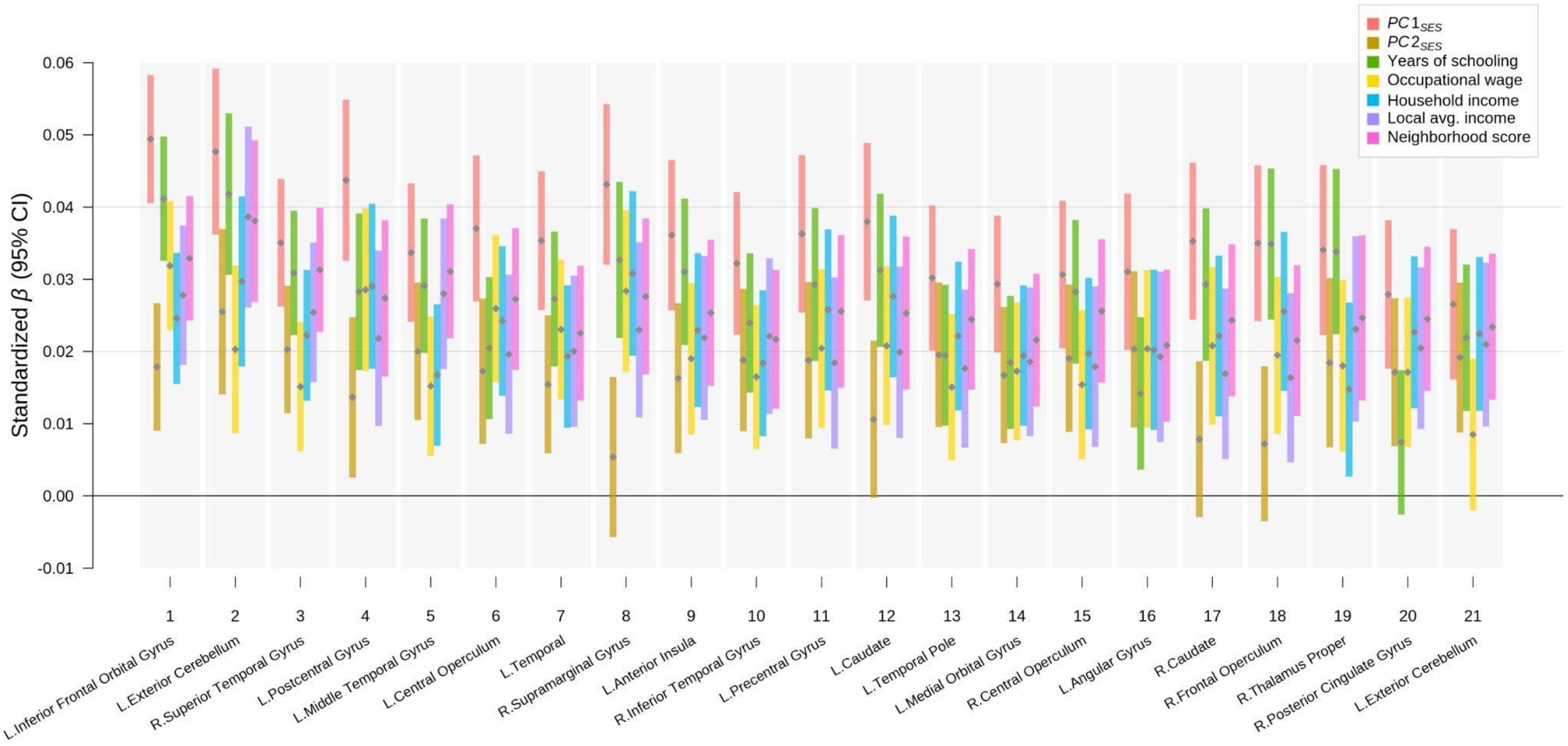


**Fig. S8. Standardized effect sizes of associations between various socioeconomic measures and grey matter volume (GMV) in voxel clusters**

Results from regressing GMV in each cluster on each of the socioeconomic measures separately. The standardized coefficient estimates (grey points) are plotted with their uncorrected 95% confidence intervals (color bars). The clusters were formed with at least 200 voxels showing significant associations at FWE rate of 5% level in the baseline voxel-based morphometry (VBM) results on $PC1_{SES}$ and $PC2_{SES}$. The clusters are ordered by the strength of joint associations with $PC1_{SES}$ and $PC2_{SES}$. For each cluster, the anatomical location of the peak voxel from the VBM results is indicated at the bottom. See Table S8 for more information about the clusters.

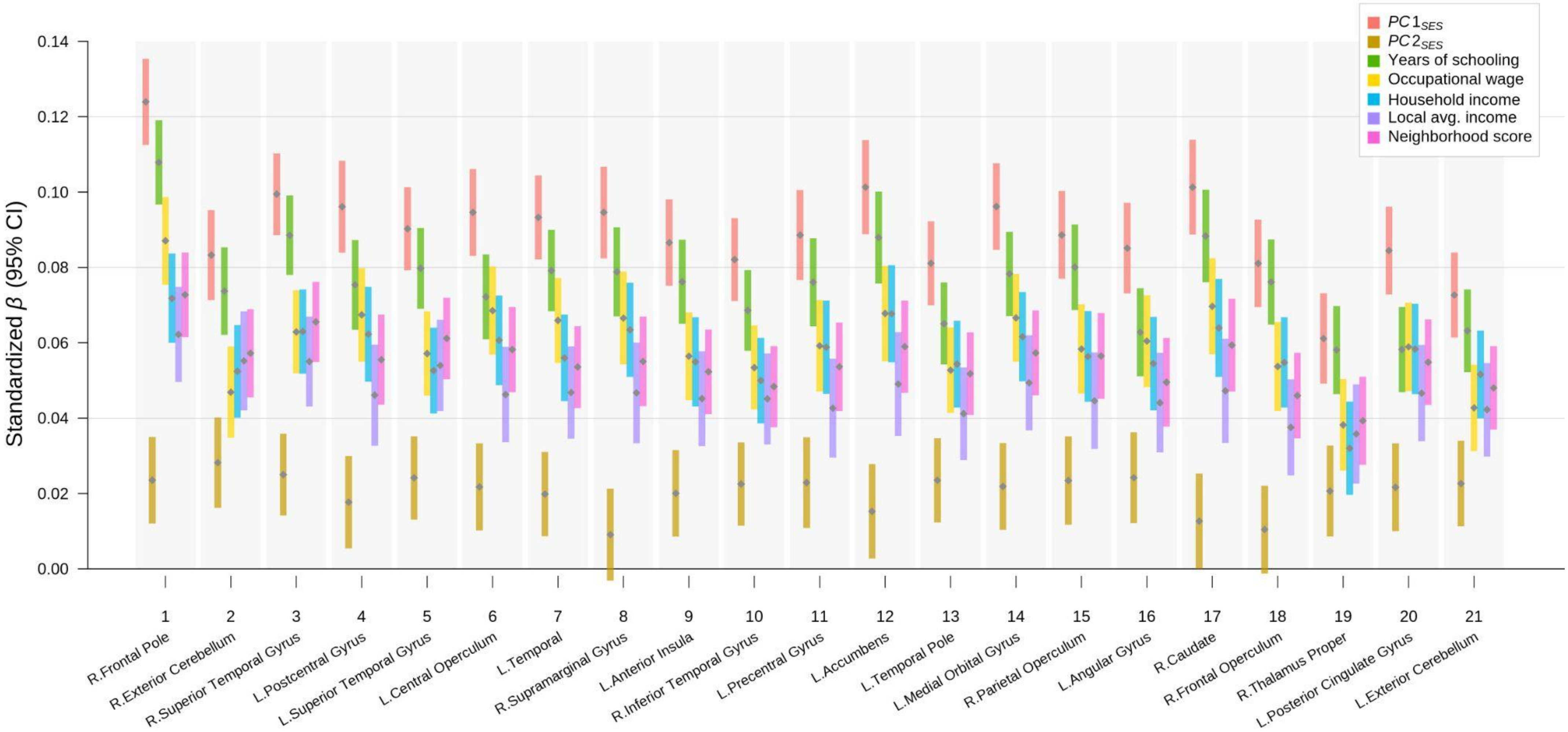


**Fig. S9. Standardized effect sizes of TIV-unadjusted associations between various socioeconomic measures and grey matter volume (GMV) in voxel clusters**

Results from regressing GMV in each cluster on each of the socioeconomic measures separately while not controlling for total intracranial volume (TIV). The standardized coefficient estimates (grey points) are plotted with their uncorrected 95% confidence intervals (color bars). The clusters were formed with at least 200 voxels showing significant associations at FWE rate of 5% level in the baseline voxel-based morphometry (VBM) results on $PC1_{SES}$ and $PC2_{SES}$. The clusters are ordered by the strength of joint associations with $PC1_{SES}$ and $PC2_{SES}$. For each cluster, the anatomical location of the peak voxel from the VBM results is indicated at the bottom. See Table S8 for more information about the clusters.

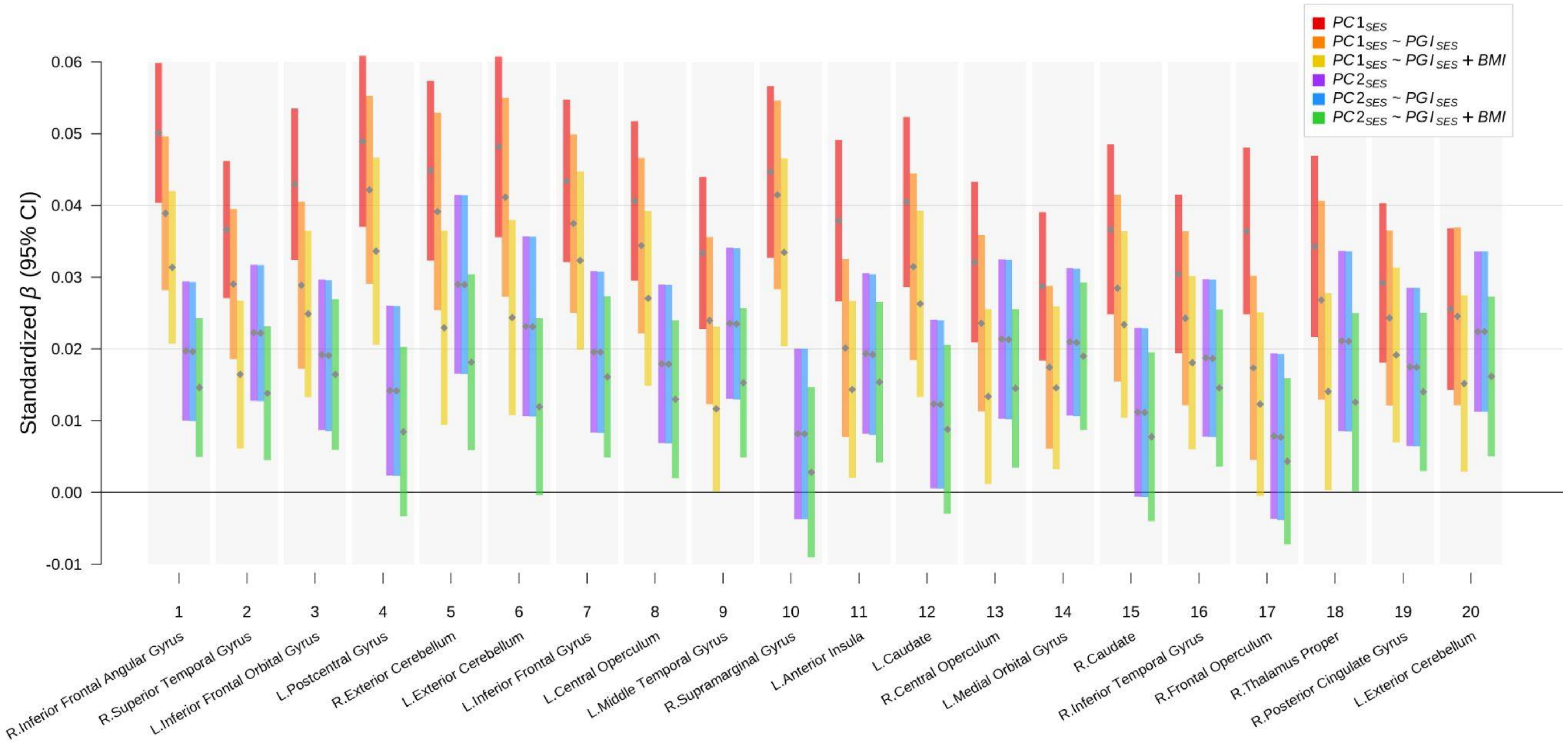


**Fig. S10. Standardized effect sizes of associations between socioeconomic status (SES) and grey matter volume (GMV) in voxel clusters with additional controls**

Results from regressing GMV in each cluster on $PC1_{SES}$ and $PC2_{SES}$ while additionally controlling for $PGI_{SES}$ and BMI. The standardized coefficient estimates (grey points) are plotted with their uncorrected 95% confidence intervals (color bars). The sample was restricted to individuals of European ancestry. Measurement error in $PGI_{SES}$ is adjusted for with genetic instrument variable (GIV) regression. The clusters were formed with at least 200 voxels showing significant associations at FWE rate of 5% level in the baseline voxel-based morphometry (VBM) results on $PC1_{SES}$ and $PC2_{SES}$. The clusters are ordered by the strength of joint associations with $PC1_{SES}$ and $PC2_{SES}$. For each cluster, the anatomical location of the peak voxel from the VBM results is indicated at the bottom. See Table S9 for more information about the clusters.

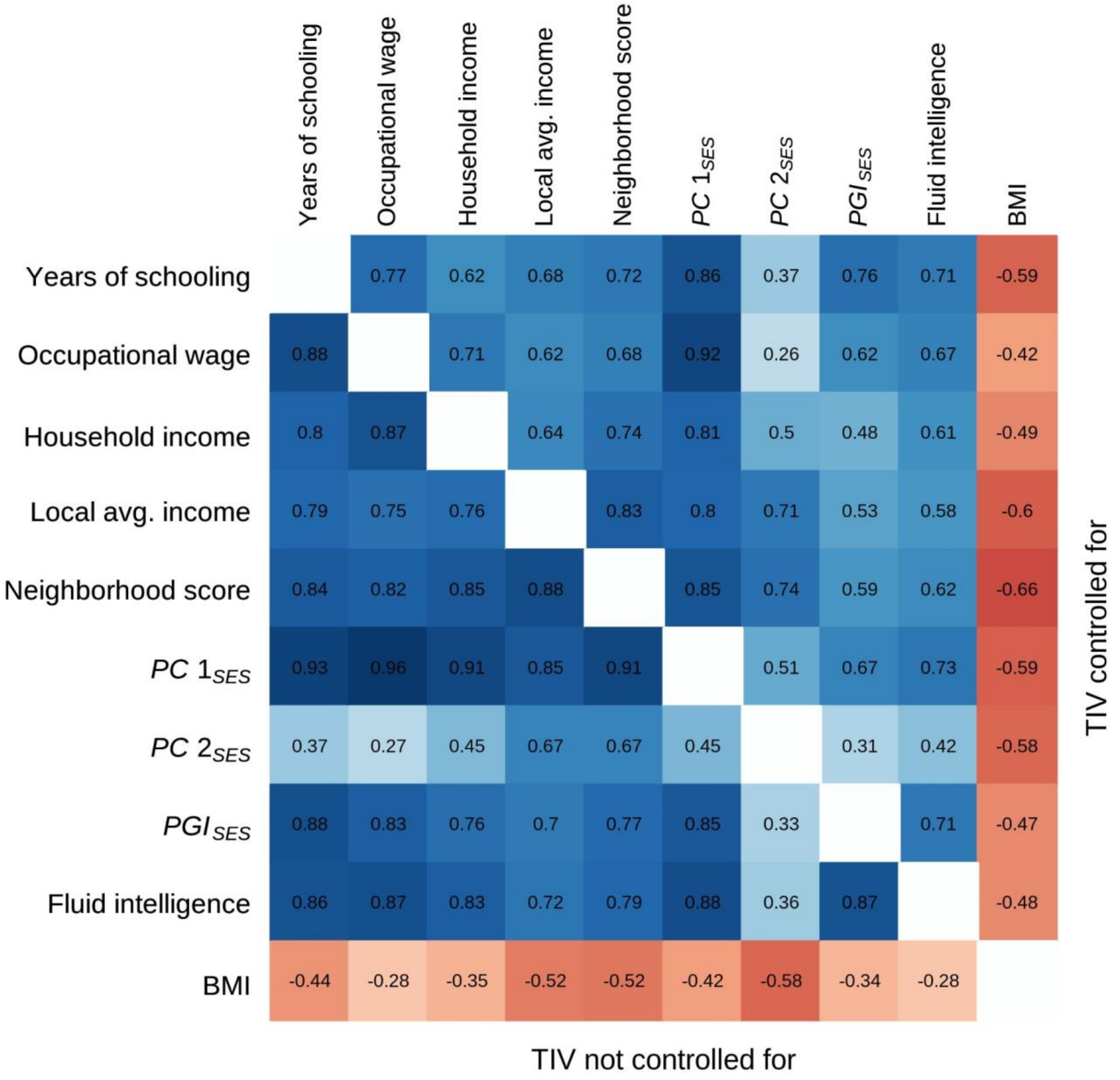


**Fig. S11. Grey-matter neuroanatomical correlations of various measures**

The figure plots pairwise Pearson correlations computed from *T*-statistics of univariate grey-matter VBM results for each measure. The upper triangle reports the correlations of the *T*-statistics from the VBM analyses that adjusted for total intracranial volume (TIV), while the lower triangle from those that did not adjust for TIV.

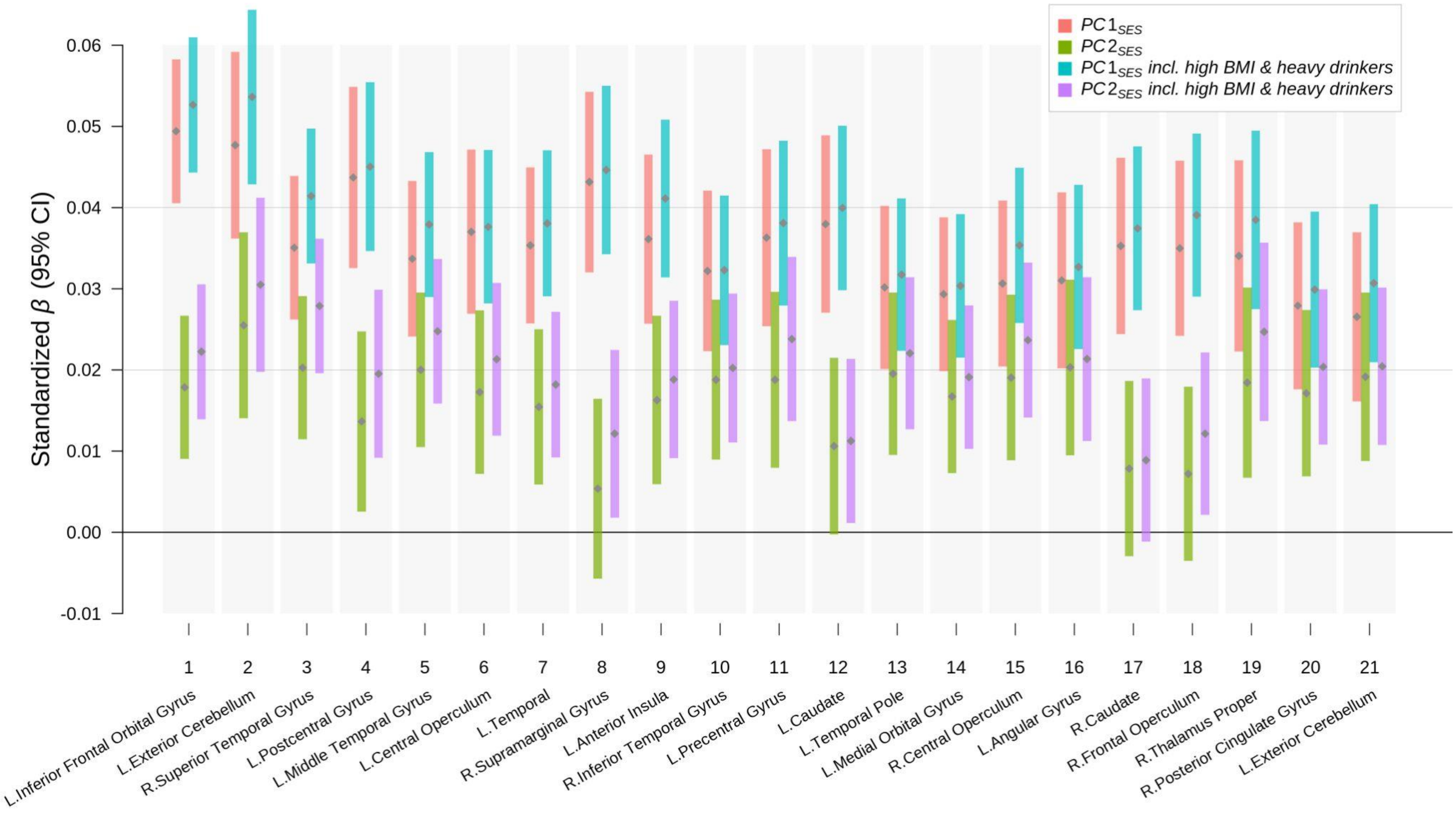


**Fig. S12. Standardized effect sizes of associations between socioeconomic status and grey matter volume (GMV) in voxel clusters with and without morbidly obese and heavy drinking individuals**

Results from regressing GMV in each cluster on $PC1_{SES}$ and $PC2_{SES}$ with and without morbidly obese and heavy drinking individuals. The standardized coefficient estimates (grey points) are plotted with their uncorrected 95% confidence intervals (color bars). The clusters were formed with at least 200 voxels showing significant associations at FWE rate of 5% level in the baseline voxel-based morphometry (VBM) results on $PC1_{SES}$ and $PC2_{SES}$. The clusters are ordered by the strength of joint associations with $PC1_{SES}$ and $PC2_{SES}$. For each cluster, the anatomical location of the peak voxel from the VBM results is indicated at the bottom. See Table S8 for more information about the clusters.

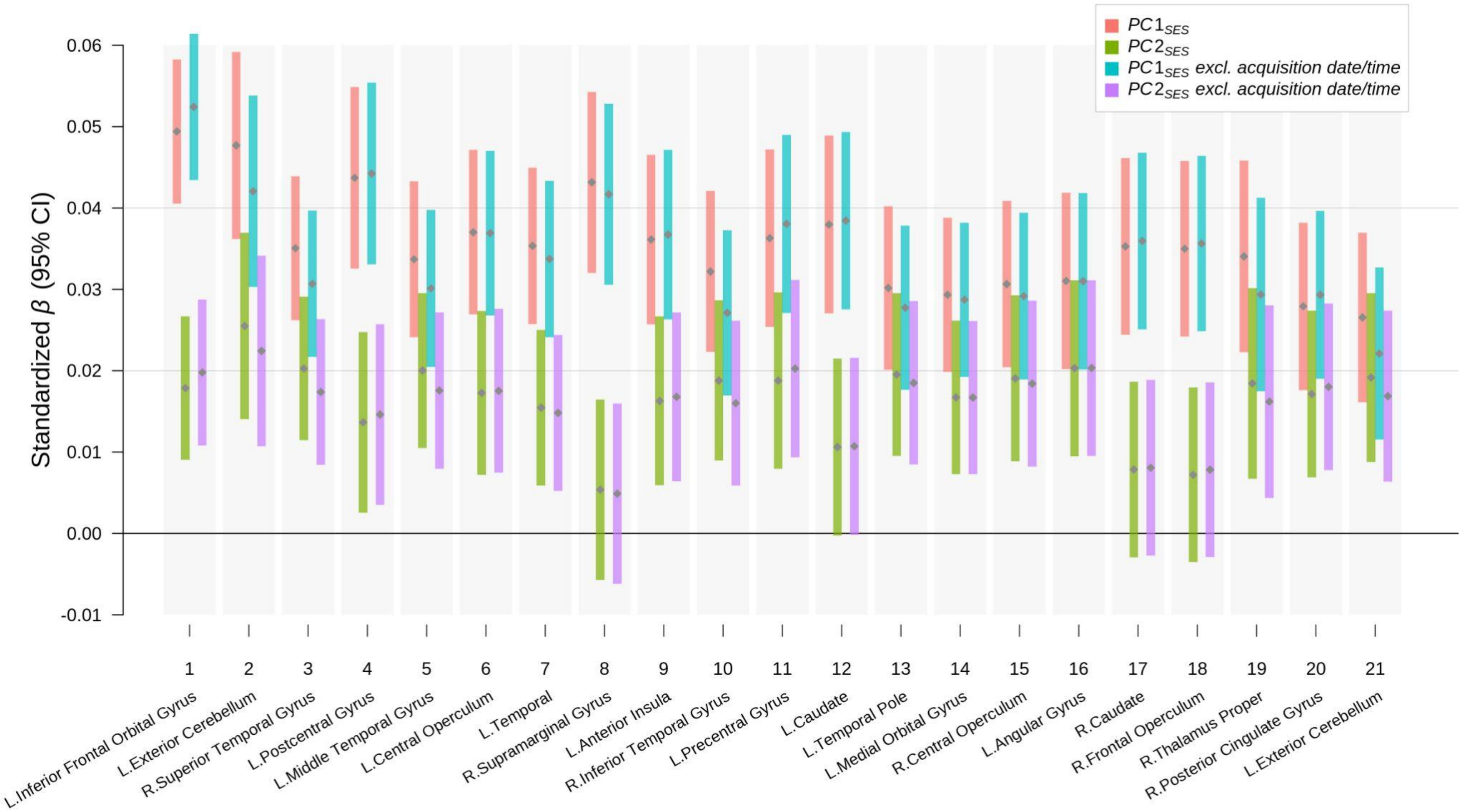


**Fig. S13. Standardized effect sizes of associations between socioeconomic status and grey matter volume (GMV) in voxel clusters with and without acquisition date and time as control variables**

Results from regressing GMV in each cluster on $PC1_{SES}$ and $PC2_{SES}$ with and without the acquisition date and time as control variables. The standardized coefficient estimates (grey points) are plotted with their uncorrected 95% confidence intervals (color bars). The clusters were formed with at least 200 voxels showing significant associations at FWE rate of 5% level in the baseline voxel-based morphometry (VBM) results on $PC1_{SES}$ and $PC2_{SES}$. The clusters are ordered by the strength of joint associations with $PC1_{SES}$ and $PC2_{SES}$. For each cluster, the anatomical location of the peak voxel from the VBM results is indicated at the bottom. See Table S8 for more information about the clusters.

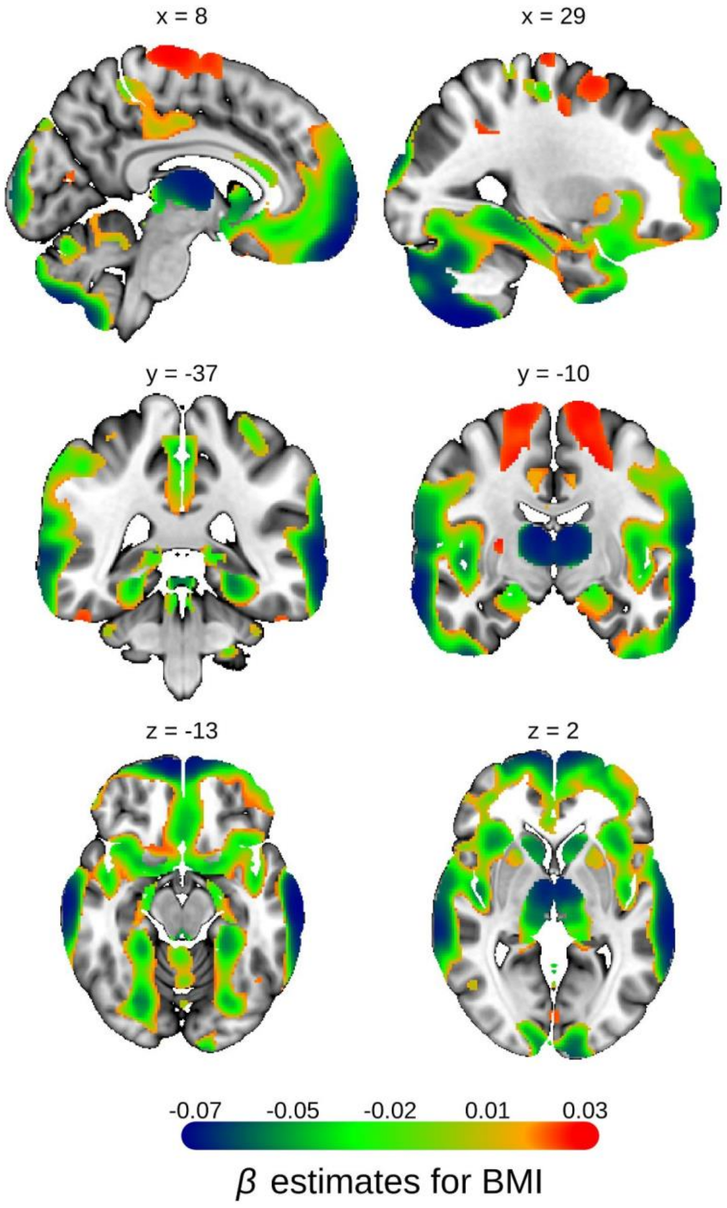


**Fig. S14. Voxel-based morphometry (VBM) of body mass index (BMI)**

Univariate VBM results on BMI, with grey matter volume (GMV) as the dependent variable. The Beta estimates are plotted for voxels significant at FWE rate of 1% level with partial $R^2$>0.02%. The Beta estimates are reported in the standard deviation unit of GMV. MNI coordinates are indicated.

#### Table S1 Descriptive statistics

| Variable | Stats / Values | Frequency | Graph | Valid |
|---|---|---|---|---|
| Year or birth<br>[integer] | Mean (sd) : 1954.4 (7.1)<br>min < med < max:<br>1937 < 1954 < 1970<br>IQR (CV) : 11 (0) | 34 distinct values | 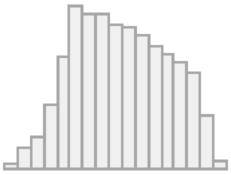 | 23931<br>(100.0%) |
| Year of assessment<br>[factor] | 1. 2014<br>2. 2015<br>3. 2016<br>4. 2017<br>5. 2018<br>6. 2019 | 1180 ( 4.9%)<br>3138 (13.1%)<br>3227 (13.5%)<br>4224 (17.7%)<br>6132 (25.6%)<br>6030 (25.2%) | 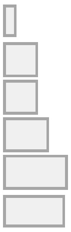 | 23931<br>(100.0%) |
| Age<br>[numeric] | Mean (sd) : 62.4 (7.2)<br>min < med < max:<br>44 < 63 < 81<br>IQR (CV) : 11 (0.1) | 38 distinct values | 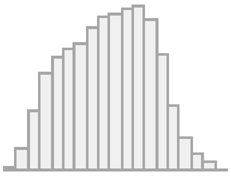 | 23931<br>(100.0%) |
| Sex<br>[factor] | 1. Female<br>2. Male | 12613 (52.7%)<br>11318 (47.3%) | 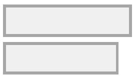 | 23931<br>(100.0%) |
| European ancestry<br>[factor] | 1. No<br>2. Yes | 3132 (13.1%)<br>20799 (86.9%) | 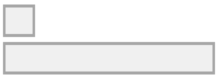 | 23931<br>(100.0%) |
| SES PC1<br>[numeric] | Mean (sd) : 0 (1)<br>min < med < max:<br>-4.6 < 0.1 < 2.7<br>IQR (CV) : 1.4 (-5.64027e+16) | 23876 distinct values | 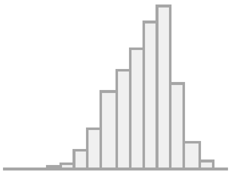 | 23931<br>(100.0%) |
| SES PC2<br>[numeric] | Mean (sd) : 0 (1)<br>min < med < max:<br>-5.6 < 0 < 3.3<br>IQR (CV) : 1.3 (-5.166366e+16) | 23876 distinct values | 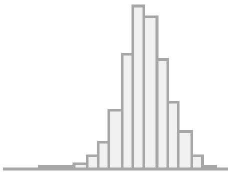 | 23931<br>(100.0%) |
| Occupation (SEC)<br>[factor] | 1. Higher managerial, administrative, professional<br>2. Lower managerial, administrative, professional<br>3. Intermediate<br>4. Routine and manual | 7272 (30.4%)<br>8339 (34.8%)<br>5022 (21.0%)<br>3298 (13.8%) | 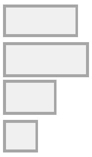 | 23931<br>(100.0%) |

| | | | | |
|---|---|---|---|---|
| Years of schooling<br>[numeric] | Mean (sd) : 16.4 (4.2)<br>min < med < max:<br>7 < 20 < 22<br>IQR (CV) : 7 (0.3) | 16 distinct values | 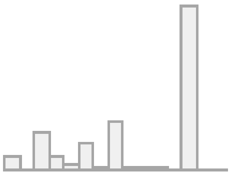 | 23867<br>(99.7%) |
| Occupational hourly wage<br>[numeric] | Mean (sd) : 19.6 (8.6)<br>min < med < max:<br>6.8 < 18.2 < 76.2<br>IQR (CV) : 11.4 (0.4) | 613 distinct values | 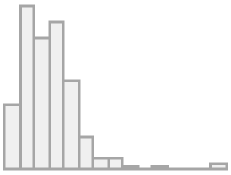 | 23931<br>(100.0%) |
| Household income<br>[factor] | 1. < 18K<br>2. 18K to 30,999<br>3. 31K to 51,999<br>4. 52K to 100K<br>5. > 100K | 2760 (11.5%)<br>6330 (26.5%)<br>7302 (30.5%)<br>5762 (24.1%)<br>1777 ( 7.4%) | 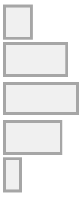 | 23931<br>(100.0%) |
| Local average income<br>[numeric] | Mean (sd) : 756.3 (185.8)<br>min < med < max:<br>350 < 730 < 1730<br>IQR (CV) : 230 (0.2) | 111 distinct values | 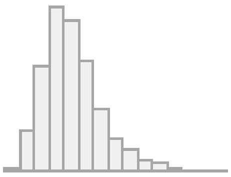 | 23931<br>(100.0%) |
| Housing<br>[factor] | 1. Own outright<br>2. Own with a mortgage<br>3. Rent, public<br>4. Rent, private<br>5. Pay part rent and part mortgage<br>6. Free rent | 17854 (74.6%)<br>5094 (21.3%)<br>343 ( 1.4%)<br>482 ( 2.0%)<br>60 ( 0.3%)<br>98 ( 0.4%) | 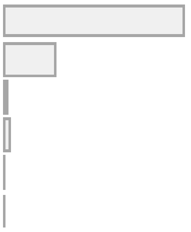 | 23931<br>(100.0%) |
| Alcohol intake<br>[numeric] | Mean (sd) : 7 (5.9)<br>min < med < max:<br>0 < 6 < 24<br>IQR (CV) : 9 (0.8) | 25 distinct values | 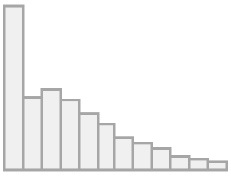 | 23931<br>(100.0%) |
| BMI<br>[numeric] | Mean (sd) : 25.9 (3.5)<br>min < med < max:<br>13.4 < 25.6 < 35<br>IQR (CV) : 5 (0.1) | 13485 distinct values | 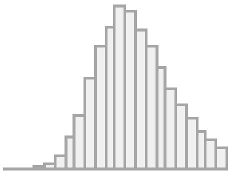 | 23931<br>(100.0%) |
| Scanning site<br>[factor] | 1. Cheadle<br>2. Reading<br>3. Newcastle | 16293 (68.1%)<br>3135 (13.1%)<br>4503 (18.8%) | 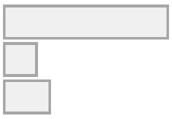 | 23931<br>(100.0%) |

| | | | | |
|---|---|---|---|---|
| Image quality rating [numeric] | Mean (sd) : 84.8 (1.4)<br>min < med < max:<br>71 < 85.2 < 87<br>IQR (CV) : 1.3 (0) | 901 distinct values | 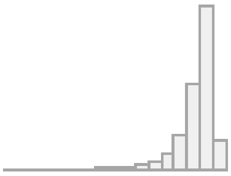 | 23931 (100.0%) |
| Total intracranial volume [numeric] | Mean (sd) : 1531.9 (145.1)<br>min < med < max:<br>993.3 < 1523.3 < 2286.9<br>IQR (CV) : 206.2 (0.1) | 23930 distinct values | 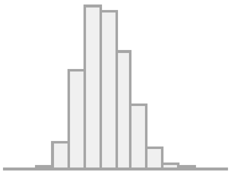 | 23931 (100.0%) |
| SES Polygenic index [numeric] | Mean (sd) : 0 (1)<br>min < med < max:<br>-4 < 0 < 4.4<br>IQR (CV) : 1.4 (2.060389e+17) | 20267 distinct values | 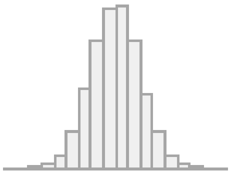 | 20799 (86.9%) |
